# REM sleep prefrontal high-frequency oscillation chains mediate distinct cortical – hippocampal reactivation patterns compared to NREM sleep

**DOI:** 10.1101/2025.09.15.676366

**Authors:** Justin D. Shin, Michael Satchell, Paul Miller, Shantanu P. Jadhav

## Abstract

REM (rapid eye movement) and non-REM (NREM) sleep stages contribute to systems memory consolidation in hippocampal-cortical circuits. However, the physiological mechanisms underlying REM memory processes remain relatively unclear compared to NREM memory reactivation. Here we report, in rodents, the existence of prefrontal cortical (PFC) high-frequency oscillation (HFO) chains in REM sleep during consolidation of recently acquired spatial memory. High-density tetrode recordings in hippocampal area CA1 and PFC reveal that REM cortical HFOs occur in characteristic chains that are phase modulated by theta oscillations, corresponding to increased CA1-PFC theta coherence and delineating periods of enhanced hippocampal-cortical communication. REM HFO chains sequentially organize sparse PFC ensemble reactivation of behavioral activity during periods of local suppression, distinct from widespread reactivation bursts during NREM ripple oscillations. REM HFO chains also preferentially engage CA1 neuronal populations that demonstrate a shift in their preferred theta-phase from behavior to REM sleep. CA1 neuronal activation during REM HFO chains was correlated with CA1 activity suppression during NREM PFC ripples, and linked to differential changes in CA1 firing rates in sleep, suggesting REM-driven regulation of hippocampal excitability. A cortical network model incorporating the effects of acetylcholine can reproduce the distinct REM and NREM activity patterns, providing a mechanistic basis for widespread coactivity during NREM cortical ripples, compared to sparse, temporally extended reactivation on a background of local suppression during REM HFO chains. Overall, these findings establish a role for PFC high-frequency oscillations in regulating distinct dual sleep stage reactivation patterns.

## INTRODUCTION

Rapid eye movement (REM) sleep, often referred to as paradoxical sleep due to its electrophysiological similarities to wakefulness, is a relatively transient stage of sleep that occurs after NREM sleep stages in cycles, and plays a crucial role in a wide range of brain functions.^1,2^ Although largely appreciated for its role in emotional processing, recent evidence has highlighted its role in the consolidation of spatial, declarative, and procedural memory, indicating a broader function of REM sleep in mnemonic processing.^3-5^ Importantly, REM dysregulation is associated with memory deficits and is thought to contribute to the progression of neuropsychiatric disorders and neurodegenerative diseases such as depression and dementia, highlighting the critical need for a deeper understanding of this sleep state to benefit human health.^6^

Primary models of systems memory consolidation, including the standard two-stage theory of consolidation or trace transformation theory,^7-10^ rest under the assumption of cooperative hippocampal-cortical reactivation during sleep to transform initially encoded episodic hippocampal representations into broadly distributed hippocampal-cortical representations for long-term storage and semantic memory formation. In support of this view, several rodent and human studies have confirmed that, in non-rapid eye movement (NREM) sleep, coordination of hippocampal sharp-wave ripples (SWRs) with cortical slow oscillations (1-4 Hz), spindles (12-16 Hz), and ripples (150-250 Hz) underlies the neurophysiological mechanism supporting consolidation.^11-14^ While the role of offline memory reactivation in the hippocampus and prefrontal cortex (PFC) during NREM sleep has been extensively studied in relation to memory function, the precise mechanisms through which REM sleep contributes to these processes, particularly in the context of hippocampal-cortical interactions, is still unclear.

REM sleep is characterized by high theta-delta ratio,^15,16^ and while hippocampal SWRs are primarily reported in NREM states with relatively low theta power, hippocampal theta rhythms during REM sleep have been shown to be important for memory consolidation.^17^ Correspondingly, phasic theta activity bursts during REM are often coupled with elevated gamma and cortical high-frequency oscillation (HFO) power in hippocampal and cortical networks, potentially organizing neural activity in distributed circuits for mnemonic processing.^18-25^ Furthermore, REM sleep is associated with a unique neuromodulatory landscape that is conducive for the induction of plasticity within and across circuits.^26^ However, the role of REM sleep cortical oscillations, whether they are coordinated with hippocampal activity patterns or with cortical-hippocampal reactivation, and the relationship between REM and NREM activity for dual sleep state regulation of reactivation and excitability are all open questions.

Cortical ripples (high-frequency or fast oscillations, ∼150-250 Hz) have been reported in NREM sleep,^27-29^ and our recent study showed that independent PFC ripples that are dissociated from hippocampal SWRs are prevalent in NREM sleep and mediate suppression of hippocampal activity and reactivation, potentially facilitating the selection and consolidation of specific hippocampal memory ensembles, while facilitating local cortical reactivation.^28^ A related study had similar findings, suggesting that these cortical ripples reflect intracortical processing that occurs independently of hippocampal input, allowing for interference-free consolidation of memory representations within cortical circuits.^30^

In REM sleep, previous studies have established cortical HFOs (in a similar frequency range to high-frequency NREM ripples) along with gamma oscillations – REM HFOs have been detected in different frequency ranges and sometimes reported with different nomenclature.^19,21,22,31^ Since recent studies note the possibility of spurious spectral detection in high frequency ranges,^32^ monitoring spiking activity simultaneously with LFP/EEG activity can therefore provide key confirmation about the existence and impact of high-frequency oscillations on neuronal activity via spiking modulation in local circuits.

Previous studies in rodents have reported a role of REM sleep in homeostatic regulation of firing rates, using recordings lasting from less than an hour to across the 24-hour cycle, but without the context of a behavioral task.^33-35^ Studies have also reported prominent phenomena such as hippocampal REM-theta phase reversing cells^36^ and hippocampal reactivation,^37^ using behavioral tasks and examining brief sleep periods in post-task rest sessions. Further, a recent study in mice reported PFC coding and activation of recent experiences and inferred knowledge in NREM and REM sleep using imaging methods, but did not examine any potential role of oscillations.^38^

Despite these results, an explicit examination of REM cortical HFO events, and how they impact cortical and hippocampal activity during consolidation of recent experiences is not known, despite the similarity of these high-frequency oscillations to cortical ripples in NREM. Furthermore, how NREM and REM sleep activity patterns may together complementarily shape the development and refinement of cortical-hippocampal ensembles to support learning is unclear. Therefore, we used high-density electrophysiological recordings in PFC and CA1 during learning and subsequent sleep to investigate whether and how coordinated activity during REM sleep may drive circuit changes underlying systems memory consolidation, and whether prefrontal and hippocampal dynamics differ during cortical ripple and HFO activity patterns in NREM and REM sleep, respectively.

## RESULTS

### High-frequency events in PFC during REM sleep

We simultaneously recorded local field potential (LFP) and neuronal populations in CA1 (n = 1,468) and PFC (n = 1,151) from animals (n = 10 rats) during REM sleep (**Figure S1A, Table S1**, REM duration = 164.6 ± 10.8 s/epoch for the 36 epochs included in analysis) throughout the course of learning a spatial memory task that requires hippocampal-prefrontal interactions^39,40^ (**Figure S1B**). Activity was continuously acquired across multiple run (8 epochs) and sleep (9 epochs) sessions during learning within a single day.^40^ We separated NREM and REM sleep stages based on theta-to-delta (TD) ratio in CA1,^15,16^ which revealed coherent shifts in TD ratio across CA1 and PFC at the onset and offset of REM sleep (**Figures 1A-C**). Waking periods showed higher movement and intracranial EMG compared to REM sleep, validating the behavioral state designations (**Figures S1C-I**).

**Figure 1.**
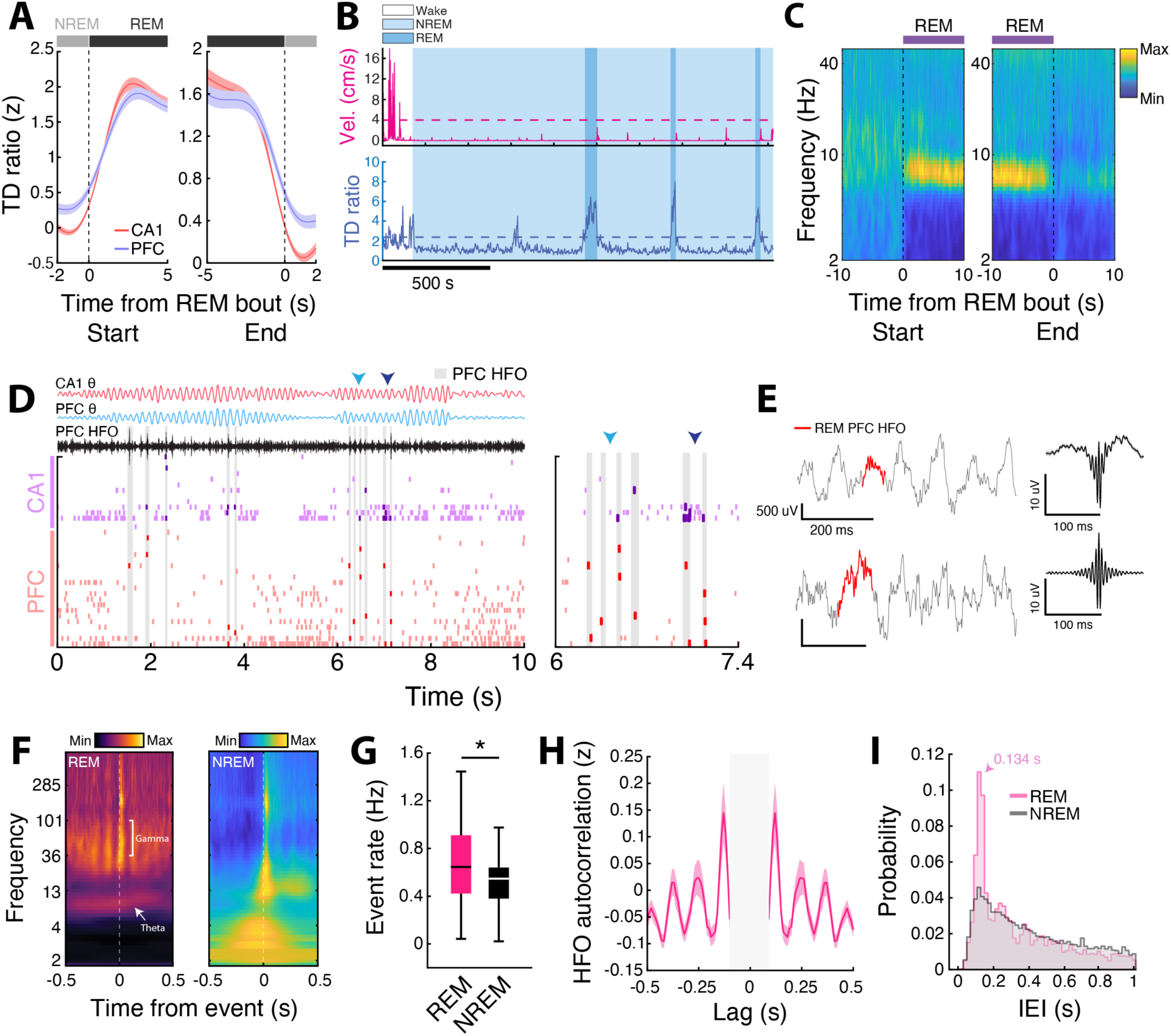
High-Frequency Oscillations (HFOs) in PFC during REM sleep. (**A**) CA1 and PFC theta-delta (TD) ratio time locked to either the start or end of a detected REM sleep bout. Start and end times were determined using CA1 TD ratio. Note the similar increases and decreases in CA1 and PFC TD ratio surrounding REM bout starts and ends, respectively. (**B**) An example sleep session for an animal showing each behavioral state detected using the sleep state algorithm that utilizes theta-to-delta ratio (TD ratio) and animal velocity (cm/s). (**C**) Average spectrograms across all REM bouts centered on the start (left) and end (right) of each bout. Note the sharp increase and decrease in theta power at the start and end of REM sleep, respectively. (**D**) Raster plot of CA1 and PFC cells during REM sleep with PFC theta and HFO amplitude plotted above. Gray shaded areas indicate PFC HFOs, and arrows denote periods expanded on the right to show single unit spiking during HFO events. (**E**) (Left) Example PFC HFOs during REM sleep. Note the theta frequency fluctuations in the local field potential. (Right) Example grand average PFC HFO waveform (top) and amplitude (bottom) across all events in an example animal. (**F**) Example PFC event triggered spectrogram for REM HFOs and NREM ripples. Note the elevated gamma (40-100 Hz) and theta (6*-*12 Hz) band activity associated with REM HFOs, in contrast to the elevated slow oscillation (0.1-4 Hz) and spindle (12-16 Hz) band power in during NREM ripples. (**G**) PFC high-frequency event rates in REM (HFOs) and NREM (ripples) sleep (REM = 0.66 ± 0.06, NREM = 0.54 ± 0.05, *p = 0.013, WSR). (**H**) PFC HFO autocorrelation in REM sleep. Note the peaks in the autocorrelation that repeat approximately every 125 ms. (**I**) Distribution of inter-event intervals (IEIs) for NREM ripples and REM HFOs. Note the prevalence of IEIs less than 200 ms for REM HFOs (Mode = 0.134 s). Only IEIs up to 1 s are shown for visualization purposes.

To investigate the high-frequency component of phasic burst activity in PFC, cortical ripples during NREM sleep and HFOs during REM sleep were detected as transient events in the 150-250 Hz range, as previously described^28^ (**Figures 1D,E** and **S2A,B**). Similar criteria were used to detect cortical high-frequency events in NREM and REM states; however, to avoid ambiguity and to conform to previous nomenclature,^21,22^ we refer to cortical NREM events as ripples, cortical REM events as HFOs, and hippocampal sharp-wave ripples in NREM as SWRs throughout. Cortical event rates during REM were higher than NREM, and REM HFOs were associated with elevated theta and gamma power,^21,31^ in contrast to spindle- and slow oscillation-coupled ripple events in NREM^28,41^ (**Figures 1F,G**). Interestingly, examining the temporal structure of event occurrence, using both the event autocorrelation (the likelihood of finding another HFO at a given time lag) and the distribution of inter-event-intervals (IEIs, the time between consecutive HFOs), we find that REM HFOs often recur at intervals of ∼130 ms (**Figures 1H,I**) as HFO “chains”, which is consistent with theta frequency modulation and elevated theta power during these events (**Figure 1F**).

REM HFOs are associated with fast, theta timescale fluctuations in PFC population/multiunit activity (MUA) during transient periods of locally reduced activity, as evidenced by the overall decrease in population activity during these events (**Figure 2A**). Furthermore, to isolate the spectral component driving this PFC population response, we re-detected HFOs using a broader 100–250 Hz passband and separated them by co-occurrence with the original 150–250 Hz HFOs (coordinated vs. non-coordinated). HFOs that occurred independently of 150–250 Hz detections elicited little to no phasic PFC spiking modulation, whereas those temporally coordinated with 150–250 Hz events reproduced the phasic theta-modulated response, indicating that this phasic PFC response is selectively associated with HFOs containing prominent 150–250 Hz activity (**Figure 2B**). This phenomenon was also present when using different parameters for HFO detection or REM sleep designation (**Figures S2D-F**). In contrast, NREM cortical ripples were associated with a singular burst of activity spanning several hundred milliseconds^28^ (**Figures 2C** and **S2C**). Thus, these findings establish distinct high-frequency event-associated activity profiles in PFC that may underlie differential processing of memory related information during NREM and REM sleep.

**Figure 2.**
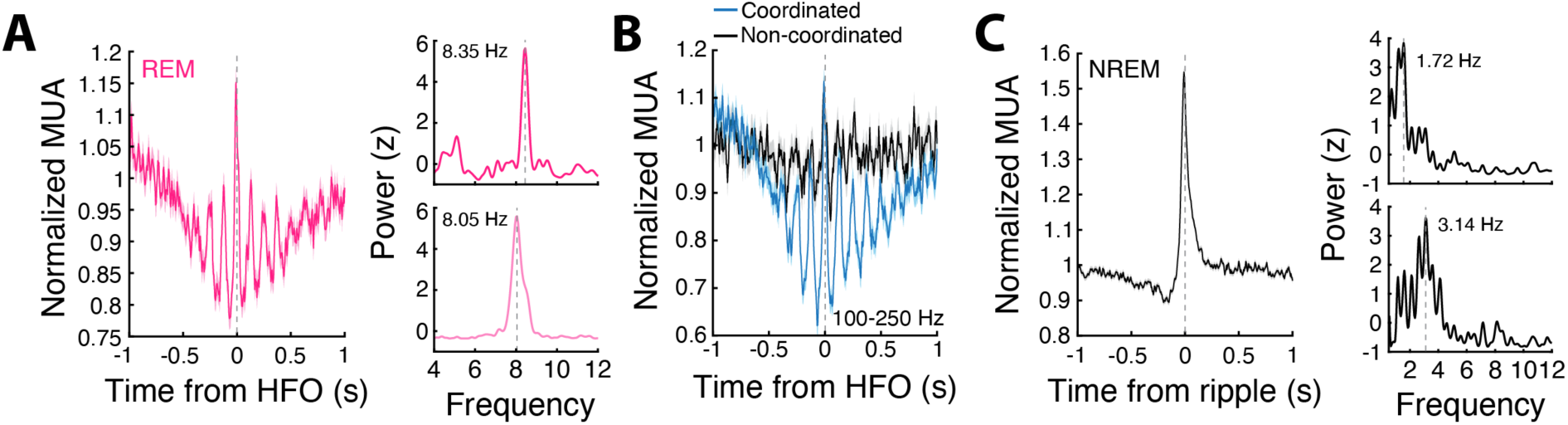
HFOs in PFC during REM sleep are associated with theta-modulated population activity and transient suppression. (**A**) (Left) REM HFO aligned multiunit activity (MUA) in PFC (n = 10 animals, 36 epochs), and (Right) the power spectral density (PSD) calculated for two example animals. The vertical dashed gray lines indicate the peak theta frequency of the population activity fluctuations surrounding HFO events. (**B**) PFC multiunit activity aligned to REM HFO events extracted from a wider frequency range (100-250 Hz). The separation into coordinated and non-coordinated events (Coordinated = 48.75 ± 2.6%) was determined by the overlap of these HFOs with HFOs extracted using the standard frequency range (150-250 Hz). This analysis was performed to demonstrate that the HFOs extracted from the 150-250 Hz range primarily drives the theta frequency multiunit activity and transient suppression in PFC. (**C**) Same as in (**A**) but for NREM ripples. Note the synchronous activity burst time-locked to NREM ripples, in contrast to the distinctive activity pattern during REM HFOs in (**A**). Top and bottom PSDs are from the same animals as in (**A**).

### Coupling of theta, gamma, and prefrontal HFOs in REM sleep

Theta-gamma phase-amplitude coupling (PAC) is one of the most widely studied cross-frequency coupled phenomena in the brain and involves the coupling of distributed, lower frequency theta oscillations and local higher frequency bursts.^42^ Thus, cross-frequency PAC is considered to be a physiological marker that links distributed brain regions during bouts of coherent activity to integrate information across different spatiotemporal scales.^42^ Furthermore, previous work has proposed that phasic REM sleep may support cortical-hippocampal dialogue through this theta frequency-based coordination.^43^

Phasic REM is characterized by higher theta power and frequency as compared to tonic REM in both the neocortex and hippocampus.^18,19^ Thus, we detected putative bouts of phasic REM as periods of elevated theta power (**Figure S3**). We further quantified the inter-peak-intervals of the filtered CA1 theta signal during low and high theta power bouts to determine whether they may correspond to putative tonic and phasic substages, respectively. Inter-peak intervals were significantly shorter during high theta power bouts than low power bouts, consistent with these periods reflecting putative phasic REM (**Figures S3A-C**). The rate of REM HFOs, as well as chaining of events, was significantly elevated during periods of high theta power (**Figures S3D-G**), indicating that the prevalence of REM HFO chains is elevated during bouts of putative phasic REM sleep. Consistent with this, we found that theta cycles were more likely to contain embedded HFOs during putative phasic REM than during tonic REM (**Figure 3A**). Thus, we restricted further analyses for quantifying gamma and HFO coupling to theta oscillations to these phasic bouts of high theta power (**Figures 3B,C** and **S4A-F**). To further validate the coupling of theta, gamma, and HFOs (illustrated in **Figure 1F**), we quantified PAC, which is a measure of how strongly the amplitude of fast oscillations vary as a function of the phase of a slower one, between theta phase and gamma to HFO frequency (40-250 Hz) amplitudes (**Figure 3B**). Examining the modulation indices at two discrete frequency bands, gamma (40-100 Hz) and HFO (150-250 Hz), revealed a higher frequency, HFO component that is also modulated by theta phase (**Figures S4A-F**). The modulation index summarizes this coupling as a single number, with higher values indicating tighter phase-locked amplitude modulation.

**Figure 3.**
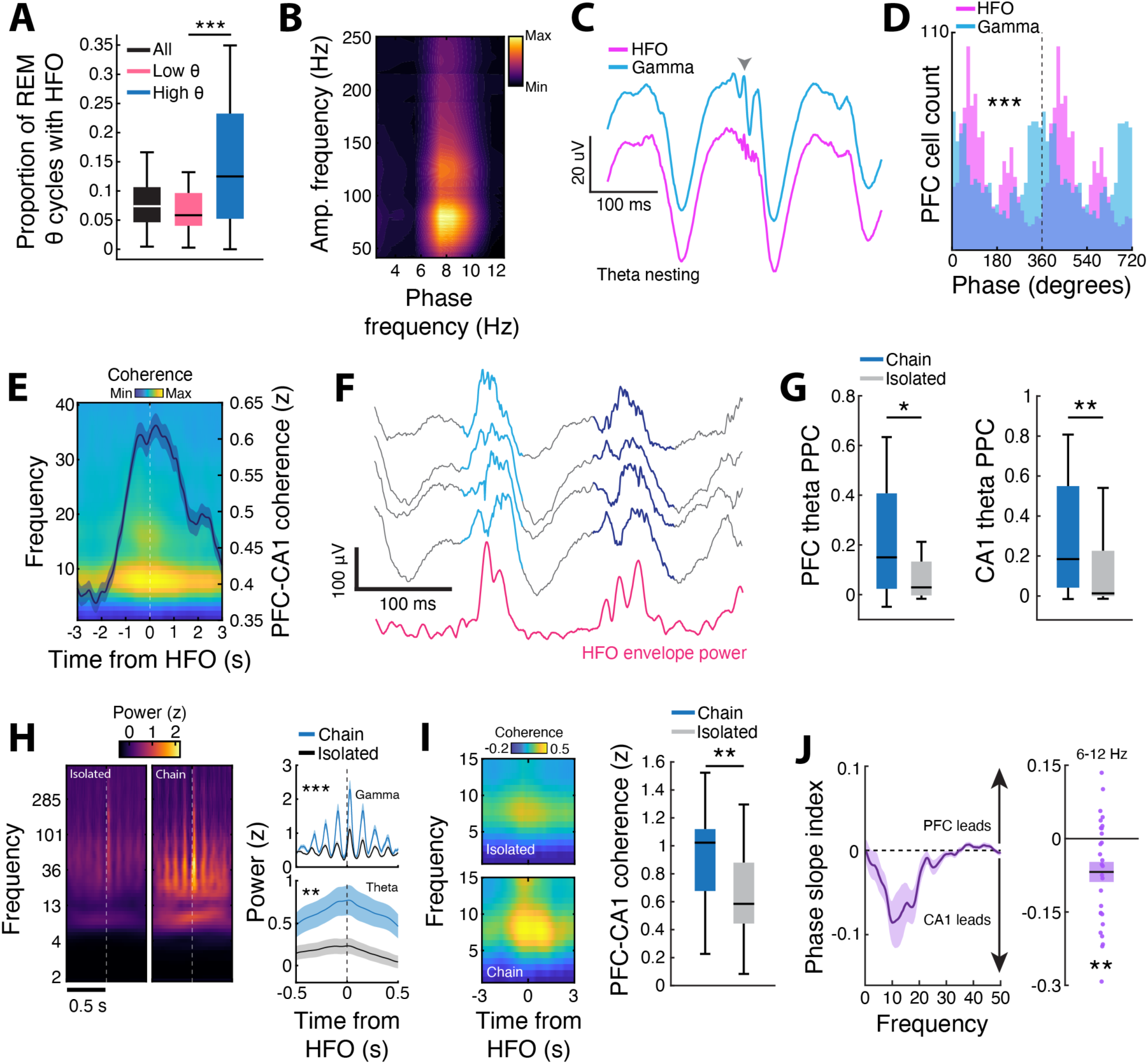
Theta, gamma, HFO coupling underlies distinct REM modulation states in PFC. (**A**) Proportion of theta cycles with HFOs across all, low theta power (putative tonic), and high theta power (putative phasic) cycles (All = 0.075 ± 0.007, Low = 0.065 ± 0.006, High = 0.140 ± 0.018, ***p = 4.90×10^-5^, WSR). (**B**) Phase-amplitude coupling (PAC). Averaged modulation across phase-amplitude pair occurrences during high theta power REM periods. The phase of the theta oscillation at a peak frequency of 7-8 Hz strongly modulates the amplitude of higher frequency oscillations. (**C**) Theta nesting for gamma and HFO frequency oscillations for an example tetrode illustrating coupling of theta, gamma, and HFO oscillations in PFC during REM sleep (see **Methods**). Briefly, averaged nesting plots were generated by aligning the unfiltered LFP to the peak power in the preferred theta frequency phase bin for either gamma or HFO oscillations separately. (**D**) Distribution of preferred phases for PFC cells locked to HFO or gamma events (U^2^ = 4.74, ***p = 4.56×10^-41^, Watson’s U^2^ test). (**E**) CA1-PFC theta (6-12 Hz) coherence centered on PFC REM HFOs is elevated relative to baseline theta coherence. The overlaid curve on the spectral plot represents the mean z-scored coherence across epochs, with shading indicating ± SEM. (**F**) Example REM HFO chain (doublet) plotted across 4 separate tetrodes with the average HFO power plotted below. (**G**) HFO chains showed higher PFC (Left) and CA1 (Right) theta PPC (pairwise phase consistency, a measure of phase-locking strength) compared to isolated HFOs (PFC: Chain = 0.22 ± 0.06, Isolated = 0.09 ± 0.04, *p = 0.013, WSR; CA1: Chain = 0.30 ± 0.08, Isolated = 0.14 ± 0.05, **p = 0.0016, WSR). (**H**) (Left) Chain and isolated HFO triggered spectrograms. (Right) Power in the gamma (top, 40-100 Hz) and theta (bottom, 6-12 Hz) frequency bands for isolated and chained HFOs. The average power in each band in the ±500 ms window surrounding HFO onset was used for comparison (Gamma: Chain = 0.72 ± 0.06, Isolated = 0.45 ± 0.03, ***p = 8.12×10^-7^, WSR; Theta: Chain = 0.60 ± 0.20, Isolated = 0.15 ± 0.11, **p = 0.0021, WSR) (**I**) (Left) CA1-PFC theta (6-12 Hz) coherence during isolated vs chained events and (Right) quantification of the peak theta coherence in a ±1 second window from HFO onset (Chain = 0.93 ± 0.08, Isolated = 0.65 ± 0.05, **p = 0.0069, WRS). (**J**) (Left) Phase slope index (PSI), which is a measure of phase lag consistency across different frequencies, peaked at theta frequency, indicating a CA1 to PFC directed flow of information during HFO chains. (Right) Quantification of PSI in the theta band (6-12 Hz). Each data point corresponds to the averaged PSI across each electrode pair in an epoch. (PSI = -0.068 ± 0.021, **p = 0.0025, t-test against zero).

To further verify that gamma and HFO are separable rhythms, we examined theta nesting of both oscillations, which quantifies the oscillatory components of the average raw LFP waveform centered on peaks of gamma or HFO power within the preferred theta phase bin (see **Methods**). This revealed two distinct fast oscillatory components embedded within the theta cycle that are tightly coupled, with gamma events leading HFOs (**Figures 3C** and **S4C,D**). Since the coupling metrics used here quantify relationships across theta cycles in aggregate rather than on a cycle-by-cycle basis, analyses of theta coupling for gamma and HFO activity were restricted to periods of high theta power (putative phasic REM), as mentioned above. Including periods of relatively low HFO occurrence could obscure genuine coupling dynamics. Focusing on epochs with robust theta oscillations, where HFOs are more prevalent, therefore provides a stronger basis for distinguishing gamma from HFOs and yields a more faithful representation of theta, gamma, and HFO cross-frequency coupling.

Furthermore, the phase locking preferences of PFC neurons to these two oscillatory components differ (**Figures 3D** and **S4G**), confirming gamma and HFO separation, and potentially reflecting distinct modes of local ensemble memory related processing. Importantly, we observed elevated CA1-PFC theta coherence during these REM HFOs as compared to baseline (**Figure 3E**), suggesting that increased theta coherence is associated with HFO generation in PFC during REM sleep.

Since we observed short-latency HFO recurrence during REM (**Figures 1H,I**), we reasoned that these HFO “chains” may be functionally different than HFOs that occur in isolation. Indeed, previous studies have characterized oscillatory events in multiple brain areas that occur in temporal clusters.^44-47^ Importantly, chains of spindle events (termed, “spindle trains”) in NREM sleep have been shown to underlie persistent reactivation of awake patterns and are associated with precise spatiotemporal coupling of sleep oscillations.^45^ Similar to this previous study, we find spindle train events in PFC during NREM sleep (**Figures S5A-D**), supporting the claim that PFC can exhibit temporally clustered events during sleep.

We identified chains of HFOs as events that occur within 200 ms of each other (37.9% of all events). Chains were more prevalent in REM than NREM sleep (**Figure S5E**) and in putative phasic REM than tonic REM (**Figures S3E,F**). Similar to spindle trains, REM HFO chains predominantly occurred in doublets (**Figures 3F** and **S5C,G**). REM HFOs were more consistently locked to a preferred theta phase than NREM ripples, as quantified by pairwise phase consistency (PPC), which is a bias free measure of phase-locking strength (**Figure S5F**). Furthermore, REM HFO chains were strongly phase modulated by theta in both PFC and CA1 and had similar properties to isolated HFOs (**Figures 3G** and **S5J**). The strength of gamma was also elevated during HFO chains and exhibited phasic power fluctuations as well as suppression of population activity (as in **Figure 2A**) when coupled (within 500 ms) with HFOs, further demonstrating strong temporal coupling (**Figures 3H** and **S4I,J**).

Corroborating the strong phase locking of chain HFOs to theta oscillations in PFC and CA1 (**Figure 3G**), we find that hippocampal-prefrontal theta coherence is elevated during chain HFOs as compared to isolated events (**Figure 3I**), with CA1 theta leading PFC theta, as indicated by the phase slope index (PSI; negative values denote a CA1 to PFC direction of information flow at theta frequencies; **Figure 3J**). Together, these findings suggest that chains of HFOs can underlie coordinated hippocampal-prefrontal activity that supports REM sleep-mediated memory consolidation.

### Differential modulation of PFC neuronal activity during NREM prefrontal ripples and REM prefrontal HFOs

Since we observed theta-modulated population activity aligned to REM HFOs that differed from activity aligned to NREM ripples (**Figures 2A-C**), we next examined PFC responses to determine how activity surrounding these events are organized at the level of single units and ensembles. Similar to population responses, many PFC neurons exhibited oscillatory spiking aligned to REM HFOs (**Figure 4A**). Unlike NREM ripples, REM events were associated with a broader distribution of activity peaks, suggesting a more sequential structure of neuronal firing surrounding these events (**Figures 4A,B**). Furthermore, we trained a binary linear classifier on the spike count vector across PFC neurons during each event and tested its ability to label held-out events as NREM ripples versus REM HFOs (10-fold cross-validation, with event counts equalized across types). PFC neuronal activity reliably distinguished NREM and REM events, confirming differences in spiking content (**Figure 4C**). In line with elevated CA1-PFC theta coherence surrounding REM HFOs (**Figure 3E**), we found that a subset of CA1 neurons were modulated by these events (**Figure 4D**), indicating enhanced interregional coactivity, potentially delineating periods of CA1-PFC ensemble reactivation to support memory consolidation.

**Figure 4.**
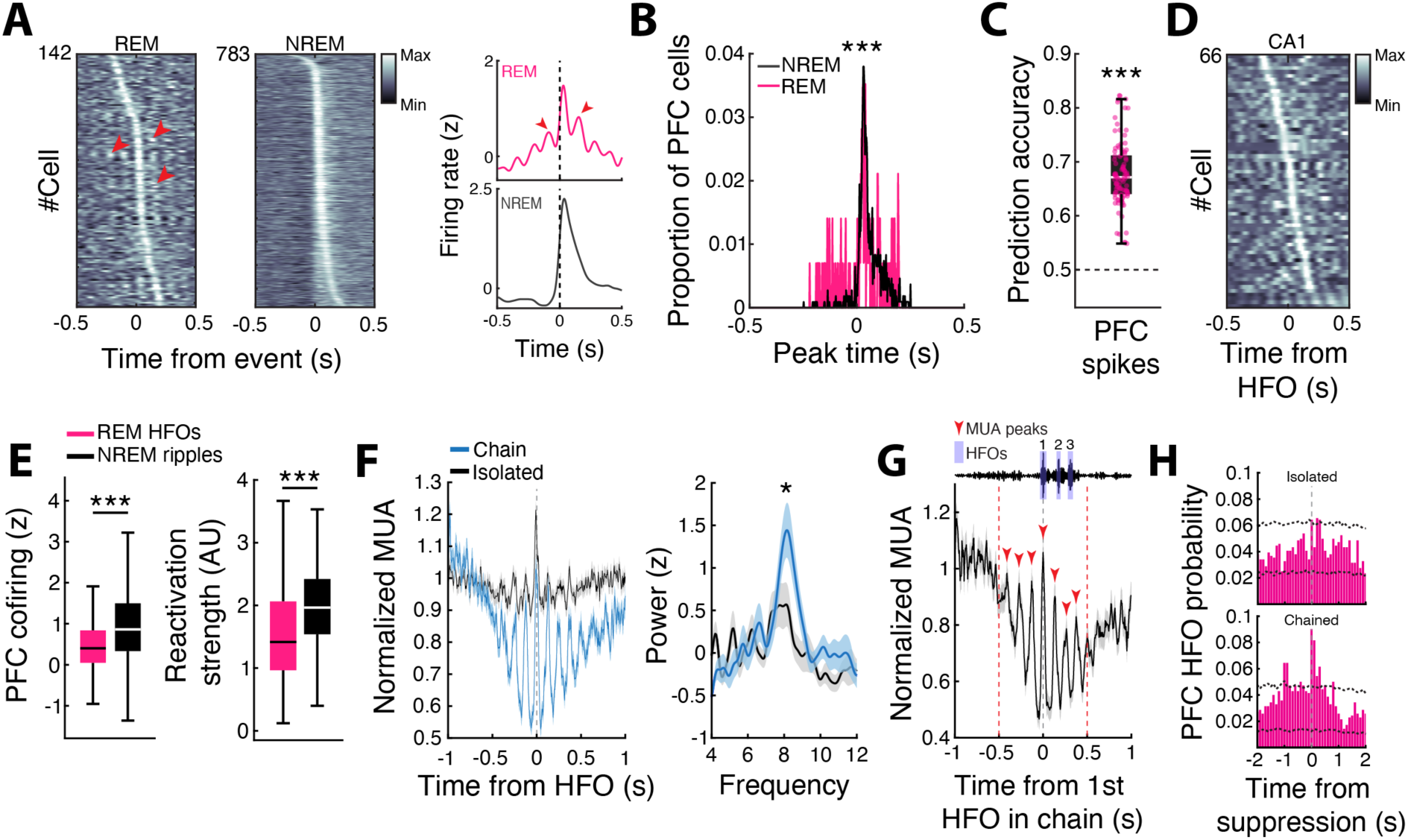
Differential modulation of PFC neuronal activity during NREM prefrontal ripples and REM prefrontal HFOs. (**A**) (Left) PFC neurons that were positively modulated (EXC) during REM PFC HFO events and NREM PFC ripple events (REM EXC modulation = 142 out of 922 candidate neurons; NREM EXC modulation = 783 out of 1079 candidate neurons; See Methods section Ripple/HFO aligned modulation). (Right) Average z-scored peri-event time histograms (PETHs) aligned to events. Note the peaks in activity adjacent to REM HFO alignment at theta timescale denoted by the red arrowheads. (**B**) Timing of peak excitation for significantly modulated PFC cells during NREM and REM. Note the broader distribution of peaks during REM sleep (Comparison of peak timing distributions, ***p = 8.29×10^-5^, WRS). (**C**) PFC event type (NREM vs REM) can be predicted by population spiking activity (PFC data = 0.68 ± 0.01, shuffle = 0.50 ± 8.00×10^-5^, ***p = 2.56×10^-34^, WRS). (**D**) Modulation of putative CA1 pyramidal neurons aligned to PFC REM HFO events (REM EXC modulation = 66 out of 680 candidate neurons). (**E**) PFC cofiring and assembly reactivation strength were overall stronger during NREM ripples (Cofiring: NREM = 0.96 ± 0.03, REM = 0.48 ± 0.03, ***p = 3.58×10^-27^; Reactivation strength: NREM = 2.01 ± 0.05, REM = 1.56 ± 0.07, ***p = 6.91×10^-16^, WRS). (**F**) (Left) PFC multiunit activity aligned to chained vs isolated REM HFO events. (Right) HFO aligned multiunit PSD illustrating stronger theta-modulated spiking activity in PFC during chained events (mean power in the 6-10 Hz band, *p = 0.029, WRS). (**G**) PFC multiunit activity aligned to the first HFO in each chained event. At top is a schematic of the multiunit alignment procedure. The initial HFO in a chain is at time 0. Note the peaks in the multiunit activity prior to the initial HFO within a chain, suggesting that fluctuations in PFC multiunit activity precede chained events. (**H**) The probability of (Top) isolated and (Bottom) chained REM PFC HFOs centered on population suppression events detected in REM sleep. Note the strong peak around 0 in the HFO chain distribution, indicating that REM PFC HFO chains are associated with overall decreases in population activity as compared to isolated HFOs. The black dotted lines indicate the 99% confidence intervals based on suppression events extracted from shuffled multiunit activity.

Overall cortical cofiring, which is a measure of coincident activity between neuron pairs during discreet events,^48^ and assembly strength were reduced during REM HFOs relative to NREM ripples (**Figure 4E**), consistent with overall activity suppression during HFO chains (**Figures 4F-H**), which can create a permissive background for gating selective reactivation of PFC ensembles in REM sleep. Examining population spiking activity, we found that PFC activity was strongly theta-modulated during REM HFO chains on the background of transient local suppression, compared to both isolated and NREM events (**Figures 4F-H** and **S5H-J**). This phasic population activity pattern was seen during HFO chains in both putative phasic and tonic REM substates (**Figure S3F**).

To address the possibility that this theta-modulated spiking activity and suppression may simply reflect high-theta-power bouts (rather than HFO occurrence), we generated surrogate event times by randomly sampling time points that matched the preferred theta phase of HFOs, but were at least 250 ms away from any detected HFO. PFC activity aligned to these surrogate events did not exhibit the same theta-modulated activity associated with real HFOs, even when the surrogate events were drawn from the highest-theta-power quartile (**Figure S5K**). Furthermore, we examined population activity as a function of distance from detected HFOs across all REM bouts, or subset by putative phasic REM, and did not observe the same magnitude of phasic response (**Figure S5L-N**), confirming that the transient theta-modulated population activity in PFC is specifically associated with the occurrence of HFOs. Interestingly, only HFO chains were uniquely associated with a transient suppression in overall local population activity (**Figures 4F-H** and **S6**), which is in contrast to the strong excitation seen in PFC and CA1 during NREM PFC ripples and SWRs, respectively.^28^ Since REM HFOs predominantly occur in doublets (**Figure S5G**), the suppression of overall population activity is consistent with sublinear integration, in which subsequent events elicit less spiking activity than the first, thus likely creating privileged temporal windows for selective PFC ensemble reactivation and strengthening.

### PFC ensemble activity is sequentially organized by REM HFO chains

Since we observed a broader temporal distribution of spiking activity (**Figure 4B**) as well as multiple peaks in population activity during REM HFO chains (**Figures 4F,G**), we next examined, in more detail, how ensemble activity is structured around these events. Previous studies have shown that the refractoriness of certain oscillatory events, such as spindles and SWRs, create distinct segregated periods for selective population activation.^44,45,47,49^ Thus, we first examined population dynamics based on the temporal separation of REM HFOs. We observed selective activation of subsets of PFC ensembles during individual events in HFO chains (**Figure 5A**). To quantify this structure, we represented each HFO as a binary vector of PFC neurons active during the event, and computed the Pearson correlation between the vectors of every pair of consecutive HFOs. Spike content similarity as a function of Inter-Event-Interval (IEI) was examined using quartile ranges for the intervals between HFOs. Quartile 1 had the highest correlation, with the average IEI of this quartile as 114 ms (theta frequency, corresponding to HFO chains), indicating that adjacent HFO events within chains show a high degree of similarity as compared to adjacent isolated events (**Figures 5A,B** and **S7A**). Furthermore, we examined whether the order in which individual PFC neurons fired during HFOs within a chain was preserved across chains. For each chain, we extracted each cell’s first-spike rank order and compared it to a leave-one-out template constructed from the average normalized rank across all other chains. This procedure revealed preserved sequences of PFC activity across multiple chain events (**Figures 5C** and **S7B**). These findings suggest that HFO chains stereotypically organize PFC activity, likely to support sequential reactivation of distinct assemblies.

**Figure 5.**
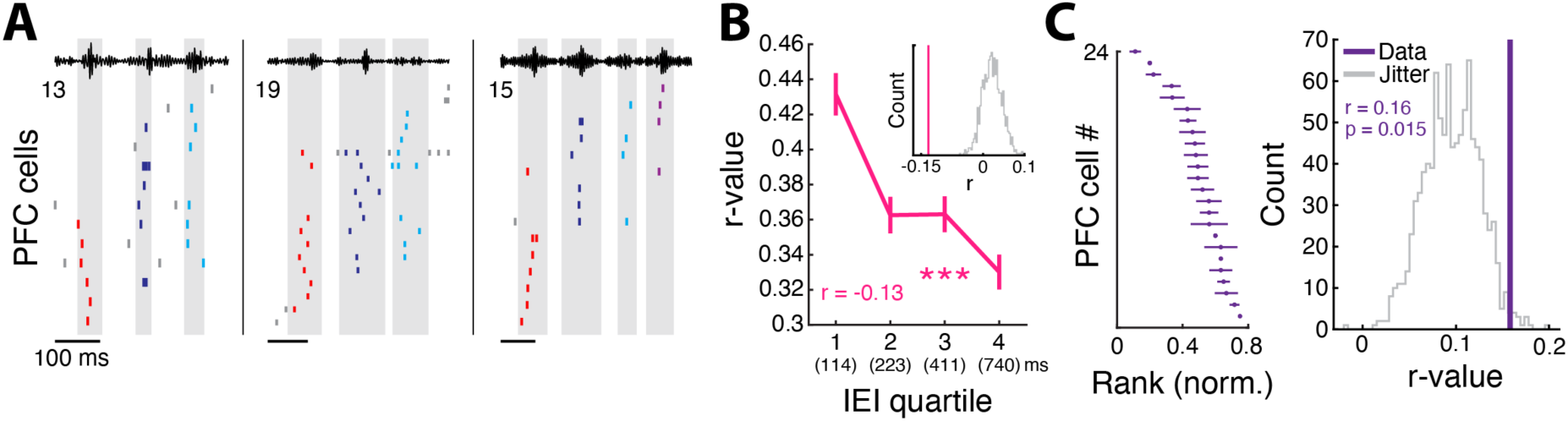
Temporally constrained sequential activity in PFC. (**A**) Example rasters during HFO chains. Spikes are color-coded based on the HFO number within the chain during which the cell was active. Note the different cell assemblies active during each HFO within a chain. (**B**) Similarity of PFC population spiking between adjacent REM HFOs and its relationship with inter-event-intervals (IEI). Here, the PFC spiking activity during each HFO was binarized across all neurons and the Pearson correlation coefficient was calculated between adjacent events as a measure of pattern similarity. Then the relationship between pattern similarity and IEI was reported (r = -0.13, ***p = 6.00×10^-8^, Pearson correlation). Below each quartile is the average IEI of that quartile in milliseconds. Note that the average for quartile 1 is 114 ms, which corresponds to 8-9 Hz during HFO chains. (Inset) The Pearson correlation coefficient compared to a distribution of r values generated from shuffling (n = 1000) HFO identity. (**C**) Rank-order correlation analysis. (Left) Example rank-order template from an example sleep epoch. Templates were generated by concatenating all HFOs in chained events (inter-event periods excluded) and averaging the normalized rank across chains. (Right) The mean rank-order correlation from the leave-one-out cross-validation procedure. Each event’s rank was correlated with the averaged rank across all other events. The average across all events compared to a distribution of means generated by jittering (n = 1000) spike times is shown.

Next, to test our hypothesis of sequential reactivation, we assessed PFC assemblies by examining time-locking of PFC reactivation strength to NREM and REM events. We detected behavioral PFC assemblies that are reactivated in sleep using established methods.^50,51^ Similar to single unit modulation, the timing of PFC assembly reactivation aligned to NREM ripples was largely restricted to within 200-500 ms after ripple onset, depending on the ripple type examined (**Figures 6A** and **S7F**). For REM events, only the first HFO within a chain was selected for alignment, since we found that peaks in population activity occur prior to the onset of HFO chains (**Figure 4G**), possibly facilitated by strong phasic gamma surrounding HFO chains (**Figures 3H** and **S4I**). We found that unlike NREM ripples, where assembly reactivation is synchronous and consistently highest at the onset of events, PFC assemblies were sequentially organized on longer timescales surrounding REM HFOs, similar to single unit modulation (**Figures 4A,B** and **6A**). This temporal tiling of assembly reactivation aligned to REM HFOs manifested in a relatively flat average response, highlighting the differential engagement of PFC assemblies during NREM and REM events (**Figure 6B**).

**Figure 6.**
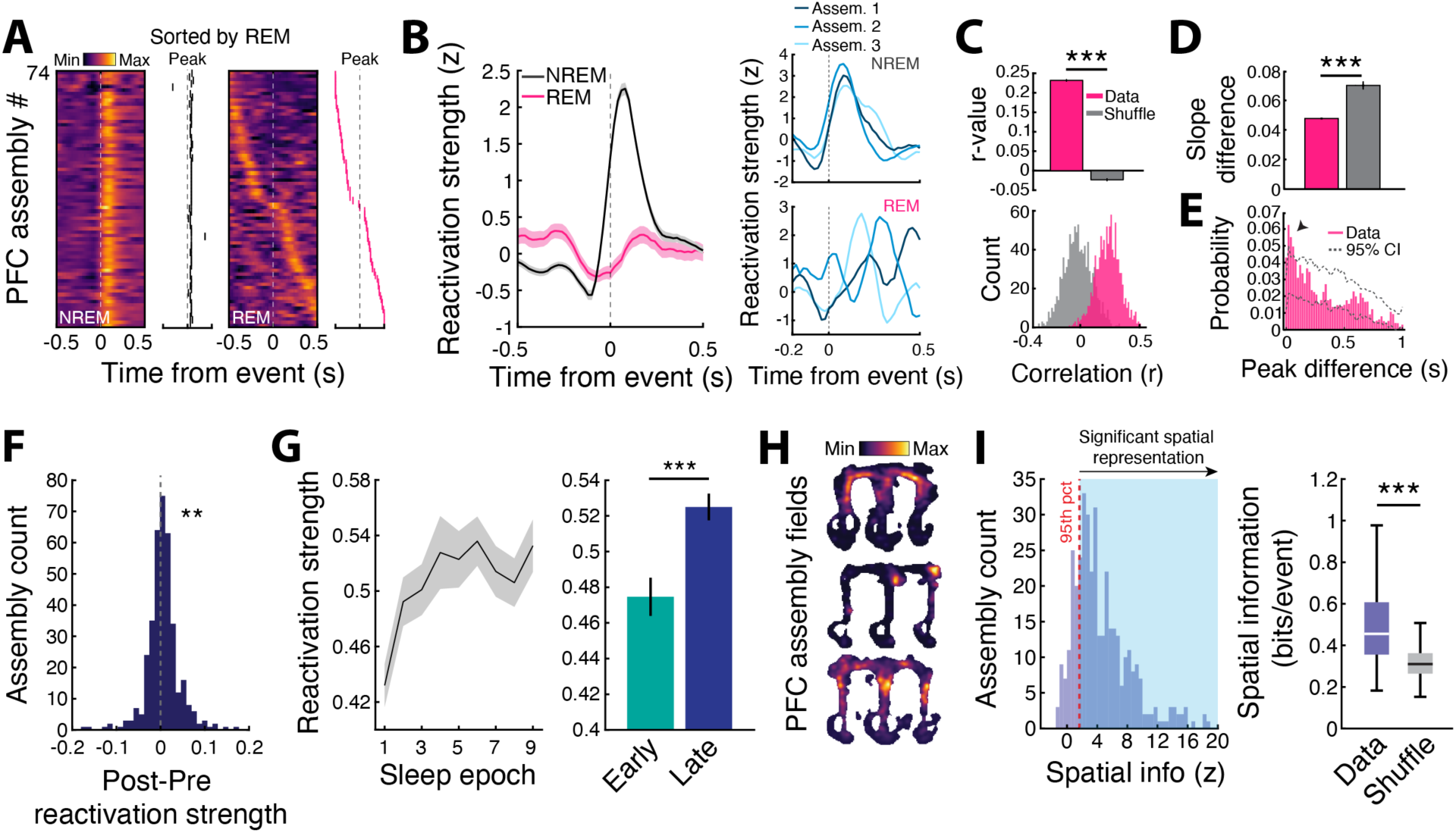
Structured reactivation in PFC during REM HFOs. (**A**) (Left) PFC assembly reactivation aligned to NREM ripples or (Right) the first HFO of each REM HFO chain. Reactivation peak locations are shown alongside each plot. The assemblies are sorted based on the average peak reactivation bin within a ±500 ms window surrounding the first HFO in a chain. Note the tiling of assembly peaks aligned to HFO events in REM sleep, which manifests in the relatively flat average response in REM vs. a singular peak of synchronous assembly reactivation in NREM as shown in (**B**). (**B**) (Left) Averaged event triggered reactivation strength during NREM and REM. (Right) Mean reactivation strength of three example assemblies aligned to high-frequency events in NREM and REM, illustrating the tiling of assembly reactivation relative to events. The example assemblies shown come from the same animal and epoch. (**C**) Cross-validation of the REM reactivation sequences shown in (**A**), suggesting a consistent structure of reactivation surrounding REM HFO chains. (Bottom) Distributions of r values calculated from the Pearson correlation between peak reactivation bins across all assemblies for two randomly chosen halves (n = 1000 random splits) of the HFO aligned data (Data = 0.232 ± 0.003, Shuffle = -0.023 ± 0.004, ***p = 1.13×10^-264^, WRS). (**D**) Differences in slope of the linear fit between the time of peak reactivation (in milliseconds) and assembly identity for two halves of the HFO aligned data (Data = 0.048 ± 2.21×10^-4^, Shuffle = 0.070 ± 2.67×10^-4^, ***p = 2.31×10^-302^, WRS). (**E**) Probability of assembly peak displacement between randomly chosen halves of the HFO aligned data. For each assembly, the absolute difference in peak location was calculated between the two halves. To calculate 95% confidence intervals, assemblies were circularly shuffled, and the peak displacement probabilities were calculated over 1000 random splits of the data. Note the peak at small displacement values, indicating consistency of assembly peak activity across splits. (**F**) The distribution of PFC assembly reactivation strength differences between pre and post sleep. The same W-Track session was used as the template for each pair of sleep epochs. Thus, assemblies were matched across pre and post sleep epochs (**p = 0.006, t-test against zero). (**G**) (Left) PFC assembly reactivation strength during sleep over the course of learning. (Right) Comparison of reactivation strength during early (sleep sessions 1-3; epochs prior to grouped average of 75% correct performance on W-Track, see Figure S1B) and late (sleep sessions 4-9) sleep epochs (Early = 0.475 ± 0.010, Late = 0.525 ± 0.007, ***p = 1.78×10^-4^, WRS). Except for sleep epoch 1, the preceding W-Track session was used as the template for reactivation. Only neurons that were tracked throughout the entire experiment were included for analysis. (**H**) Example PFC assembly rate maps. (**I**) (Left) Distribution of z-scored spatial information values for real assembly maps compared to maps generated from circularly shuffled (n = 1000) activation times. Each assembly’s spatial information was expressed as a z-score relative to its shuffled distribution. The red dotted line at a z = 1.65 marks the one-tailed 95th percentile (p = 0.05). (Right) Observed spatial information vs. the median of each assembly’s shuffle null distribution (Data = 0.535 ± 0.013, Shuffle = 0.325 ± 0.005, ***p = 2.82×10^-65^, WSR).

To confirm that the sequential tiling of assembly reactivation peaks in **Figure 6A** was a reliable phenomenon associated with HFO chains rather than an artifact, we used a split-half cross-validation. HFO events were first randomly partitioned into two halves, each assembly’s peak reactivation bin was computed independently within each half, and the similarity of the temporal ordering of assembly peaks between the two halves was calculated by taking the Pearson correlation (see **Methods**). Consistent with the structured PFC activity during chain events (**Figures 5A-C**), we found the greatest degree of sequence similarity during REM HFO chains compared to shuffled data (**Figure 6C**), confirming the tendency for PFC assemblies to be reactivated in a stereotyped, sequential manner as compared to isolated REM HFOs and NREM ripples (**Figures S7C-F**).

Finally, consistent with task-related reactivation, we found stronger PFC ensemble expression during sleep epochs following experience (**Figure 6F**). Importantly, assembly reactivation in sleep strengthened over the course of learning (**Figures 6G** and **S1B**), and a large proportion of detected assemblies had behaviorally relevant task representations with high spatial information (**Figures 6H-I**).

### Differential engagement of distinct populations of CA1 neurons by PFC REM HFO chains

Theta coherence and phase amplitude coupling are two prominent modes of interregional coordination thought to underlie mnemonic processing.^42,52^ Since we observed an overall increase in CA1-PFC theta coherence and gamma power during HFO chains (**Figures 3H,I**), as well as sparse modulation of CA1 activity during REM HFOs (**Figure 4D**), we reasoned that there may be specific populations of CA1 neurons that are engaged by these events. Examining CA1-CA1 cofiring during these cortical HFO events, we observed a higher degree of spatial rate map correlation for high cofiring pairs, suggesting HFO specific reactivation of behaviorally relevant hippocampal assemblies to support memory consolidation^12^ (**Figures 7A,B**). Additionally, the absolute CA1 cofiring, comprising both negative and positive magnitudes of cofiring, was higher during HFO chains, indicating a distinct activity profile compared to isolated HFOs (**Figure S8A**).

**Figure 7.**
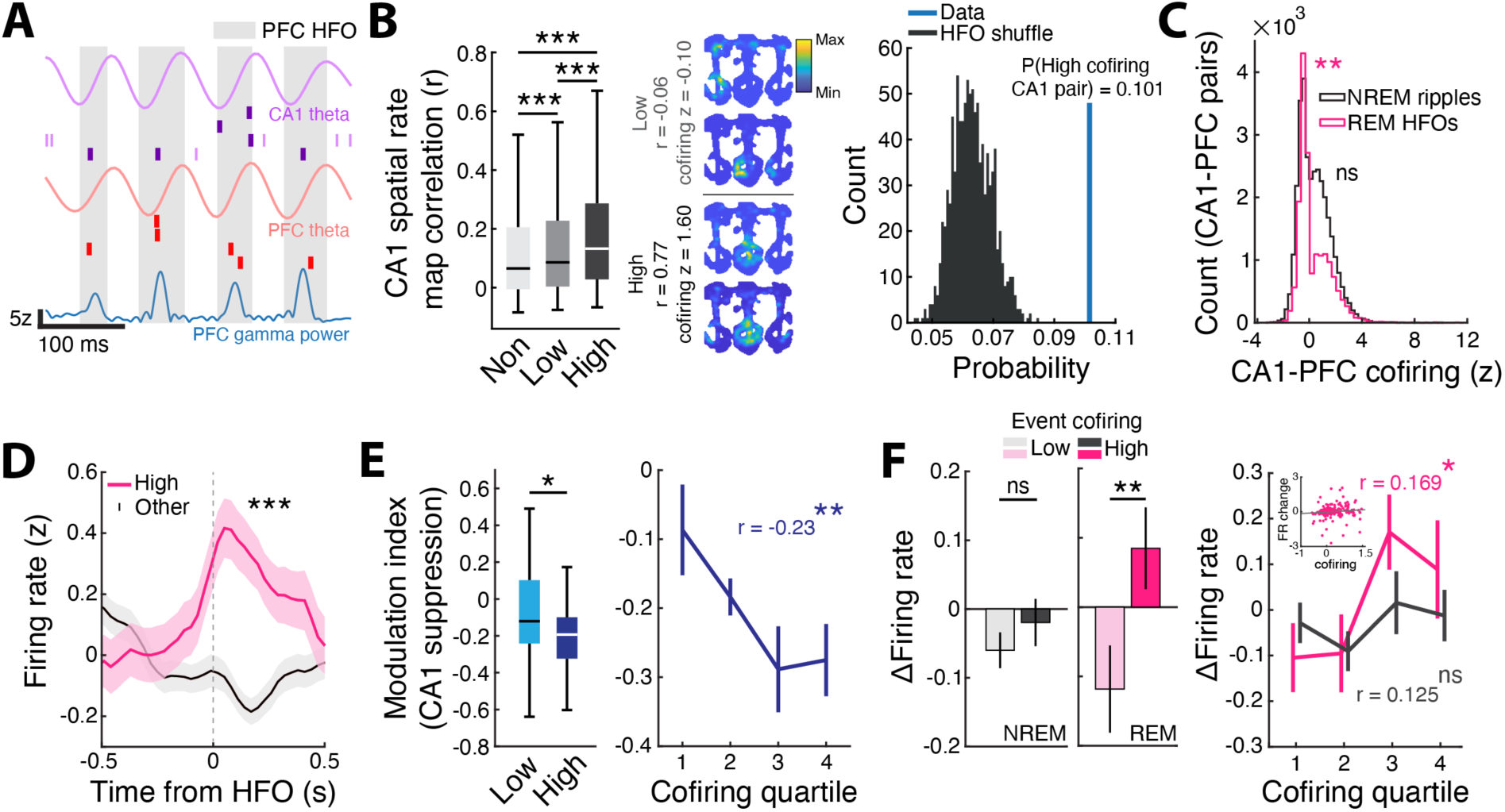
Differential engagement of distinct populations of CA1 neurons during PFC REM HFO chains. (**A**) Example raster plot of CA1 and PFC cells during a HFO chain event. Only cells that were active during the plotted time window are shown. Note the coherence of theta band activity in CA1 and PFC with elevated PFC gamma power during each HFO event in the chain. (**B**) (Left) 2D spatial rate map correlation for non-active, low, and high HFO chain cofiring CA1 pairs (Non-active = 0.123 ± 0.001, Low cofiring = 0.127 ± 0.001, High cofiring = 0.176 ± 0.004, Non vs Low, ***p = 1.26×10^-10^, Low vs High, ***p = 2.91×10^-21^, Non vs High, ***p = 2.33×10^-51^, Kruskal Wallis test with Bonferroni correction). Here, ‘non-active’ indicates cell pairs where cofiring was not assessed because at least one cell was inactive across all events. (Middle) Rate maps and spatial correlation of example low and high cofiring CA1 pairs. (Right) Real probability of high cofiring CA1-CA1 pairs during PFC HFOs (probability = 0.101) compared to a null distribution calculated by shuffling HFO times (n shuffles = 1000). (**C**) Histograms of CA1-PFC HFO cofiring during CA1-independent NREM PFC ripples and chained REM PFC HFOs. Cofiring during REM HFOs showed a bimodal distribution indicating separable populations of CA1 neurons based on PFC cofiring. (REM, **p = 0.001; NREM p = 0.23, Hartigan’s dip test for bimodality). (**D**) Firing rate of high cofiring and “other” CA1 cells aligned to HFO chains. Here, ’other’ includes low cofiring cells and cells that did not have a cofiring value because at least one of the cells in the pair emitted zero spikes across all events (similar to ‘non-active’ above). (Comparison of mean firing rates in the ±100 ms window from HFO onset, ***p = 7.77×10^-5^, WRS). For comparisons between cofiring and other metrics (e.g. firing rate), a single cofiring value was calculated for each neuron by averaging the cofiring metric across all neuron pairings. Additionally, high and low cofiring CA1 neurons were split based on average cofiring values > 0 and < 0, respectively. (**E**) (Left) High REM PFC HFO cofiring CA1 neurons exhibited a greater degree of suppression during NREM PFC ripples. For a description of modulation index, see Methods section Ripple/HFO aligned modulation. Here, since CA1 neurons exhibit a robust decrease in activity in response to NREM PFC ripples,^28^ we refer to the modulation as suppressive (Low cofiring = -0.08 ± 0.06, High cofiring = -0.25 ± 0.03, *p = 0.022, WRS). (Right) The CA1 cofiring magnitude during HFO chains was correlated with the degree of suppression during NREM PFC ripples (r = -0.23, **p = 0.0045, Pearson correlation). (**F**) (Left) Changes in CA1 firing rates from the first NREM bout to the last NREM bout within a sleep epoch plotted according to degree of cofiring with PFC (high, > 0; low, < 0), either during CA1-independent NREM ripples or chained REM HFOs (Low NREM = -0.06 ± 0.03, High NREM = -0.02 ± 0.03, p = 0.32; Low REM = -0.12 ± 0.063, High REM = 0.085 ± 0.059, **p = 0.0052, WRS). (Right) Firing rate change separated by cofiring quartile. The change in CA1 firing rates over the course of sleep was correlated with PFC cofiring during REM HFOs but not NREM ripples (NREM, r = 0.125, p = 0.46; REM, r = 0.169, *p = 0.012, Robust linear regression).

Next, we computed CA1-PFC cofiring across all cross-regional cell pairs and found that, during REM HFO chains, the distribution of cofiring values was bimodal, revealing a subpopulation of CA1 cells that are highly engaged with PFC neurons (**Figure 7C**). This distribution was unimodal during NREM PFC ripples, suggesting that the dual-population structure is specific to REM HFO chains. Analysis of low and high cofiring CA1 neurons during REM HFOs showed that high cofiring neurons exhibited elevated activity during HFO chain events as compared to low cofiring neurons (**Figure 7D**), independent of baseline firing rates (**Figure S9A**).

Interestingly, we also found a relationship between CA1 activation during REM PFC HFO chains and CA1 suppression by NREM PFC ripples. CA1 neurons that exhibited high cofiring with PFC (termed, “high HFO cofiring CA1 neurons”) specifically during REM HFO chains were more strongly suppressed during independent NREM PFC ripples, and the degree of suppression was correlated with the strength of REM CA1-PFC cofiring (**Figures 7E** and **S8C,D**). Given the sequential occurrence of NREM-REM sleep stages, this finding suggests that NREM ripples may tag specific CA1 populations for subsequent REM-dependent reorganization. However, although there was a strong relationship between CA1 suppression during independent PFC NREM ripples and excitation during coordinated SWRs in NREM sleep, as shown in a previous study^28^ (**Figures S8C,E**), we did not find correlations between CA1 REM HFO cofiring and CA1 SWR reactivation (**Figure S8F**), indicating that this CA1 sub-population that we identified as high co-firing with PFC neurons during REM HFOs is specifically engaged by high-frequency PFC events in sleep, with increased activity during REM and suppression during NREM.

Previous studies have shown that NREM and REM sleep are involved in modifying hippocampal excitability, with a net decrease in firing rates over the course of sleep.^33-35^ To test whether REM HFO engagement predicted shifts in hippocampal excitability across sleep, we computed each CA1 cell’s firing rate change from the first to the last NREM bout within a sleep epoch and asked whether this change was related to how strongly the cell co-fired with PFC during HFOs. Interestingly, we observed that high REM HFO cofiring CA1 neurons differentially shifted their firing rates upward over the course of a sleep session, whereas most of the neuronal population, including low REM HFO cofiring cells, decreased their firing rates (**Figures 7F**, **S8B**, and **S9B-E**), consistent with the role of REM sleep in hippocampal firing rate diversification.^53^ This effect was hippocampal specific; there was no differential regulation of PFC firing rates over the course of sleep (**Figure S9G**). Furthermore, the degree of cofiring with PFC during REM HFOs predicted overall CA1 firing rate changes during sleep, and this relationship was absent for NREM PFC ripples (**Figures 7F** and **S9E**). Splitting the population in half based on mean firing rate changes further confirmed this relationship (**Figure S9F**), suggesting a REM specific role of PFC HFOs in modifying hippocampal excitability.

### REM theta-phase shifting CA1 neurons are preferentially engaged during PFC REM HFOs

The dorsal CA1 pyramidal cell layer can be separated into subpopulations that are molecularly defined, have differential engagement with hippocampal SWRs and cortical areas,^54-57^ and employ complementary spatial codes for effective coding of environments.^58^ Notably, a subset of CA1 pyramidal cells preferentially shift their preferred theta phase from run to REM sleep^15^ – a phenomenon that is experience dependent.^36^ We therefore investigated whether there were differences in REM HFO recruitment for non-shifting and REM-shifting cells. We categorized REM-shifting neurons as cells that had a difference in phase preference greater than 90°.^15^ As expected, there was a subset of CA1 neurons that shifted theta phase preference (n = 124/355 neurons significantly locked to both run and REM theta, Shift magnitude = 133.14 ± 2.35°) and were strongly engaged by SWRs (**Figures S9H-J**), as previously reported.^55^ Similar to high cofiring CA1 neurons, these REM-shifting cells had higher cofiring with PFC than non-shifting cells (**Figure 8A**), and there was a significant relationship between HFO chain cofiring and suppression in the REM-shifting neurons, suggesting a specific modulatory effect on these neurons (**Figures 8B** and **S9K**). REM-shifting neurons also exhibited strong bursting activity during HFO chains as compared to isolated events and were more strongly phase locked to the PFC theta oscillation surrounding HFO chains (**Figures 8C,D** and **S9L**). Thus, PFC HFOs during REM sleep preferentially engage a specific population of REM-phase-shifting CA1 neurons, which are suppressed during NREM cortical ripples, and overall show an increase in firing rates during sleep, thus potentially increasing or stabilizing excitability to enhance signal-to-noise for efficient expression during future behavior.

**Figure 8.**
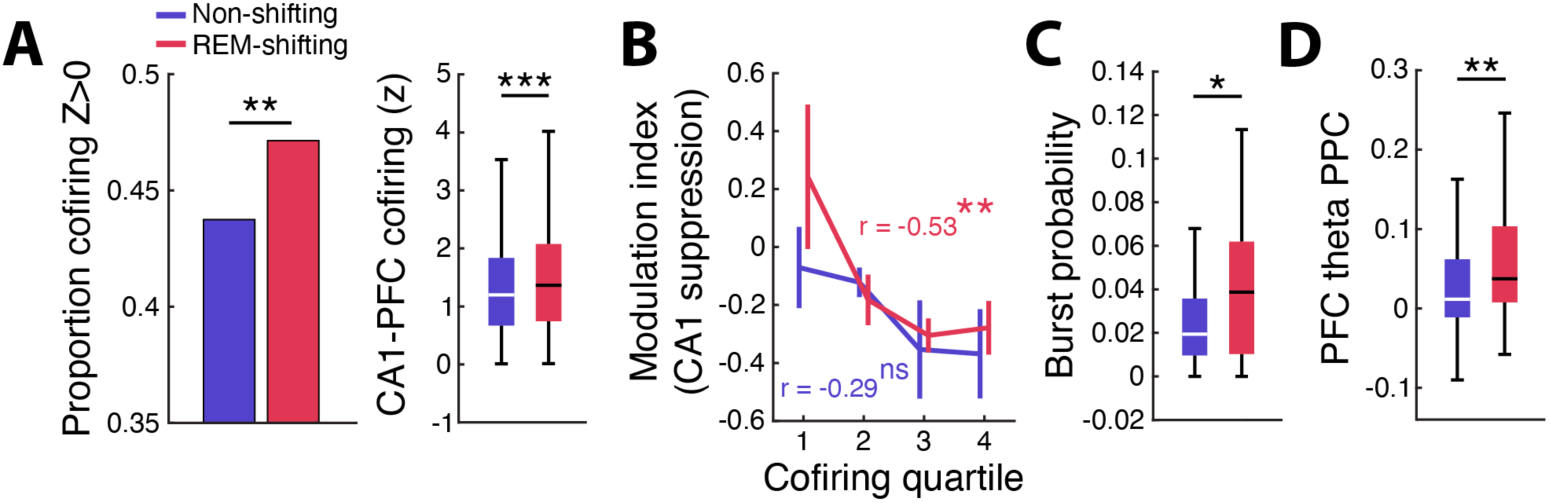
REM theta phase shifting CA1 neurons are preferentially active during PFC REM HFO chains. (**A**) (Left) Proportion of CA1-PFC pairs with cofiring greater than 0 during REM HFO chains that contained either non-shifting or REM-shifting CA1 neurons (Non-shifting = 2216/5066 pairs, 43.7%, REM-shifting = 1057/2242 pairs, 47.2%, z = 2.70, **p = 0.0097, Z-test for proportions). (Right) Magnitude of CA1-PFC HFO cofiring (greater than 0) compared for non-shifting and REM-shifting CA1 neurons (Non-shifting = 1.35 ± 0.02, REM-shifting = 1.51 ± 0.03, ***p = 8.30×10^-5^, WRS). Note that all cofiring pairs are assessed here. (**B**) Similar to Figure 7E, the REM PFC HFO cofiring magnitude of REM-shifting CA1 neurons was correlated with the degree of suppression during independent NREM PFC ripples (Non-shifting, r = -0.29, p = 0.066, REM-shifting, r = -0.53, **p = 0.0093, Pearson correlation). (**C**) Burst probability of non-shifting and REM-shifting CA1 neurons during REM HFO chains (Non-shifting = 0.028 ± 0.006, REM-shifting = 0.059 ± 0.017, *p = 0.022, WRS). (**D**) REM-shifting CA1 neurons showed higher PPC, indicating stronger phase locking to the PFC theta oscillation surrounding HFO chains (Non-shifting = 0.032 ± 0.006, REM-shifting = 0.064 ± 0.011, **p = 0.0026, WRS). Only periods surrounding REM HFO chains were used to evaluate phase locking strength of CA1 neurons.

### Modeling high-frequency event-associated PFC activity during NREM and REM sleep

To gain a better understanding of how a cortical network can exhibit different activity patterns during NREM ripples and REM HFOs, we built a simplified network model (**Figure 9A**). 1,600 excitatory pyramidal and 400 inhibitory interneurons were pseudo-randomly connected based on cortical layer 2/3 connectivity probabilities (see **Methods**). Acetylcholine (ACh) concentration is an especially important factor for differentiating NREM and REM sleep states, with strong impacts on cell-spiking properties and network dynamics.^59-61^ To incorporate one of the crucial impacts of ACh concentration on cortical neurons, each neuron in the model used a modified Hodgkin-Huxley (HH) formalism with an additional slow potassium (M-type) leak conductance term.^62^ During NREM simulations, this slow potassium channel is opened to simulate low ACh concentration, and the channel is closed for REM simulations to simulate high ACh concentration.^63,64^ By incorporating this additional conductance, a primary effect of the most abundant muscarinic ACh receptor in the brain (the M1 receptor) is included in the model. *In vivo* and in the model, activation of this potassium leak current alone reduces cell excitability and increases network synchronicity.^65-69^ In the model, modulation of this M-type current also changes the excitatory-inhibitory balance of the network, supporting higher average interneuron rates (**Figure S10A**).

**Figure 9.**
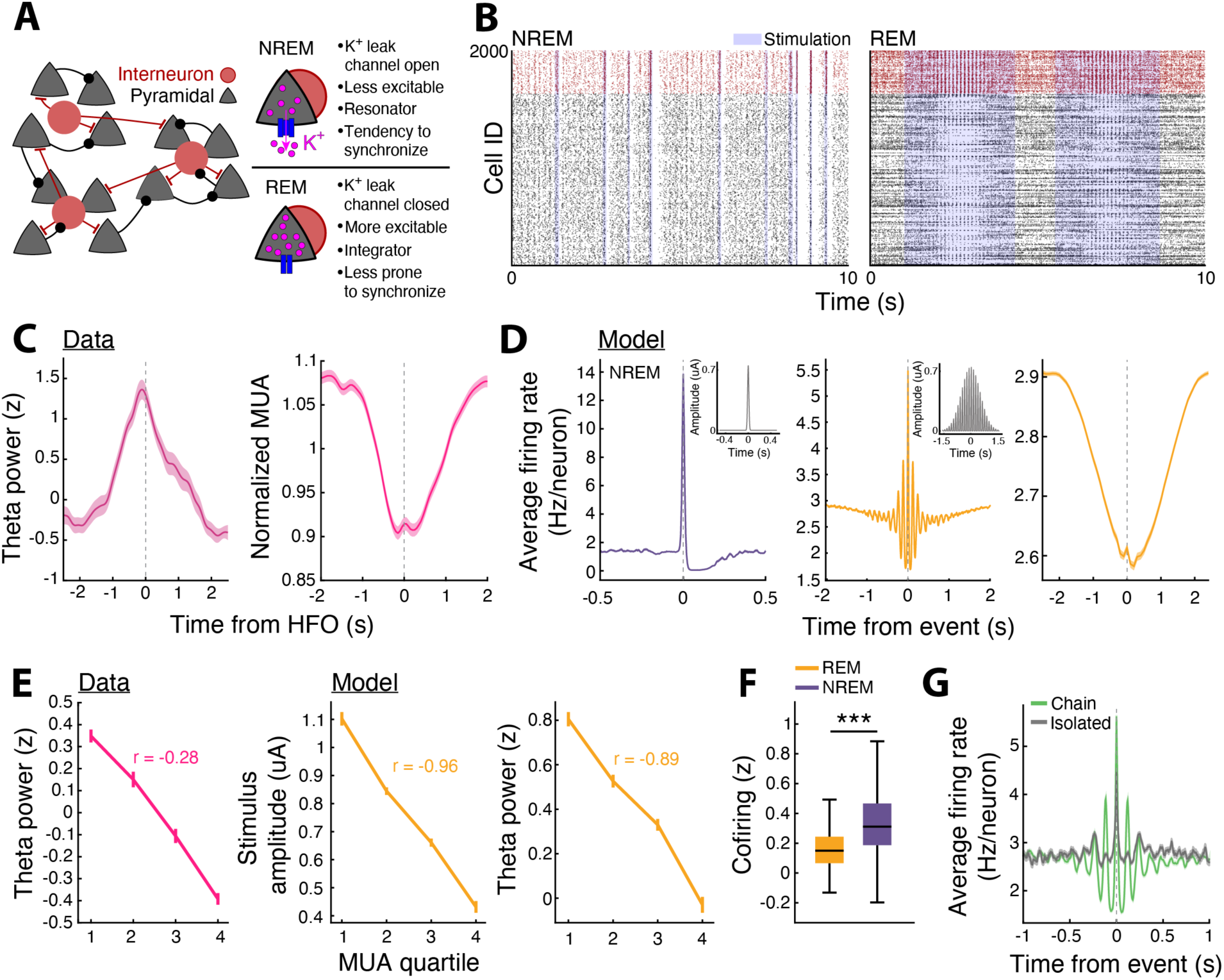
A simple network model incorporating acetylcholine recreates REM results. (**A**) Model schematic. Network of Hodgkin-Huxley (HH) pyramidal and interneurons with an added slow potassium leak conductance. This channel is open in NREM, introducing an adaptation current that lowers cell excitability and bestows resonator properties to all neurons, altering their phase response curves such that connected cells are better able to synchronize spiking. In REM the channel is closed to simulate high ACh concentration, eliminating the adaptation current. (**B**) Example raster plots of network activity in NREM and REM. Spikes from 400 interneurons shown in red, spikes from 1600 pyramidal neurons shown in black. Shaded regions indicate periods of applied input current (see Methods). (**C**) Experimental data. (Left) Z-scored theta power surrounding REM PFC HFOs shows an average 3 second increase in theta power. (Right) Normalized MUA aligned to REM PFC HFOs using 100 ms bins shows suppression. (**D**) Modeling MUA results. (Left) NREM average network firing rate aligned to MUA peaks during stimulus periods, with inset depicting the average stimulus profile. (Center) same as (Left), but for REM. (Right) REM 100 ms binned firing rate aligned to MUA peaks within stimulus periods (i.e., a smoothened version of the Center panel) shows suppression, similar to experimental data. (**E**) (Left) Experimental data showing the negative correlation between peak theta power during REM HFOs and average MUA during HFOs. (Center) Peak amplitude of applied stimulus vs. average MUA quartiles during stimulus period in the model. (Right) Same as (Center), but for z-score of average theta power (measured from network MUA) during stimulus period. (**F**) Z-scored cofiring within MUA peaks in stimulus periods in the model. REM peak duration 37 ± 9 ms (mean ± std), NREM peak duration 35 ± 9 ms. (**G**) Average firing rate surrounding MUA peaks within stimulus periods in the model split between isolated and chained MUA peaks. Isolated peaks defined as having no other peak within a 200 ms.

While the model does not possess the complexity to replicate the same HFO-range spiking frequencies, we found that by applying a stimulus input to a randomly chosen subset of cells of the network, we could recreate some essential characteristics of network activity patterns during NREM and REM cortical events. A brief moderately strong input during NREM, representing input during NREM ripples, was sufficient to induce a larger burst of synchronous spiking in the network, causing a large peak in MUA (**Figure 9B,D**, left). In order to recreate the MUA surrounding REM PFC HFOs, we found that a theta-modulated gaussian input with a similar profile to the theta power observed around PFC REM HFOs (representing activity during phasic theta bursts in REM) was able to capture oscillatory and inhibitory features of the MUA (**Figure 9C,D**). We applied stimulus inputs of a variety of peak amplitudes to the network, and found that for temporally brief, discrete stimuli in NREM, stronger stimulation induced greater MUA (**Figure S10D**). Conversely, the strength of theta-oscillating input (and associated theta power as measured in the model’s MUA) during REM correlated negatively with the MUA level during stimulation **(Figure 9E**). We found a similar negative relation in the experimental data between theta power surrounding REM HFO times and MUA level, suggesting that increased theta power in PFC is associated with stronger external input.

The cofiring analysis done on experimental spiking data during REM and NREM events (**Figure 4E)** was implemented for the spikes of a matched number of randomly chosen model neurons during REM and NREM stimulation periods, showing greater cofiring during NREM similar to the experimental data (**Figure 9F**). In addition, we divided peaks in MUA during REM stimulation periods in the same manner used to separate isolated and chained REM HFOs in the data (minimum 200 ms between MUA peaks, **Figure 3**). Similar to the data, we observed greater theta modulation in the MUA surrounding chained peaks compared to isolated peaks, as well as a larger dip in overall MUA (**Figure 9G**), with the difference especially clear in the pyramidal population of the model (**Figure S10H**). Overall, these results indicate an ability of the model to capture essential features of PFC activity in NREM and REM sleep, and suggest that REM HFOs reflect transient periods of elevated theta input to PFC neurons, leading to characteristic activity patterns in the network due to high cholinergic tone.

### Stimulation in NREM induces widespread network activation compared to stimulation in REM

Analyzing stimulation periods in the model’s NREM and REM states revealed intriguing differences in network activity (**Figure 10A**). While stimulus periods in REM tended to induce only a slight increase in the total fraction of cells spiking at any given time, stimulation in NREM induced a much wider spread of coactivity throughout the network (**Figure 10B**). We showed that the wider spread of coactivity depended on the decreased ACh concentration characterizing the NREM state in our model, and not the difference between REM- and NREM-inputs, because application of the brief NREM stimulus during REM did not produce the NREM level of peak coactivity (**Figure S10G**). Analysis of the percentage of PFC cells spiking in periods surrounding NREM and REM events in the model also showed this increased spread of coactivity in NREM, in agreement with the experimental results (**Figures 4A,E**, **6A,B,** and **10B**). However, it is unclear whether the widespread coactivity in NREM is a result of propagation of spiking due to recurrent excitation, or simply greater efficacy by the stimulus in directly inducing spikes in stimulated neurons during NREM. To investigate this, we analyzed model results by separating neurons within each stimulus event into stimulated and non-stimulated fractions. When stimulating in NREM, a widespread barrage of spiking from stimulated cells was followed by a smaller increase in the percentage of non-stimulated cell firing among both pyramidal and interneuron populations (**Figure 10C**). In contrast, during REM stimuli, the number of active stimulated neurons increased, but the effect on non-stimulated neurons was suppressive (**Figure 10D**). Again, swapping stimulus profiles showed that the cholinergic tone in the model plays a large role in creating this difference, especially as the brief NREM-like stimulus applied during REM produces no increase in percentage of coactive non-stimulated neurons (**Figure S10I,J**). These findings nicely align with experiments showing that ACh both suppresses the spread of cortical recurrent excitation and drives inhibition.^70-73^ Our results suggest that the ACh level regulates the degree to which inputs can induce widespread synchronous network activity, leading to the distinct activity patterns in NREM and REM. Low ACh concentration and the associated reduced inhibitory tone in NREM leads to a rapid spread of activity to many non-stimulated cells upon transient excitatory input to a subset of the network. Such a result suggests that the level of ACh regulates the fraction of cells in a circuit able to respond to a stimulus, accounting for the observation of synchronous bursts of activity among a large portion of cells during NREM (low ACh and low inhibition) whereas coactivity is restricted to smaller subsets of cells in the network during REM (high ACh and high inhibition).

**Figure 10.**
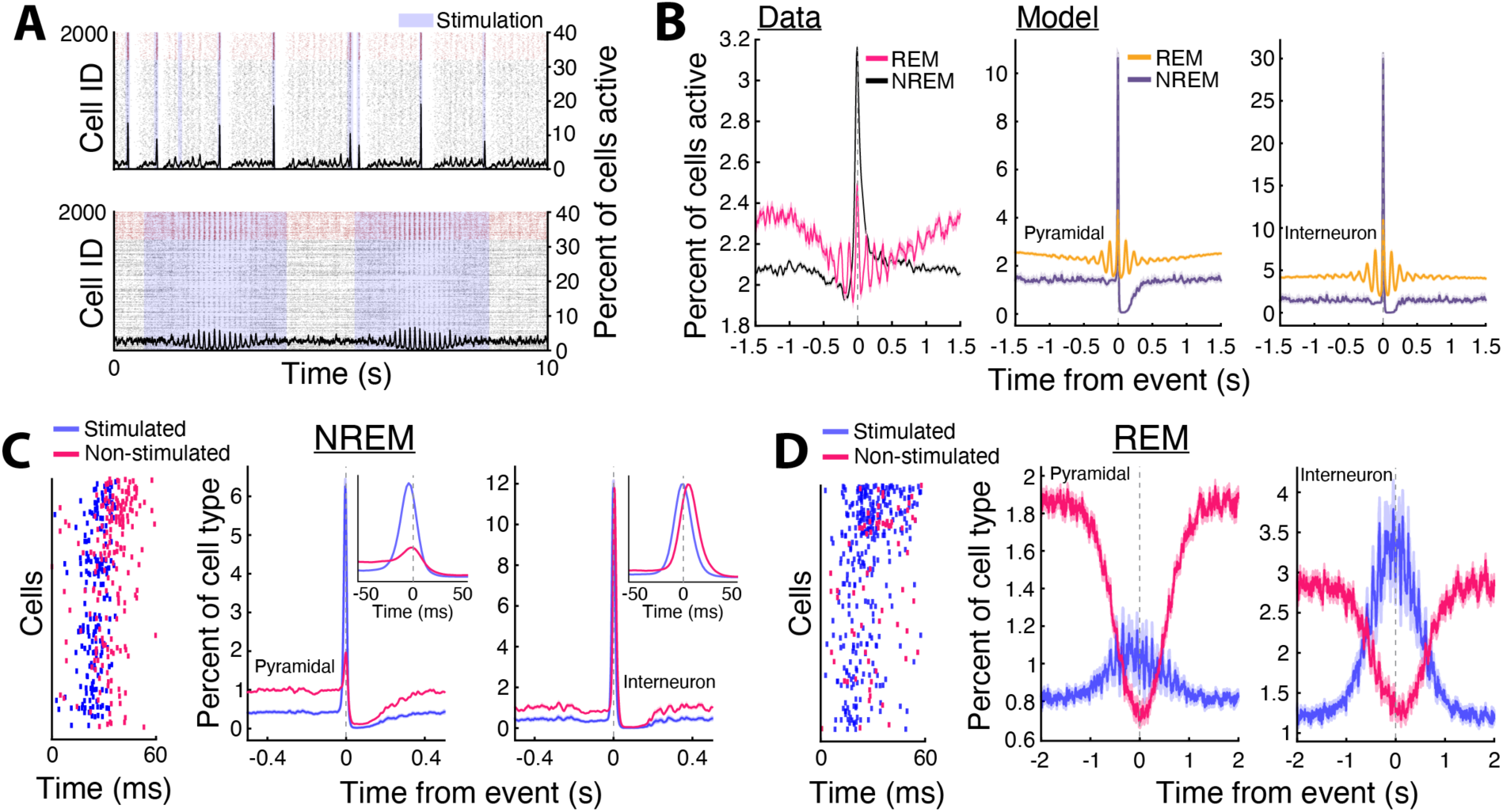
Stimulation in NREM induces widespread coactivity compared to stimulation in REM. (**A**) Model raster plots during example NREM (Top) and REM (Bottom) periods with black trace indicating the percent of all pyramidal cells with at least one spike in 10 ms bins. (**B**) The same percent active measure (not distinguishing by cell type) calculated on experimental data aligned to NREM and REM events (Left), and in the model aligned to MUA peaks in NREM and REM as a fraction of all pyramidal neurons (Center). Similar plot for all interneurons in the model (Right). (**C**) Model NREM activity split by cells directly receiving stimulus input (blue) and receiving no external stimulus (pink). Raster of a single example population burst due to stimulation in NREM (Left). Percent of cells with at least one spike in 10 ms bins (as a percent of all pyramidal cells in the network) aligned to stimulus midpoints, the time of peak input amplitude (Center). Similar plot for interneurons (Right). (**D**) Same as (**C**), but for REM. Note that non-stimulated cell population (pink) comprise a greater percent of total active neurons outside stimulation times (average stimulation time of 3 sec centered at time of event), as there are more neurons in the non-stimulated group (only 30% of neurons are stimulated per stimulus event).

## DISCUSSION

One of the key challenges in understanding the role of REM sleep in memory consolidation has been disentangling the interregional neurophysiological processes involved in memory-related processing. Previous studies have primarily focused on the role of theta oscillations in REM sleep as a brain-wide pattern that coordinates activity across various brain regions,^74^ due to their similarity with patterns observed during wakeful behavior. While much attention has been focused on investigating theta oscillations due to their prominence in the hippocampus during REM sleep, the precise dynamics that drive the coordination of multiple brain areas to potentially support systems memory consolidation have been elusive. Furthermore, the prominence of oscillatory phenomena in NREM sleep—including slow oscillations, spindles, cortical ripples, and CA1 SWRs—has enabled systematic dissection of their spatiotemporal dynamics and their relationship to hippocampal-cortical reactivation that supports memory consolidation,^11-14^ leaving REM sleep dynamics poorly understood in comparison.

Here, we show that phasic activity bursts in PFC during REM theta coincide with chains of local HFOs and elevated gamma power. These chains are associated with increased measures of oscillatory coupling between PFC and CA1, and show characteristic theta-modulated spiking activity patterns on background of local suppression. REM PFC HFO chains organize sparse, sequential reactivation in PFC over extended time periods, and recruit a sub-population of CA1 neurons that exhibit differential regulation of activity over the course of sleep. This contrasts with NREM sleep, where independent PFC ripples are associated with relatively brief, synchronous bursts of local reactivation along with suppression of CA1 activity. We also show that CA1 co-activation during REM PFC HFOs is related to the CA1 activity suppression seen during NREM PFC ripples, with CA1 neurons that are highly coactive with PFC during REM HFO chains being strongly suppressed during NREM ripples, suggesting a dual sleep-stage role of PFC ripples and HFOs in regulating excitability in hippocampal circuits. Thus, these results uncover a novel, high-frequency pattern of PFC activity in REM that can potentially bridge the gap between structured PFC and CA1 firing patterns in NREM-REM stages and changes in hippocampal plasticity during sleep.

In contrast to PFC ripples in NREM sleep, which are involved in suppressing CA1 activity as a possible mechanism for tagging specific memory traces for consolidation,^28^ here we found that REM cortical HFOs engage specific populations of CA1 neurons to potentially regulate activity homeostasis. Of particular interest regarding this phenomenon of activity regulation is a recent study examining the differential roles of NREM and REM sleep in reorganization of drifting hippocampal assemblies over the course of sleep.^75^ This study focused solely on hippocampal assemblies, and the results are in line with our finding of the potential roles of PFC ripples and HFOs in NREM and REM sleep, respectively. We have previously shown that independent NREM PFC ripples that are dissociated from hippocampal SWRs broadly suppress CA1 activity, and that this suppression is related to the strength of CA1 reactivation during SWRs and CA1 assembly reinstatement during subsequent behavior.^28^ This bidirectional modulation of hippocampal representations during PFC ripples and CA1 SWRs may underlie assembly reorganization during NREM sleep. Relatedly, we report here that CA1 activity during REM PFC HFOs is associated with differential changes in excitability in a subset of hippocampal neurons during sleep, which may, in part, contribute to the stabilization of assemblies in REM sleep and thus drive the sleep-stage dependent reorganization of hippocampal assemblies. This finding is in line with REM sleep’s role in recalibration of neural activity in hippocampal-cortical circuits.^5^ We speculate that PFC REM HFO chains enable selective linkages between the sparsely reactivated PFC ensembles in coordination with a subset of CA1 neurons.

Cortical high-frequency oscillations (∼120-160 Hz) in REM have been reported before, especially in parietal cortex, although they have been variously termed fast gamma or HFOs (high-frequency oscillations),^19,21,22,31^ with previous reports of strong coupling with theta oscillations in REM.^31^ HFOs have also been reported in parietal cortex and hippocampus during active waking behavior and REM, and it has been argued that HFOs in hippocampus are distinct from hippocampal SWRs.^22^ We detected HFOs in REM sleep using a similar frequency band (150-250 Hz) as cortical NREM ripples. These high-frequency events in NREM are denoted as “ripples”, a term used by previous studies for NREM high-frequency events in cortex,^23,25,27-29,76^ although this is likely a separate phenomenon from hippocampal SWRs. PFC NREM ripples that occur independently of hippocampal SWRs are associated with temporally synchronized local PFC reactivation and suppression of hippocampal activity and reactivation,^28^ while in REM, we show here that they present as PFC HFO chains during theta-modulated activity, associated with sparse, temporally extended local reactivation in the backdrop of suppression, along with co-activation of a hippocampal-subpopulation. These structured cortical sequences seen during HFO chains are gated by theta oscillation bursts in REM sleep, with higher rate of occurrence in phasic REM than tonic REM substages, and are reminiscent of hippocampal theta sequences,^77^ but may have different underlying mechanisms. We examined PFC HFO events during behavior on the W-maze, but did not find similar spiking activity patterns as REM HFO events (**Figure S11**), confirming that this is a REM-exclusive phenomenon.

Based on previous studies, external drive can lead to the generation of PFC ripples and synchronized activity in NREM, whereas high acetylcholine (ACh) and high inhibitory tone in REM can lead to local suppression in response to external drive.^73,78,79^ Notably, a recent computational study demonstrated that differential cholinergic modulation, together with the sequential architecture of sleep (NREM to REM transition), can support memory consolidation through the complementary shaping of ensemble activity during NREM and REM sleep.^64^ We show here that a simple network model that incorporates the effect of differential ACh tone on excitatory and inhibitory neurons in REM vs. NREM states is able to reproduce the observed experimental activity patterns in response to stimulation, akin to phasic theta bursts during REM and isolated, brief inputs in NREM. It is likely that PFC NREM ripples and REM HFOs are generated locally and modulated by external inputs. We speculate that theta-modulated input to PFC during REM is driven by hippocampal inputs, but it is also possible that theta is driven by inputs arising from regions other than the hippocampal network, a possibility which will require further experimental investigation.

We observed that isolated REM HFOs in PFC were more prevalent than chains, especially in tonic REM, potentially reflecting the need for coordinated activation of distributed circuits to elicit HFO-associated phasic events. One possible mechanism driving this phenomenon could involve PGO waves, which are prominent during phasic REM and are known to play a role in modulating hippocampal plasticity during REM sleep.^80^ These waves are also coordinated with both SWRs in NREM and theta states in REM,^80,81^ suggesting that PGO-associated amplification of hippocampal theta oscillations could drive activity above a certain threshold for HFO generation in PFC. Additionally, ACh may also play a role in altering the threshold for HFO generation in PFC in conjunction with PGO coordination. It has been reported that there is coordinated release of ACh in PFC and dorsal hippocampus, potentially elevating theta coherence between the two areas.^82^ Indeed, a number of studies have shown that ACh boosts theta oscillations.^83^ Furthermore, ACh induces decorrelation in cortical circuits, which improves cortical representations of relevant stimuli,^84^ and our finding that REM HFOs are associated with an overall decreases in PFC population activity, reactivation strength and cofiring, in stark contrast to NREM ripples, suggests a role for REM HFOs in amplifying local cortical signal to noise to enable linking of cortically encoded information with specific populations of hippocampal neurons. Lastly, cholinergic input to the hippocampus also suppresses SWR generation,^85,86^ which could facilitate the sparse CA1 reactivation during REM sleep for selective changes in excitability and stabilization of salient representations in coordination with cortical networks for subsequent expression during behavior. Future experimental and network modeling studies can investigate how neuromodulatory mechanisms and NREM and REM state-specific cortical activity patterns act in concert to support learning and memory consolidation.

Taken together, these results establish a new cortical LFP pattern in REM sleep (PFC REM HFO chains) associated with a distinctive neuronal population activity signature. The findings are aligned with REM sleep’s role in memory consolidation and provide a neural basis for the cortical-hippocampal modifications that occur during sleep to support activity reorganization and mnemonic processing. The complex interplay between oscillations of varying frequencies revealed here further highlight the importance of segregating sleep into multiple states to uncover how systems memory consolidation can be supported through the multiplexing of distinct phases, both during NREM and REM sleep. The interareal dynamics associated with PFC REM HFOs involve both bottom-up and top-down processes operating in precise temporal coordination, with theta-mediated communication leading to PFC HFO-associated reactivation and modifications in hippocampal excitability. These findings, in conjunction with PFC-mediated suppression of CA1 during NREM sleep, suggests a dual sleep stage processing of mnemonic representations that have implications for understanding the circuit mechanisms underlying systems memory consolidation, and can reveal how neural processes in sleep go awry in disease states.

## LIMITATIONS

While the present results establish a potential role for REM sleep PFC HFOs, several limitations of our study are acknowledged here. First, we were unable to directly link REM HFO specific reactivation to learning. Spatial memory tasks in rodents similar to our study have revealed causal links of NREM reactivation driven by CA1 SWRs to learning and memory consolidation,^87-90^ but reports of REM reactivation are rare.^37^ Investigation of the role of REM reactivation in learning and memory consolidation will need utilization of tasks known to require both REM and NREM states, such as schema-based inference or emotional memories.^38,91-93^ Second, behavioral task learning and interleaved sleep sessions were conducted in a few hours within a single day, which did not allow us to densely sample the circadian cycle. Third, we did not record eye movements or ponto-geniculo-occipital (PGO) waves, both of which would have allowed for more accurate segregation of tonic and phasic REM sleep states.^94^ Although we observed a bias for isolated and chain HFOs to occur in putative tonic and phasic REM substates, respectively, the scarcity of putative phasic REM bouts made the direct comparison based on substage difficult. Finally, although our model predicts that distinct cell-type activity profiles shape REM sleep HFO dynamics, we did not record a sufficient number of interneurons to test these predictions directly. Future studies using appropriate behavioral tasks, longitudinal sleep recordings, and cell-type specific opto-tagging will be able to resolve these limitations and further clarify the roles of high-frequency oscillations in REM sleep.

## ACKNOWLEDGEMENTS

This work was supported by the National Institutes of Mental Health (R01MH112661 to SPJ).

## AUTHOR CONTRIBUTIONS

J.D.S. and S.P.J. conceived the study. J.D.S. performed the experiments. J.D.S. processed and analyzed the data. M.S. conceptualized the model and P.M. advised on model implementation. J.D.S., M.S., and S.P.J. wrote the manuscript. S.P.J. provided supervision and funding for all aspects of the study.

## DECLARATION OF INTERESTS

The authors declare no competing interests.

## SUPPLEMENTAL INFORMATION

Figures S1–S11 and Table S1

## RESOURCES AVAILABILITY

### Materials availability

This study did not generate any new unique reagents.

### Data and code availability

- Data underlying these results are available in the NWB (Neurodata Without Borders) format on DANDI Archive (ID#000978).
- Code to replicate these results will be available on our lab GitHub (http://github.com/JadhavLab).
- Any additional information required to reanalyze the data reported in this paper is available from the lead contact upon request.

## EXPERIMENTAL MODEL AND SUBJECT DETAILS

All experimental procedures were approved by the Institutional Animal Care and Use Committee at Brandeis University and conformed to US National Institutes of Health guidelines. Ten adult male Long-Evans rats (450-600 g, 4-6 months; RRID: RGD_2308852) were used in this study. Animals were individually housed and kept on a 12-hr regular light/dark cycle.

## METHOD DETAILS

### Surgical implant and electrophysiology

Surgical implantation procedures were as previously described.^95^ Animals (n = 10) were implanted with a microdrive array containing 30, 32, or 64 independently moveable tetrodes targeting right dorsal hippocampal region CA1 (-3.6 mm AP and 2.2 mm ML) and right PFC (+3.0 mm AP and 0.7 mm ML). For 64 tetrode recordings (n = 1 animal), CA1 and PFC were targeted bilaterally. Tetrodes were split equally between PFC and CA1 (15, 16, or 32 tetrodes in each region). On the days following surgery, hippocampal tetrodes were gradually advanced to the desired depths using characteristic EEG patterns (sharp wave polarity, theta modulation) and neural firing patterns as previously described.^95^ A CA1 tetrode in corpus callosum served as hippocampal reference, and another tetrode in overlying cortical regions with no spiking signal served as prefrontal reference. A ground (GND) screw installed in the skull overlying cerebellum also served as a reference. All spiking activity, gamma, and ripple/HFO-filtered LFPs (gamma: 40-100 Hz; ripple/HFO: 150-250 Hz; see below) were recorded relative to a local reference tetrode. For detection of slower frequency components (i.e., theta and delta), LFP was recorded with respect to the GND screw. Electrodes were not moved at least 4 hours before and during the recording day.

Data were collected using a SpikeGadgets data acquisition system (SpikeGadgets LLC). Spike data were sampled at 30 kHz and bandpass filtered between 600 Hz and 6 kHz. LFPs were sampled at 1.5 kHz and bandpass filtered between 0.1 Hz and 400 Hz. The animal’s position was recorded with an overhead color CCD camera (30 fps) and tracked by color LEDs affixed to the headstage. Additionally, the animals’ speed was calculated based on predetermined cm/pixel values and the position displacement between frames captured at 30 fps.

### Behavior

Ten animals were trained on a novel W-maze in a single day, as previously described.^28,40,95^. This single-day W-maze alternation learning task is SWR-dependent, and requires hippocampal-prefrontal interactions.^39,96,97^ Briefly, following recovery from surgical implantation (∼7–10 days), animals were food-deprived and again retrained on a linear track for 2-3 days before exposure to the W-Maze task. During the recording day, animals were introduced to the novel W-maze (∼80 × 80 cm with ∼7 cm wide tracks) for the first time and learned the task rules over eight behavioral sessions during the animals’ light phase between the hours of 9 AM and 6PM. Each behavioral session lasted 15–20 min and was interleaved with 20–40 min rest sessions in the sleep box for a total of 17 separate sessions (8 W-maze and 9 sleep sessions). The total recording duration spanned ∼6-7 hours within a single day. On the W-maze, animals were rewarded for performing a continuous spatial alternation. The task rules are as follows: returning to the center well after visits to either side well (left or right well; inbound trajectories) and choosing the opposite side well from the previously visited side well when at the center well (outbound trajectories). Rewards were automatically delivered to the animal through custom 3D printed reward wells upon nose poke.

### Histological verification of recording sites

At the completion of each experiment, electrolytic lesions were made by sending current (30 uA) down 3 electrodes per tetrode for 3-4 seconds each. 24 hours later, animals were sacrificed by intraperitoneal injection of Euthasol and subsequent intracardial perfusion with 0.9% saline followed by 4% formaldehyde. The skull, with the attached drive, was submerged in 4% formaldehyde for 24 hours before tetrode retraction and brain extraction. Brains were stored in a 4% formaldehyde and 30% sucrose solution for at least 1 week before slicing with a freezing microtome (Leica) at 50 uM. Slices were Nissl stained and imaged at 4x on a brightfield microscope (Keyence) and stitched together to generate composite images for lesion localization.

### Spike sorting

Single units were sorted using a custom manual clustering program (MatClust, M. P. Karlsson) and were distinguished based on spike width, peak and trough amplitude, and principal components. Cluster quality was measured using isolation distance,^98^ and only well isolated neurons were included for analysis.

### Sleep state identification

To detect bouts of NREM and REM sleep, animals’ head speed and CA1 LFPs were used as previously described.^15,16^ During sleep box sessions, times of awake activity and immobility were defined as periods where head speed was greater or less than 4 cm/s, respectively. Candidate NREM sleep bouts were identified as periods where head speed remained <4 cm/s after an immobility period of at least 60 s. NREM and REM sleep were further identified and separated by using a theta to delta ratio metric. Briefly, CA1 LFP (referenced to GND) for each tetrode was bandpass filtered (6-12 Hz for theta; 1-4 Hz for delta) and averaged, and the ratio of theta to delta (TD ratio) power was calculated.

Periods that exceeded a set threshold (mean + 1 SD) for >10 s were categorized as REM bouts within candidate NREM bouts.^99,100^ Additionally, candidate REM bouts were visually inspected to further confirm the transition into REM sleep. For comparison of velocity and TD ratio, the average velocity and TD ratio in each epoch for NREM and REM sleep were computed and compared. Additionally, only epochs with at least 30 s of REM sleep were included for analysis. Unless otherwise stated, all analyses were restricted to the 36 (of 90) sleep epochs that satisfied the inclusion criteria.

### Intracranial electromyogram (EMG)

To estimate EMG from the neck, jaw, and face muscles from intracranial LFP signals, a previously established method was used.^35,101^ Briefly, we extracted the 300-600 Hz filtered signal (Butterworth filter at 300 – 600 Hz; filter shoulders from 275 – 625 Hz) and calculated the pairwise Pearson correlations in each 500 ms bin between pairs of electrodes during the entire sleep epoch. Since we used tetrode recordings and cannot determine the exact separation of electrode sites, we elected to calculate the correlations between the reference tetrode in corpus callosum and CA1 ripple detection tetrodes within the dorsal CA1 pyramidal cell layer (∼500 µm separation between corpus callosum and the dorsal CA1 pyramidal cell layer), thus maximizing the separation between the electrode sites. Previous studies have used electrode channels on probes at least two shanks apart (∼400 µm). The average of all reference/CA1 layer tetrode pairs was calculated and compared for wake and REM sleep. For the distributions of intracranial EMG values in **Figure S1D**, each 500 ms bin correlation value was reported, and for the comparison between wake and REM sleep, the averages during each wake and REM bout were calculated and compared. A strong relationship between intramuscular EMG and intracranial EMG has been reported.^101^

### LFP event detection

PFC ripple/HFO detection: High-frequency events were detected during immobility periods (<4 cm/s) as previously described.^16^ Tetrodes with recorded cells in CA1 and PFC were used as candidate electrodes for event detection as well as downstream LFP analysis. Each LFP from candidate electrodes were bandpass filtered in the ripple/HFO band (150-250 Hz), and the envelope power was calculated using a Hilbert transform and subsequently smoothed with a Gaussian kernel (σ = 6). Events were detected as contiguous periods where the ripple/HFO power exceeded 3 SD of the mean on at least one tetrode. A minimum event duration of 15 ms was implemented to exclude spurious noise in the LFP signal. Events were further refined as periods around the detected event that remained above the mean (start and end times). The amplitude of each event was reported as the number of SD above the mean.^40^ To estimate the frequency of each event, the power in each frequency bin was calculated around each event (±500 ms) using multi-taper time-frequency analysis (Chronux toolbox; http://chronux.org/)^102^ and was z-scored relative to a baseline spectrogram. The frequency of each event was defined as the frequency with the highest power in the ripple/HFO band (100-250 Hz) between the start and end time of each event.^103^ Furthermore, HFO chains were defined as 2 or more events that occurred <200 ms apart. Average amplitude plots for each animal were generated by aligning the peaks of each event and averaging.

PFC gamma detection: Bouts of high gamma power were detected as above after bandpass filtering in the gamma band (40-100 Hz).

High theta power detection: Each LFP from candidate PFC tetrodes were used to delineate periods of high theta power during REM sleep. LFPs were bandpass filtered with a theta band filter (6-12 Hz), the envelope power was calculated using a Hilbert transform, averaged across tetrodes, and periods that exceeded 4 SD of the mean during REM sleep were designated as bouts of high theta power. Only high-power periods exceeding 900 ms were included as high theta power periods.

PFC spindle detection: Spindle events were detected as previously described.^45^ The mean LFP across all candidate PFC electrodes was calculated and the resulting signal was bandpass filtered between 10-16 Hz with a zero-phase shifted, third-order Butterworth filter. The envelope power was calculated using a Hilbert transform and subsequently smoothed with a Gaussian kernel (α = 2.5). Events that exceeded 2.5 SD of the mean were extracted, and events separated by <300 ms were concatenated. Spindle events were restricted to NREM sleep before analysis. Spindle trains were defined as 2 or more spindle events that occurred ≤2.78 s apart as previously established.^45^

### Spectral analysis

Wavelet spectral analyses were utilized to calculate the ripple/HFO aligned power spectra in PFC (2–350 Hz, Morlet wavelets). For each epoch, the PFC tetrode with the greatest number of detected PFC ripples/HFOs was used. The power at each level of the wavelet transform was calculated around each event (±500 ms) and was z-scored relative to a baseline spectrogram calculated for the entire epoch. For each animal, spectrograms from all candidate sleep epochs were averaged.

### Event autocorrelation

To calculate the autocorrelation of ripples/HFOs to assess the structure of event occurrence, identical event time vectors were lagged (max time lag 1 second, 10 ms time bins), and the correlations were calculated. The results were smoothed with a Gaussian kernel (σ = 3) and averaged across all animals and epochs. Similarly, the cross-correlation between gamma and HFO time vectors was calculated by using the gamma times as the reference vector. Thus, positive and negative lag values indicate that the HFOs tended to occur before and after gamma events, respectively.

### Event aligned multi-unit activity

To characterize population activity aligned to LFP events, spike trains of each neuron were binned into 5 ms time windows (Figures 2A and **4F**). Multi-unit activity was aligned to PFC ripple/HFO and gamma events and was normalized by the mean population firing rate during either NREM or REM sleep, depending on the events assessed. Subsequently, the power spectral density of the ripple/HFO aligned multi-unit activity was assessed (frequency limits, 2-12 Hz; MATLAB, *pwelch*).

### Control for periods of high theta power

To determine whether theta-modulated population activity in PFC can be largely explained by periods of high theta power, instead of association with HFO events, we extracted surrogate timestamps to align PFC activity to (**Figures S5K-N**). Briefly, for each epoch, the preferred theta phase bin of HFO events was calculated from the PFC tetrode with the highest average theta power and used to randomly select surrogate times to align PFC multiunit activity to. Each preferred phase bin at least 250 ms away from a detected HFO was considered a candidate event. Then, times were randomly chosen, and the number of events selected matched the number of HFO events in that epoch. Theta power at each surrogate time point as well as HFO event was extracted and z-scored to control for any inter-animal variability in theta power. Lastly, the event aligned multiunit activity was split into quartiles based on theta power and compared.

### Theta inter-peak intervals during bouts of high and low theta power

Since previous studies have segregated putative tonic and phasic substages of REM sleep based on CA1 theta frequency,^15^ we calculated the inter-peak intervals of the theta oscillation to determine whether bouts of low and high theta power may correspond to putative tonic and phasic substages, respectively (**Figure S3B**). We bandpass filtered and averaged the LFPs from candidate CA1 tetrodes between 5-12 Hz, and the peaks were extracted by detecting positive-to-negative zero crossings of the derivative of the filtered, averaged signal. Lastly, the inter-peak intervals were calculated within low and high theta power periods for each epoch and compared. Note that we defined low and high theta periods based on PFC electrodes, whereas we calculated inter-peak intervals based on CA1 electrodes.

### Cross-frequency phase-amplitude coupling

Phase-amplitude coupling was assessed for a range of frequency combinations using a previously established approach.^104^ Since the rate of REM HFOs was highest during bouts of high theta power, we focused on these periods for assessing phase-amplitude coupling. For each epoch, the tetrode with the highest HFO power was chosen, and an averaged comodulogram was generated across all animals and epochs for the phase and amplitude frequency ranges from 2-12 Hz and 40-250 Hz, respectively.

Comodulogram: Briefly, the phases were extracted from the Hilbert transform of the raw LFP of the candidate electrode and binned into 18 bins (intervals of 20°), and the envelope power of the amplitude modulated frequency band was calculated by taking the absolute value of the Hilbert transform. Then, the amplitude of each frequency from 40-250 Hz was calculated in each phase bin from 2-12 Hz and visualized (Figure 3B).

Modulation index: To quantify the depth of amplitude modulation by the phase frequencies, gamma (40-100 Hz) and HFO (150-250 Hz) signals were separately processed, and the mean gamma or HFO amplitude in each phase bin was calculated. The resulting values were used to calculate the modulation index,^104^ which specifies the strength of phase modulation. Additionally, the phase bin with the maximum amplitude was reported as the preferred phase. For comparison, the phase and amplitude vectors were circularly shifted independently, and the modulation index was calculated.

### Theta nesting

To further investigate the coupling of theta, gamma, and HFOs, in PFC during REM sleep, we examined theta nesting^105,106^ in two different frequency bands (40-100 Hz for gamma and 150-250 Hz for HFO) relative to the theta oscillation (Figures 2C and **S4E,F**). Since we are evaluating the gamma and HFO coupling specifically during these transient HFO events, we restricted our analysis to high theta periods since the rate of REM HFOs was highest during these REM bouts (**Figure S3D**). For each PFC electrode, the theta phase bin for which gamma power was maximal was identified. Then, each instance of this phase bin during high theta periods in REM sleep was detected and the index of the highest gamma power within each preferred phase bin was identified. Only electrodes where there were at least 20 instances of the preferred phase were included. Lastly, the averaged waveform was calculated by averaging the unfiltered LFP centered on every instance of maximal gamma power in the preferred bin. The above procedure was performed separately for HFO band activity. For comparison between the frequency of averaged gamma and HFO waveforms and the preferred phase of theta coupling, electrodes that exhibited at least 5 local maxima within a 100 ms window surrounding the preferred phase were analyzed.

To further validate coupling of gamma and HFO to theta, surrogate waveforms were generated by randomly selecting a different phase bin for each theta cycle for alignment of gamma or HFO power. Then, peak-to-trough amplitudes were calculated from the average waveform within the 100 ms window centered on peak power, averaged, and compared between the preferred phase bin aligned data and the random phase bin surrogate.

### Phase-locking

Phase-locking during active behavior on the W-Track and during REM sleep was quantified using previously developed methods.^107^ The CA1 or PFC tetrode with the greatest ripple/HFO power was filtered in the theta (6-12 Hz), gamma (40-100 Hz), or ripple/HFO (150-250 Hz) range and was used to extract phase information. For HFO and gamma phase locking of PFC cells, each spike that occurred across all extracted events were assigned a phase, and the distributions of the peak phases of all cells were compared between gamma and HFO. During run sessions (for ‘REM theta phase shifting’ analysis below, **Figures S9H,I**), spikes were restricted to CA1 theta periods where the animals’ speed was >5 cm/s and were assigned a phase between 0-360°. Only neurons with at least 10 spikes during theta periods were included in this analysis. The Rayleigh test was used to determine whether the phase distribution of each cell deviated from circular uniformity (p < 0.05 criterion). Additionally, for phase locking to gamma and HFOs, the preferred phases of each cell was calculated separately for gamma or HFOs. Thus, a cell could be significantly phase locked to both gamma and HFOs. To test for phase preference differences, the distributions were compared using a nonparametric Watson’s U^2^ test. Subsequently, for each theta-modulated cell, phase locking strength to the theta oscillation was measured using pairwise phase consistency (PPC).^108^ PPC was calculated by taking the dot product of the angular difference between each pair of phase values then averaging. Similar procedures were performed to determine the HFO and gamma phase preference of PFC cells.

Additionally, PPC was used to assess the strength of theta phase locking of HFO/ripple events in REM or NREM sleep. Only epochs with at least 20 events of each type were included for analysis.

### Theta coherence

Theta coherence between pairs of CA1 and PFC tetrodes was calculated during REM sleep periods centered on REM HFOs, and each frequency was normalized by the mean and SD of the baseline coherogram computed across the entire epoch (**Figures 3E,I**). Coherograms were averaged over CA1-PFC tetrode pairs. For quantification and comparison, theta coherence was measured as the peak coherence between 6-12 Hz in a ±1 s window around HFO events. For visualization of coherograms, the averaged coherograms across all animals and epochs were smoothed with a 2D Gaussian kernel (σ = 4). All coherence-based analyses were conducted using the Chronux toolbox.^102^ Only epochs with at least 20 events were included for quantification of peak theta coherence.

### Phase slope index

Since we observed elevated theta coherence between CA1 and PFC during chained events, we assessed the directionality of information flow at different frequency bands (0-50 Hz) centered on these events (±1 s) using the phase slope index (PSI, **Figure 3J**),^109^ which was calculated with the FieldTrip analysis toolbox.^110^ In practice, PSI is used to assess the consistency of phase lag relationships between two signals across different frequency bands and is a measure that is weighted by oscillatory coherence. We opted to use PSI to estimate the directional flow of information instead of other methods, such as Granger causality, since it has been demonstrated that PSI is less prone to false positives.^109^ Briefly, for each epoch, the PSI was calculated and averaged across each candidate CA1-PFC tetrode pair. Here, positive PSI values indicate that PFC theta leads CA1, while negative values indicate that CA1 leads PFC. Then the distribution of mean PSI values within 6-12 Hz was tested against 0 with a t-test. Only epochs with at least 5 chained events were included for analysis.

### Ripple/HFO aligned modulation

Single unit ripple/HFO modulation in CA1 or PFC was calculated as previously described (**Figures 4A,D**).^28,111^ This analysis was restricted to neurons that fired >50 spikes cumulatively across all peri-event windows (±2 s). For each candidate neuron, an averaged event-triggered (NREM or REM event) time histogram (event-PSTH) was calculated. For comparison, 1000 shuffled event-PSTHs were generated by circularly shifting the spikes around each ripple/HFO event by a random amount. To determine whether a neuron was significantly modulated, the squared difference between the real event-PSTH and the shuffled data in the -200 to +200 ms window surrounding event onset was calculated. If this real value exceeded 95% of the shuffled values, the neuron was categorized as significantly modulated. The modulation index for each cell was given by calculating the average difference between the activity in the response window and a background window -600 to -200 ms from event onset. Furthermore, excited (EXC) and inhibited (INH) neurons were separated based on whether the modulation in the event response window was positive (EXC) or negative (INH). Modulation timing distributions for EXC PFC neurons during NREM and REM PFC events were compared by binning (2 ms bins) the peak responses within the modulation window.

### Event type prediction

High-frequency event type (NREM or REM) in PFC was predicted by fitting a linear model (MATLAB, *fitclinear*, **Figure 4C**) to PFC spike count data. Since the NREM events largely outnumbered the REM events, the event counts were equalized prior to training by randomly excluding a subset of NREM events. The binary decoders were built using 10-fold cross validation, and prediction accuracy was assessed by comparing performance against models constructed with randomly permuted spike matrices (n = 1000). To account for the excluded events, this entire process was repeated 10 times for each animal and epoch.

### Ripple and HFO cofiring

To quantify ripple/HFO cofiring during events, a pairwise cofiring z-score was calculated as previously described (**Figures 4E** and **7B,C**).^48^ Briefly, this cofiring measure estimates how likely two cells will fire together during events and is normalized by the incidence of spiking independent of the other cell:

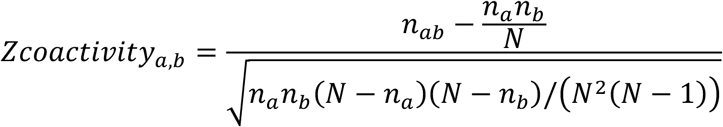

where *N* is the number of events, *n*_*a*_ and *n*_*b*_ are the number of events where cell *a* and cell *b* were active, respectively, and *n*_*ab*_ is the number of events where both cells were active. Here, cofiring assesses coincident activity between neuron pairs within the start and end times of events, and thus, no explicit temporal threshold was implemented. The same procedure was used to calculate cofiring between pairs in the model simulations.

### Detection of population suppression

To validate the suppression of PFC multiunit activity surrounding REM PFC HFOs, periods of decreased PFC activity were separately extracted, and the probability of REM HFOs surrounding these events was calculated (**Figures 4H** and **S6B,C**). First, multiunit activity was binned (5 ms bins), normalized by the average activity during REM sleep, and smoothed with a Gaussian kernel (σ = 3). The population activity was then z-scored, and periods that fell below 0 for 300-500 ms were extracted as population suppression events. Lastly, the probability of HFO occurrence surrounding these events was calculated in 100 ms bins. To obtain the 99 percent confidence intervals, suppression events were extracted from shuffled multi-unit data. Briefly, each cells’ spike count was circularly shifted independently, and events were detected. A total of 1000 shuffles were performed, and the probability of HFO occurrence surrounding these events was calculated for each shuffle.

### Relationship between inter-event-interval (IEI) and content of adjacent HFOs

To assess whether there was a relationship between IEI and population spike content between HFO events, the spiking activity of PFC neurons during HFOs was represented as ones and zeros for either active or inactive during all events (**Figures 5B** and **S7A**).

The Pearson correlation between population activity vectors of two adjacent HFOs was calculated, and the IEI was given by calculating the temporal difference between the centers of the HFO events. Then, the relationship between spike content and IEI was assessed using the Pearson correlation coefficient. To validate this relationship, we employed two other methods for quantifying similarity of activity across two events, the Jaccard index and cosine similarity. The Jaccard index, or the Jaccard similarity coefficient, was used to quantify the similarity between two sets. It is defined as the ratio of the size of the intersection of the two sets to the size of their union. The Jaccard index (J) is expressed as

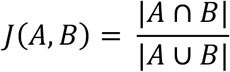

where A and B represent the two sets being compared (spike activity for HFO A vs HFO B), ∣A∩B∣ is the number of elements in the intersection of the sets, and |A∪B| is the number of elements in the union of the sets. The resulting value ranges from 0 to 1. Similarly, cosine similarity, which quantifies the cosine of the angle between two vectors, was used to measure the similarity between two spike vectors. Cosine similarity is given by

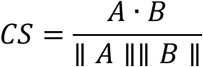

where A and B are the two vectors being compared, *A* · *B* is the dot product, and ∥ *A* ∥ and ∥ *B* ∥ represent the magnitudes of the vectors. The values range from -1 to 1 where 1 indicates identical vectors and -1 indicates that the vectors have an angular difference of 180 degrees. Actual spike counts were used for cosine similarity calculation.

### Rank-order correlation

To evaluate the consistency with which populations of neurons in PFC fired in sequence during chained events, we calculated the rank-order correlation between each chain and its template as previously described (**Figures 5C** and **S7B**).^112,113^ The population activity during each HFO in a chained event was concatenated (inter-event activity excluded), and for each concatenated event with ≥5 cells active, the spiking activity was reduced to each unit’s first spike time, chronologically ordered, and normalized from 0 to 1. Then, a cross-validated template-based method was used to calculate sequence similarity between each event and its template. For each event, a template was created by averaging the normalized rank of each cell across all events excluding the event in question. The rank correlation was reported as the mean Pearson correlation coefficient between the event and template across all events. For comparison, a null distribution was generated by jittering spike times (1000 jitters) and performing the rank-order analysis above. For the pseudo-chain control, each HFO chain was time-shifted ±5 seconds such that each HFO within the chain was shifted the same value (1000 time shifts), thus preserving the inter-event-intervals (i.e. The entire chain was shifted coherently). Additionally, the entire shifted chain must have stayed within a REM sleep bout. The above rank-order procedure was performed, and the real correlation was compared to the pseudo-chain null distribution.

### Assembly detection

To detect cell assemblies in PFC populations, a previously described method was used (**Figure 6**).^50,51^ Significant coactivity patterns in PFC were detected using a method based on principal and independent component analyses. For each W-Track epoch, the spike trains of each neuron were binned into 20 ms time windows and Z-scored to eliminate any biases due to differences in firing rates. Principal component analysis was then applied to the Z-scored spike matrix (Z). The resulting eigenvalue decomposition of the correlation matrix (C) was given by:

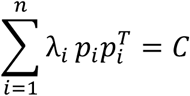

Where *p*_*i*_ is the *i*^*th*^ principal component of C and *λ*_*i*_ is the corresponding eigenvalue. Note that 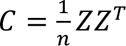 is the correlation matrix of Z. To determine the number of significant coactivity patterns in the data, the Marcenko-Pastur law was used to calculate a threshold eigenvalue λ_*max*_, which was given by 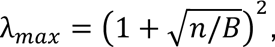 where *n* is the number of recorded units and *B* is the total number of bins. Note that an eigenvalue exceeding *λ*_*max*_indicates that the corresponding cell assembly explains more correlation than expected if the activity of neurons was independent of each other. The number of eigenvalues above λ_*max*_, *N*_*A*_, represents the number of significant patterns found in the data. These principal components were then projected back onto the original z-scored spike data:

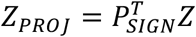

where *P*_*SIGN*_ is the *nXN*_*A*_ matrix with *N*_*A*_ columns. Independent component analysis (ICA, RobustICA)^114^ was then used to identify patterns that were maximally independent from each other. ICA was applied to the matrix *Z*_*PROJ*_ to find an *N*_*A*_*XN*_*A*_ unmixing matrix, *W*, which was used to obtain the weights of each cell in each assembly.

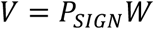

Since the sign of the output of ICA is arbitrary, the signs of the weight vector were set such that the highest absolute weight was set to positive.

### Assembly reactivation strength

To assess the reactivation of these detected assemblies during sleep sessions (**Figure 6**), the expression strength of each pattern over time was given by:

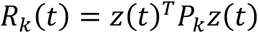

*R*_*k*_(*t*) is the reactivation strength of assembly *k* at time *t*, *z*(*t*) is the z-scored firing rate vector for all neurons at time *t*, and *P*_*k*_ is the projection matrix for assembly *k*, constructed by taking the outer product of its weight vector 𝑣_*k*_. Additionally, the diagonal of the projection matrix was set to zero to reduce the influence of highly active neurons, and the reactivation strength was calculated such that only the positive weight cells were assessed. Thus, only patterns of coactivity are detected as periods of high reactivation strength.

### Assembly sequence detection surrounding HFOs

The consistency of assembly reactivation surrounding REM HFOs was assessed by implementing a cross-validation process similar to procedures used for place cell sequences (**Figures 6A-D**).^115,116^ We employed a split-event procedure where HFO events were randomly split into two halves. The average reactivation strength for each assembly was then calculated across all HFOs across both halves separately. This was performed across all animals and epochs. The assemblies aligned to the first half were subsequently sorted by peak reactivation strength within a ±500 ms window around HFO times, and the assemblies aligned to the second half were sorted using the same sort indices such that each row in the two matrices correspond to the same assembly. The consistency of temporal assembly reactivation was quantified as the Pearson correlation between the peak reactivation strengths of the first and second halves. This procedure was conducted 1000 times using different subsets of HFOs for the first and second halves to obtain a distribution or correlation coefficients. For comparison, a distribution of shuffled data was obtained by circularly shuffling assembly reactivation strength prior to HFO alignment, and the above procedure was performed. This procedure was also performed for gamma and NREM PFC ripples for comparison.

Furthermore, the slope and peak differences were calculated as additional metrics of sequence and temporal reactivation consistency between the two halves of data, respectively. For the slope difference metric, the slope of the best fit line between assembly ID and peak reactivation index was taken for the two halves of data and compared. A smaller slope difference compared to shuffled data indicates that assembly reactivation is structured in a more similar manner across the two halves of the real data. For peak reactivation difference, the temporal displacement of the peak reactivation index between the two halves of data was calculated and compared to shuffle. A high probability of small peak differences indicates that the timing of peak reactivation of assemblies relative to HFO chain onset is similar across the two halves of data. Shuffling of assembly strength was carried out as above.

### Reactivation strength across epochs

For comparison of pre and post reactivation strengths, the same run-derived templates were projected onto the preceding and following sleep epochs (neurons were matched across pre, run, and post), and the pre and post strengths during periods of sleep were compared. Thus, each assembly was matched across pre and post. Furthermore, to assess reactivation strength changes over learning, the preceding run epoch’s template (following run epoch for sleep epoch #1, since no preceding run) was projected onto the following sleep epoch, and the average reactivation strength was calculated and compared for early (epochs 1-3) and late (epochs 4-9) sleep epochs during periods of sleep. Epochs prior to a grouped average of 75% correct performance on the W-Track task (**Figure S1B**) were defined as early.

### 2D#assembly maps

To generate assembly rate maps, a procedure similar to computing place fields was used.^51^ For each assembly, all assembly activation timestamps during W-Track behavioral sessions that exceeded 5 at positional bins where occupancy was >20 ms were used for map generation. 2-D assembly maps were calculated by dividing the binned activation count by the occupancy in each 1 cm^2^ positional bin and smoothed with a Gaussian kernel (σ = 2).

### Spatial information of assembly fields

Spatial information (bits/event activation) was computed using the following formula:

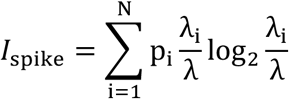

where p_i_ is the occupancy probability of bin *i*, λ_i_ is the mean assembly activation in bin *i*, and λ is the occupancy-weighted mean activation. Spatial information is reported in bits per activation event. The significance of each assembly’s spatial information was assessed by a per-assembly circular-shift procedure. The full assembly activation time series was circularly shifted by a random offset >20 s, the activation shuffled 2D assembly field was computed, and the spatial information was calculated. This procedure was repeated 1000 times per assembly to generate a null distribution. The observed spatial information was expressed as a z-score relative to the assembly-specific shuffle distribution, and a one-tailed permutation p-value was computed as p = (1 + #[*I*_shuffled_ ≥ *I*_Observed_]) / (1 + 1000). Assemblies with p < 0.05 (z > 1.65) were classified as showing significant spatial structure (**Figures 6H,I**).

### 2-D rate maps

2-D spatial rate maps during active behavior on the W-Track were calculated during periods of high mobility (>5 cm/s; all SWR times excluded) at positional bins where occupancy was >20 ms. Then, the 2-D spatial maps were calculated by dividing the binned spike count by the occupancy in each 1 cm^2^ positional bin and smoothed with a Gaussian kernel (σ = 2).

### Calculation of a single cofiring metric and separation into populations of high and low cofiring neurons

Since we wanted to relate the above cofiring metric to other measures, we needed to obtain a single value cofiring metric for each neuron. To do this, we averaged the cofiring values across all pairings for a neuron (e.g. 1 CA1 neuron paired with all PFC neurons) and reported it as the cell’s cofiring. Furthermore, since we observed a bimodal distribution of CA1-PFC cofiring values in REM sleep, we split the population based on whether the average (across all cell-cell combinations) cofiring value or correlation coefficient of a cell was above or below 0 (High cofiring > 0; low cofiring < 0). Also, since the NREM ripple cofiring distribution was unimodal, we additionally split the populations using the mean of the average cofiring or correlation coefficient distributions (High cofiring > mean; low cofiring < mean).

### Change in firing rate across sleep

To assess differences in firing rate changes in CA1 and PFC over the course of sleep, populations were separated into low or high cross-regional cofiring units based on the z-scored cofiring metric above as well as the Pearson correlation between spike count vectors across all REM HFO chains or independent NREM ripples. Here, we are only including independent ripples for NREM analysis since we have shown that CA1-PFC cofiring is elevated when PFC ripples are coordinated with CA1 SWRs.^28^ Exclusion of coordinated ripple events allows us to assess the influence of strictly PFC ripples in NREM on firing rate changes. For each cell, the change in the firing rate from the first bout to the last bout of NREM sleep was calculated and compared for low and high coactive cells (**Figures 7F** and **S9B-E**). Only epochs that had first and last NREM bouts exceeding 30 s in length were included for analysis.

### REM theta phase shifting

To assess shifts in the preferred CA1 theta phase of CA1 neurons between run sessions and REM sleep, the above procedure was performed, and only neurons that exhibited significant phase-locking during both run and REM sleep were included for analysis. Neurons that had a phase shift of >90° from the preceding run session to the subsequent REM sleep epoch were categorized as REM-shifting.

### Single unit PFC HFO bursting

To determine HFO burst probability for each cell, the number of HFOs where burst spikes were detected (at least 2 spikes <6 ms apart^117^) was divided by the total number of events.

### Network model

Computational simulations were done using a simplified model of a small cortical network. The model contains 1,600 excitatory pyramidal and 400 inhibitory interneuron modified Hodgkin-Huxley neurons laid out in a topographical grid. Connections are random, with probability of connection decaying with spatial distance on the grid. Connection probabilities were drawn from known layer 2/3 cortical connectivity rates of pyramidal cells and somatostatin (SST) interneurons. Synaptic weights were assigned randomly using log-normal distributions. Inhibitory cells in the model are regular-spiking with a voltage equation identical to pyramidal cells.

The voltage equation for the neuron model is:

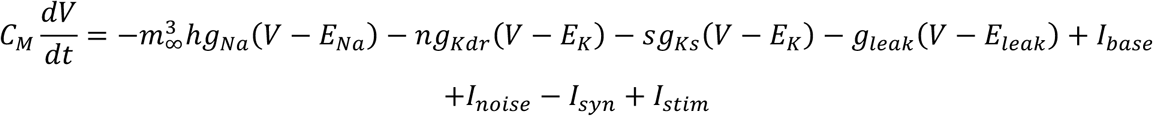

Where *g*_*x*_ are the maximal conductances, and each of the activation variables *h*, *n*, and *s* have a differential equation of the form:

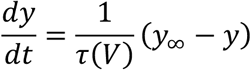

Where *y*_∞_ are the steady state values for each of the gating variables, each a function of voltage.^62^

For every presynaptic spike, the synaptic conductance between neurons is calculated as the difference between two decaying exponentials. Model pyramidal neurons and interneurons both elicit fast, large-amplitude synaptic inputs in connected postsynaptic cells modeled after glutamate and GABA-A receptors, respectively. Interneurons also elicit a slow, low-amplitude GABA-B-like conductance in connected neurons. In addition, a white noise current *I*_noise_ is applied to each neuron separately to introduce random variation.

Both pyramidal neurons and interneurons in the model feature a slow potassium conductance, *g*_*Ks*_, that is active at neuronal resting potential and modeled after the slow potassium M-current.^64,118^ The biological effect of acetylcholine (ACh) on the M-current through the M1 muscarinic ACh receptor is recreated in the model by raising or lowering *g*_*Ks*_ to simulate decreases or increases, respectively, in ACh concentration. As in biological neurons, lowering *g*_*Ks*_ shifts model neurons from Type 2 excitability (resonators) to Type 1 excitability (integrators), raising their excitability but lowering their tendency to synchronize spiking when connected.^66^ The result for a connected network of neurons is, in general, lower sensitivity to inputs and an increased propensity for synchronous spiking in simulations modeling a low ACh condition, but increased sensitivity and less correlated spiking in simulations modeling a high ACh condition, as is observed experimentally.^119,120^ Model simulations of NREM sleep use *g*_*Ks*_ = 1.5 for all neurons, while simulations of REM sleep use *g*_*Ks*_ = 0. *I*_*base*_ is used to slightly hyperpolarize interneurons during REM to maintain balanced excitation and inhibition.

Each external stimulus input to the network *I*_*stim*_ is modeled as a depolarizing current injection to a random 30% of neurons. For the NREM brief stimuli, a gaussian profile that peaks at the midpoint of the stimulus was used, with duration drawn from a normal distribution with a mean of 70 ms and standard deviation 20 ms. Stimulus peak amplitude was calculated similarly, with a mean of 0.75 and standard deviation 0.3. For REM theta stimuli, the input was a sinusoid with frequency drawn from normal distribution with mean 8.5 Hz and 3 standard deviations within 7-10 Hz range. Stimulus amplitude was convolved with a gaussian profile peaking at the midpoint of the stimulus duration. Negative amplitude phases of the sinusoid were set to zero. For REM stimulus duration, mean 3000 ms and standard deviation 250 ms duration were used. Peak amplitude parameters were identical to NREM. No negative durations or amplitudes were allowed and set to zero if drawn from the distribution. Triggering of all stimulus events was determined by a Bernoulli process with a success rate of on average 1 Hz, with overlapping stimuli not permitted. For theta stimuli, a minimum delay of 1000 ms was enforced after the end of each stimulus to allow the network time to return to baseline activity.

## QUANTIFICATION AND STATISTICAL ANALYSIS

All statistical analyses were performed with MATLAB (MathWorks, Natick, MA; R2022a) functions or custom scripts. No specific analysis was performed to predetermine sample size, but the number used in this study is similar to or larger than typically used. Nonparametric, two-tailed statistical tests (Wilcoxon rank-sum (WRS) and signed-rank (WSR), for unpaired and paired data, respectively) were used throughout the paper unless noted. P < 0.05 was considered the cutoff for statistical significance. Boxplots show the median (center line), interquartile range (box), and the furthest points not considered outliers (whiskers). Unless otherwise noted, values and errors in the text denote means ± SEM. p-values in the text are reported as follows: p > 0.05, *p < 0.05, **p < 0.01, ***p < 0.001.

---

**Table S1.** Amount of NREM and REM sleep (in seconds) across all animals and epochs.

| Animal | Epoch | NREM | REM | % REM | Animal | Epoch | NREM | REM | % REM |
| --- | --- | --- | --- | --- | --- | --- | --- | --- | --- |
|  | 1 | 590 | 0 | 0.00% |  | 1 | 715 | 121 | 14.47% |
|  | 2 | 574 | 0 | 0.00% |  | 2 | 548 | 0 | 0.00% |
|  | 3 | 389 | 11 | 2.75% |  | 3 | 1064 | 13 | 1.21% |
|  | 4 | 362 | 0 | 0.00% |  | 4 | 1063 | 137 | 11.42% |
| **JS17** | 5 | 629 | 0 | 0.00% | **BG1** | 5 | 623 | 72 | 10.36% |
|  | 6 | 404 | 0 | 0.00% |  | 6 | 1778 | 275 | 13.40% |
|  | 7 | 1197 | 10 | 0.83% |  | 7 | 1636 | 263 | 13.85% |
|  | 8 | 1139 | 0 | 0.00% |  | 8 | 1284 | 270 | 17.37% |
|  | 9 | 916 | 74 | 7.47% |  | 9 | 693 | 27 | 3.75% |
|  | Total | **6200** | **95** | **1.51%** |  | Total | **9404** | **1178** | **11.13%** |
|  | 1 | 414 | 11 | 2.59% |  | 1 | 595 | 31 | 4.95% |
|  | 2 | 499 | 15 | 2.92% |  | 2 | 49 | 0 | 0.00% |
|  | 3 | 836 | 36 | 4.13% |  | 3 | 186 | 38 | 16.96% |
|  | 4 | 1124 | 0 | 0.00% |  | 4 | 678 | 0 | 0.00% |
| **JS13** | 5 | 1040 | 139 | 11.79% | **JS14** | 5 | 605 | 0 | 0.00% |
|  | 6 | 1926 | 120 | 5.87% |  | 6 | 901 | 167 | 15.64% |
|  | 7 | 1322 | 118 | 8.19% |  | 7 | 661 | 86 | 11.51% |
|  | 8 | 1440 | 122 | 7.81% |  | 8 | 594 | 126 | 17.50% |
|  | 9 | 1180 | 141 | 10.67% |  | 9 | 814 | 95 | 10.45% |
|  | Total | **9781** | **702** | **6.70%** |  | Total | **5083** | **543** | **9.65%** |
|  | 1 | 8 | 0 | 0.00% |  | 1 | 138 | 177 | 56.19% |
|  | 2 | 840 | 98 | 10.45% |  | 2 | 750 | 158 | 17.40% |
|  | 3 | 497 | 275 | 35.62% |  | 3 | 1066 | 124 | 10.42% |
|  | 4 | 1266 | 202 | 13.76% |  | 4 | 863 | 128 | 12.92% |
| **KL8** | 5 | 1571 | 113 | 6.71% | **JS21** | 5 | 435 | 14 | 3.12% |
|  | 6 | 629 | 182 | 22.44% |  | 6 | 649 | 0 | 0.00% |
|  | 7 | 1346 | 256 | 15.98% |  | 7 | 818 | 23 | 2.73% |
|  | 8 | 1193 | 243 | 16.92% |  | 8 | 859 | 13 | 1.49% |
|  | 9 | 568 | 41 | 6.73% |  | 9 | 743 | 0 | 0.00% |
|  | Total | **7918** | **1410** | **15.12%** |  | Total | **6321** | **637** | **9.15%** |
|  | 1 | 3 | 0 | 0.00% |  | 1 | 54 | 0 | 0.00% |
|  | 2 | 76 | 12 | 13.64% |  | 2 | 5 | 0 | 0.00% |
|  | 3 | 291 | 45 | 13.39% |  | 3 | 133 | 0 | 0.00% |
|  | 4 | 145 | 0 | 0.00% |  | 4 | 637 | 26 | 3.92% |
| **JS34** | 5 | 238 | 15 | 5.93% | **JS12** | 5 | 562 | 11 | 1.92% |
|  | 6 | 1513 | 274 | 15.33% |  | 6 | 1319 | 114 | 7.96% |
|  | 7 | 453 | 69 | 13.22% |  | 7 | 1178 | 83 | 6.58% |
|  | 8 | 535 | 75 | 12.30% |  | 8 | 1333 | 190 | 12.48% |
|  | 9 | 1070 | 112 | 9.48% |  | 9 | 720 | 0 | 0.00% |
|  | Total | **4324** | **602** | **12.22%** |  | Total | **5941** | **424** | **6.66%** |
|  | 1 | 731 | 30 | 3.94% |  | 1 | 25 | 0 | 0.00% |
|  | 2 | 435 | 0 | 0.00% |  | 2 | 504 | 16 | 3.08% |
|  | 3 | 742 | 0 | 0.00% |  | 3 | 885 | 34 | 3.70% |
|  | 4 | 865 | 0 | 0.00% |  | 4 | 919 | 14 | 1.50% |
| **JS15** | 5 | 1115 | 191 | 14.62% | **ZT2** | 5 | 1658 | 240 | 12.64% |
|  | 6 | 1476 | 0 | 0.00% |  | 6 | 1484 | 151 | 9.24% |
|  | 7 | 938 | 116 | 11.01% |  | 7 | 1573 | 47 | 2.90% |
|  | 8 | 1195 | 136 | 10.22% |  | 8 | 1804 | 239 | 11.70% |
|  | 9 | 990 | 224 | 18.45% |  | 9 | 716 | 12 | 1.65% |
|  | Total | **8487** | **697** | **7.59%** |  | Total | **9568** | **753** | **7.30%** |

**Figure S1.**
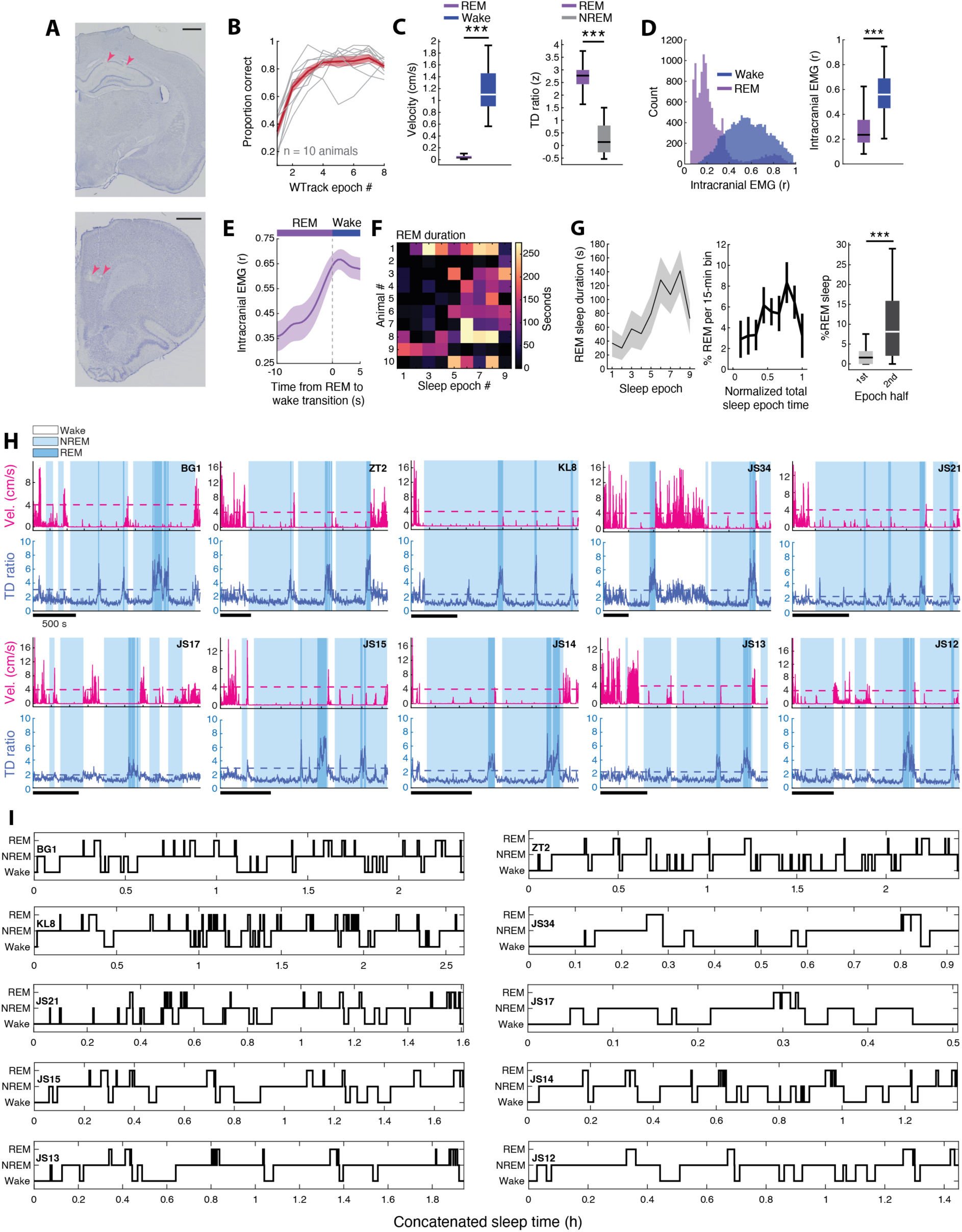
related to Figure 1. Characterization and validation of REM sleep. (**A**) Histological verification of recording locations in dorsal CA1 (Top) and PFC (Bottom). Red arrows indicate electrolytic lesions, and scale bars represent 1 mm. (**B**) Performance of all 10 animals across 8 sessions on the W-Track task. Outbound and inbound trajectories were combined to calculated overall performance on the task. (**C**) (Left) Average velocity during wake and REM sleep (Wake = 1.16 ± 0.06 cm/s, REM = 0.05 ± 0.01 cm/s, *p = 5.39×10^-7^, WSR). (Right) Theta-to-delta (TD) ratio in NREM and REM sleep (NREM = 0.30 ± 0.11, REM = 2.71 ± 0.10, ***p = 5.39×10^-7^, WSR). (**D**) Distributions of intracranial EMG values for wake and REM sleep. All 500 ms binned Pearson correlation values are shown. (**E**) Comparison of the averaged intracranial EMG for bouts of wake and REM during all sleep epochs (Wake = 0.57 ± 0.01, REM = 0.32 ± 0.02, ***p = 5.26×10^-37^, WRS) (**F**) Duration of REM sleep in each epoch (Total duration = 6089 s). Only epochs with at least 30 s of REM sleep were included for analysis. (**G**) (Left) Average REM sleep duration and (Middle) percentage of REM sleep out of total sleep epoch time (Wake, NREM, and REM) across all 9 sleep epochs. No animals had REM sleep in the first normalized bin, so that point isn’t shown. (Right) Percentage of REM sleep during the first and second halves of the 36 sleep epochs included in analysis (First = 3.85 ± 0.93%, Second = 15.29 ± 0.94%, ***p = 1.93×10^-6^, WRS). (**H**) Example sleep sessions for each animal showing each behavioral state detected using the sleep state algorithm that utilizes theta-to-delta ratio (TD ratio) and animal velocity (cm/s). Note that the plot for KL8 is the example shown in **Figure 1B**. (**I**) Hypnograms for animals showing sleep state across all concatenated sleep epochs that passed inclusion criteria (only sleep epochs are shown; behavior epochs are excluded).

**Figure S2.**
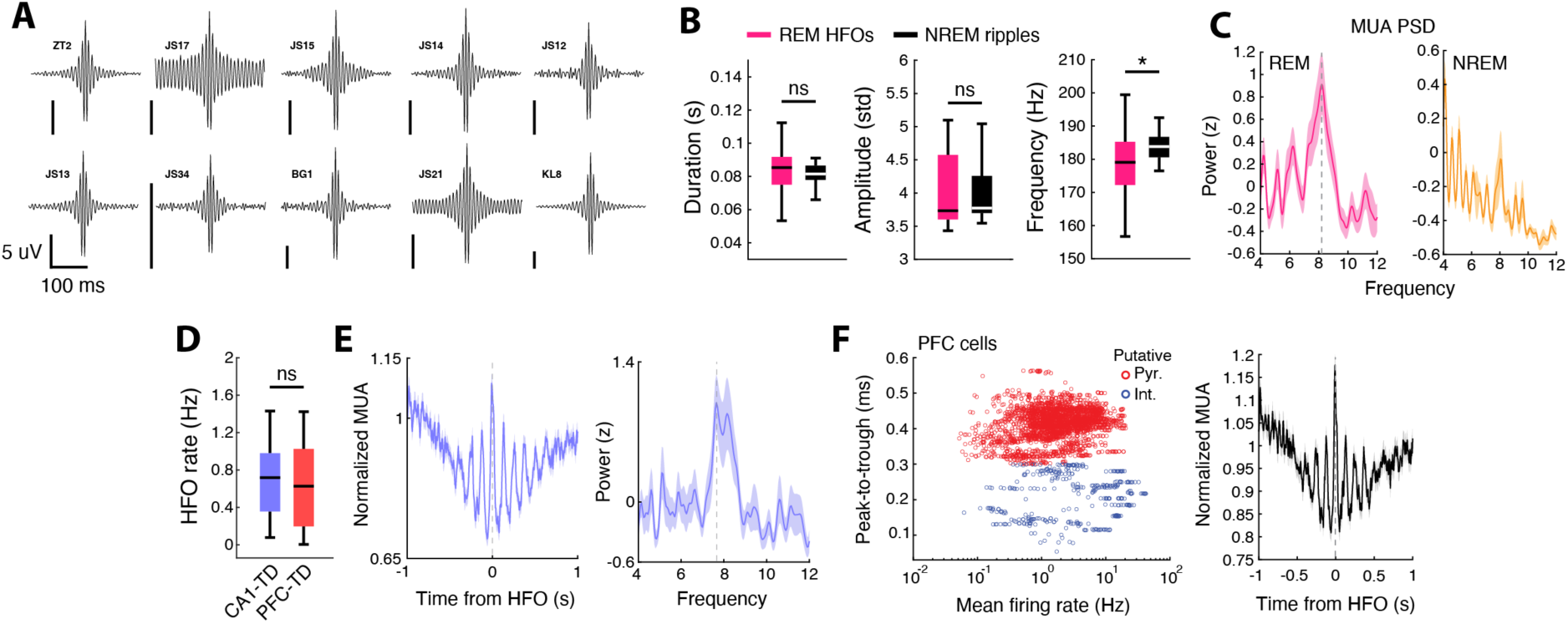
Related to Figures 1 and 2. Characterization prefrontal REM sleep HFOs and event aligned multiunit activity. (**A**) Average peak aligned amplitude plots across all detected REM HFO events for each animal. (**B**) Comparison of ripple/HFO duration, amplitude (standard deviations above the mean), and frequency for NREM and REM events (Duration: NREM = 0.111 ± 0.017 ms, REM = 0.126 ± 0.030 ms, p = 0.59; Amplitude: NREM = 4.13 ± 0.12, REM = 4.23 ± 0.20, p = 0.31; Frequency: NREM = 184.96 ± 1.68 Hz, REM = 179.02 ± 1.82 Hz, *p = 0.018, WRS). (**C**) Average power spectral density calculated from the REM and NREM event aligned multiunit activity in PFC. Note the peak around 8 Hz (theta frequency) for REM HFO aligned multiunit activity. (**D**) PFC HFO rates during REM sleep extracted from either CA1 or PFC TD ratio. HFO rates were similar during CA1 and PFC extracted REM sleep, validating the use of CA1-extracted REM sleep for further analysis. (**E**) PFC multiunit activity and average power spectral density aligned to REM HFO events extracted from the overlap of CA1 and PFC REM sleep. (**F**) (Left) Peak-to-trough of average spike waveforms and mean firing rate of PFC neurons. We separated populations based on peak-to-trough, since mean firing rate did not give us clear separation. Putative interneurons were identified as cells having a peak-to-trough <0.3 ms. (Right) REM HFO aligned PFC multiunit response when spikes from putative interneurons are excluded (compare to **Figure 2A**). Due to this similarity of the phasic PFC response when putative interneurons are omitted, we decided to pool PFC neurons for all further analyses.

**Figure S3.**
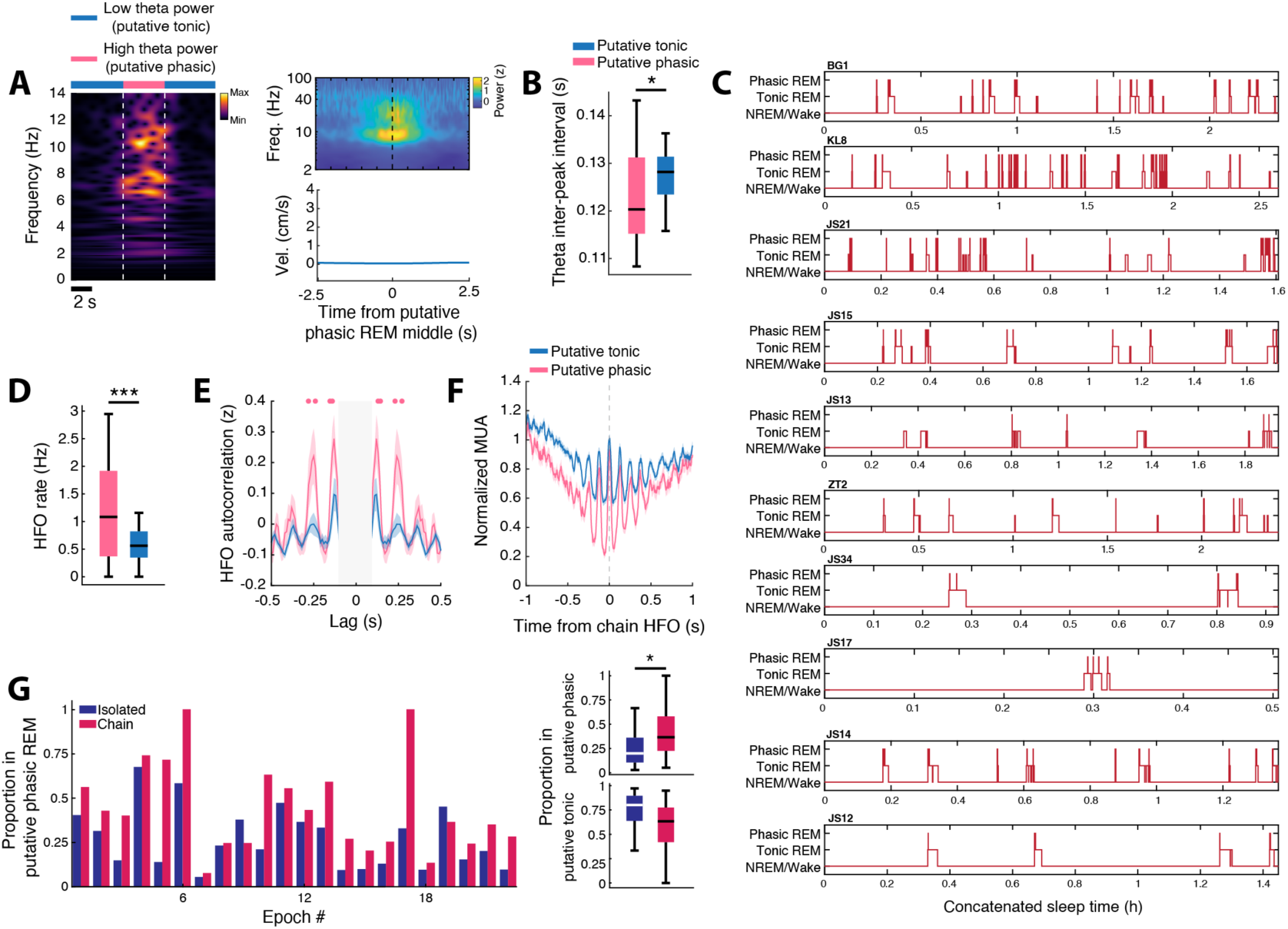
Related to Figure 3. Putative tonic and phasic REM sleep. (**A**) (Left) Example showing bouts of low and high theta power within a REM sleep bout. High theta periods were bouts within REM that exceeded 4 standard deviations above the electrode averaged theta power in REM sleep for at least 900 ms. (Right) Average spectrogram and speed surrounding putative phasic REM bouts. (**B**) The CA1 theta inter-peak intervals (IPI) during high PFC theta power periods during REM sleep were lower (corresponding to higher theta frequency) than during low power periods, indicating that these high and low theta power periods may correspond to putative phasic and tonic substages of REM sleep, respectively^15^ (High theta IPI = 0.123 ± 0.002, Low theta IPI = 0.128 ± 8.3×10^-4^, *p = 0.010, WRS). (**C**) REM sleep focused hypnograms for all animals illustrating sleep state concatenated across all epochs that were included for analysis (as in **Figure S1I**). Wake and NREM sleep are combined. (**D**) Rate of REM HFOs during bouts of high or low theta power (High = 1.19 ± 0.15 Hz, Low = 0.58 ± 0.05 Hz, ***p = 1.33×10^-4^, WSR). (**E**) HFO autocorrelation during periods of high or low theta power. Dots indicate bins where high and low theta power HFO autocorrelations differ (p<0.05, WRS). (**F**) PFC multiunit activity aligned to chain HFOs in either putative tonic or phasic REM sleep. Similar patterns of PFC spiking modulation are elicited aligned to HFOs in both REM substates. (**G**) (Left) Proportion of isolated and chain HFOs that occurred in putative phasic REM sleep in all 22 sleep epochs that had at least 5 seconds of putative phasic REM sleep and (Right, top) a quantification of the difference between isolated and chain events (Isolated = 0.25 ± 0.04, Chain = 0.43 ± 0.06, *p = 0.018, WSR). (Right, bottom) Proportion of isolated and chain events in putative tonic REM shown as well.

**Figure S4.**
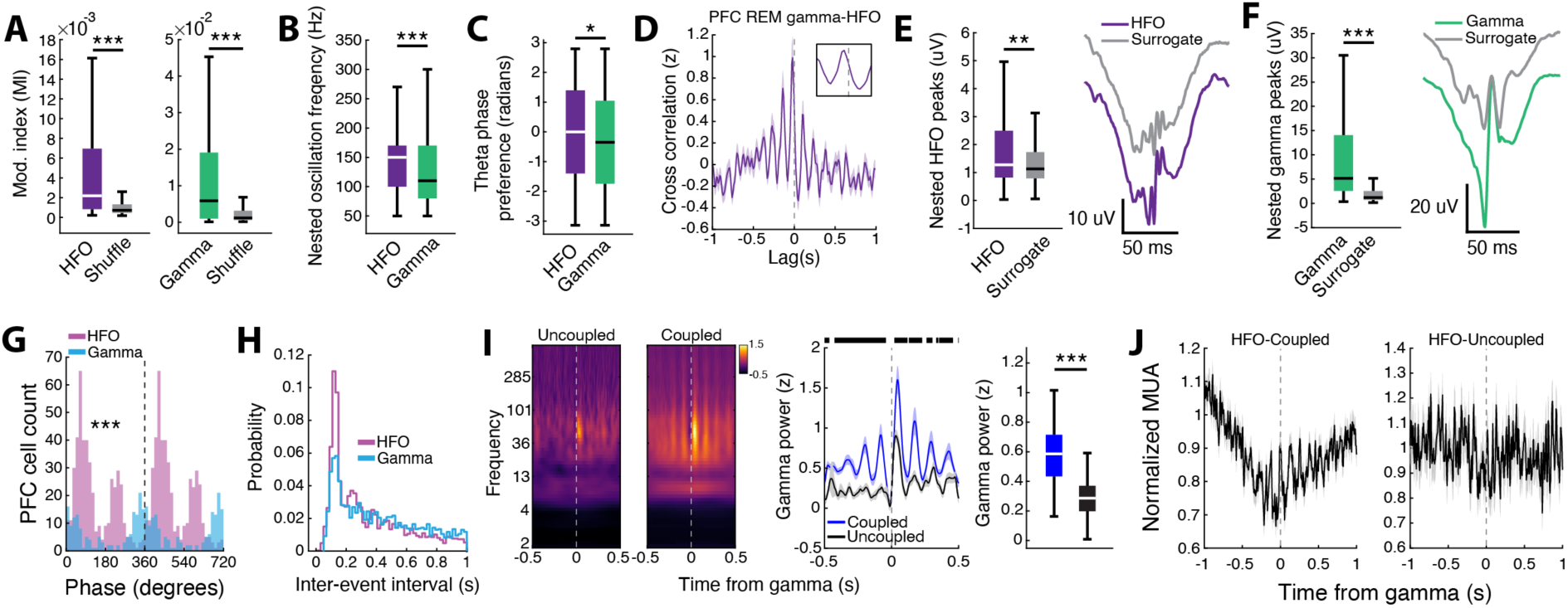
Related to Figure 3. Coupling of gamma and HFOs with PFC theta oscillations. (**A**) Modulation indices (MIs) for phase amplitude coupling (PAC) computed for HFO and gamma frequency oscillations compared to MIs calculated from circularly shifted theta phase and HFO/gamma amplitude vectors. Modulation was assessed for a phase frequency range from 2-12 Hz. The amplitude modulated frequencies were 150-250 Hz and 40-100 Hz for HFO and gamma, respectively (HFO = 0.005 ± 3.21×10^-4^, shuffle = 0.001 ± 8.94×10^-5^, ***p = 1.13×10^-31^; Gamma = 0.011 ± 6.49×10^-4^, shuffle = 0.002 ± 1.67×10^-4^, ***p = 2.97×10^-24^, WRS). (**B**) Frequency of the average theta nested oscillation either aligned to peak HFO or gamma power (See **Methods**). Note that aligning to peak HFO power at the preferred phase bin yields nested oscillations that are higher in frequency (HFO = 146.48 ± 2.82, Gamma = 128.96 ± 3.091, ***p = 2.85×10^-7^, WRS). (**C**) Preferred theta phases of elevated HFO and gamma amplitude (HFO = -0.13 ± 0.08 radians, Gamma = -0.38 ± 0.08 radians, *p = 0.016, WRS). (**D**) Cross-correlation between PFC gamma and HFO events during REM sleep. The off-center peak at a negative lag (inset) indicates that gamma events tend to precede HFOs, corroborating the phase preference difference in (**F**). (**E**) Comparison of nested HFO peaks with a theta phase bin shuffled surrogate (See **Methods**; HFO = 4.09 ± 0.48 μV, Surrogate = 1.92 ± 0.19 μV, **p = 0.0058, WRS). (**F**) Same as in (**E**) but for gamma (Gamma = 11.33 ± 0.62 μV, Surrogate = 2.33 ± 0.20 μV, ***p = 9.46×10^-94^, WRS). (**G**) Distribution of preferred phases for PFC cells that are significantly phase locked to either HFOs or gamma. Note the similarity in distribution to **Figure 3D** in the manuscript (HFO locked = 463 neurons, Gamma locked = 161 neurons, Dual locked = 110 neurons, U2 = 3.56, ***p = 6.37×10-31, Watson’s U2 test) (**H**) Distributions of inter-event intervals for PFC gamma events and HFOs (***p = 5.09×10^-30^, Kolmogorov-Smirnov test). (**I**) (Left) Gamma triggered spectrograms for events that were uncoupled or coupled to HFOs. (Middle) Comparison of gamma power (40-100 Hz power ±500 ms from event onset) for uncoupled and coupled events. Note the oscillating gamma power during coupled events. (Right) Quantification of average HFO power for coupled and uncoupled events (Coupled = 0.60 ± 0.04, Uncoupled = 0.34 ± 0.03, ***p = 1.03×10^-7^, WRS). Coupled gamma events were defined as gamma periods that were within 500 ms of a HFO event. (**J**) (Left) Multiunit PFC activity aligned to gamma events that are coupled with or (Right) uncoupled from HFOs.

**Figure S5.**
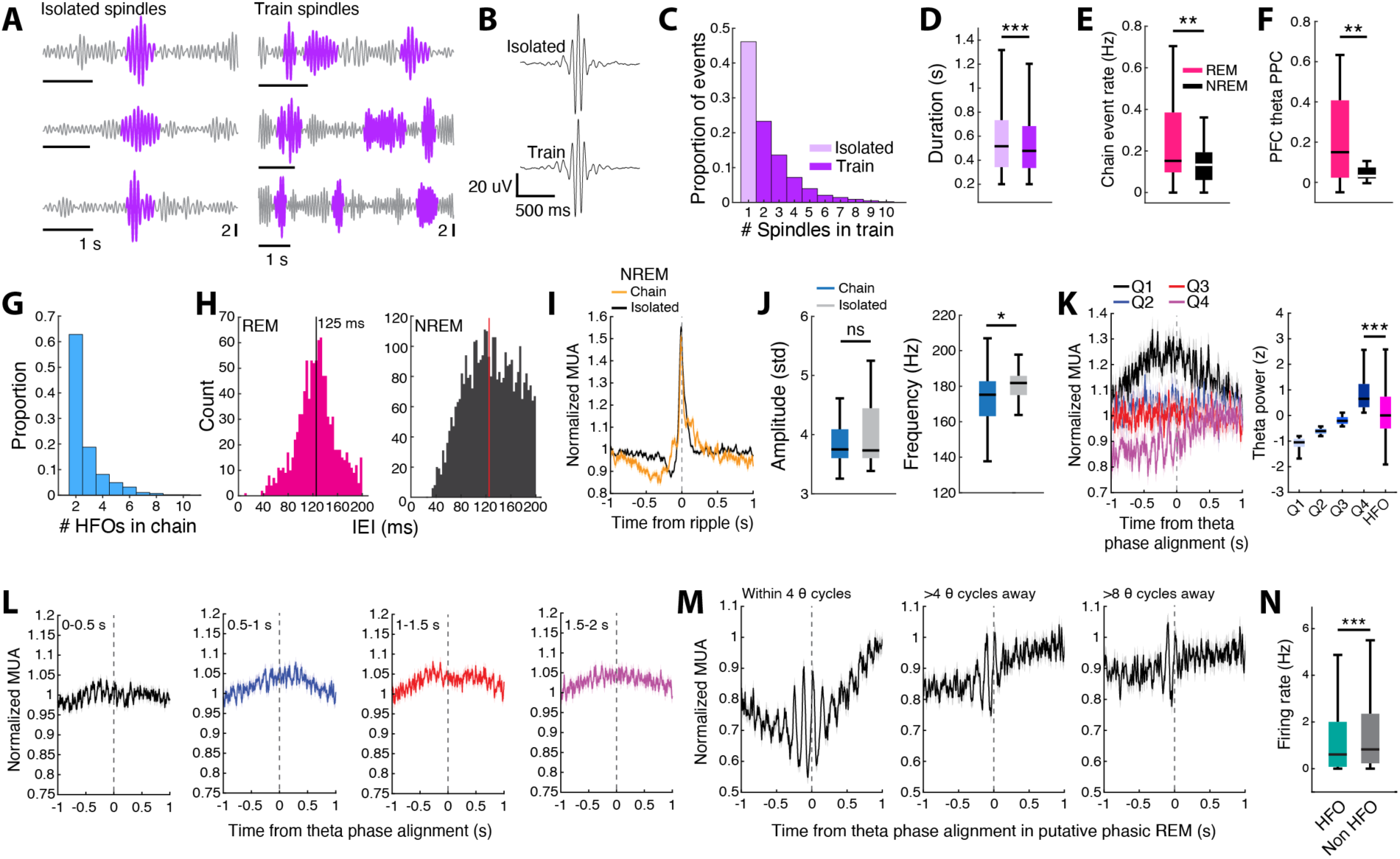
Related to Figures 3 and 4. Chains of sleep oscillations in NREM (spindles) and REM (HFOs) sleep. (**A**) Example isolated and train spindles, similar to previous reports.^45^ (**B**) Averaged spindle peak aligned LFP for isolated and train spindles across all animals. (**C**) Histogram displaying the proportions of NREM spindle train events of specific lengths (Compare with Darevsky et al., 2024). (**D**) The durations of isolated and train spindles (Isolated = 0.579 ± 0.005 s, Train = 0.550 ± 0.003, ***p = 1.77×10^-6^, WRS). (**E**) Rate of events designated as participating in a chain (<200 ms separation) during NREM and REM sleep (NREM = 0.15 ± 0.02 Hz, REM = 0.25 ± 0.04 Hz, **p = 0.0054, WSR). (**F**) PFC theta pairwise phase consistency (PPC) of HFO chains in REM vs NREM (REM = 0.22 ± 0.06, NREM = 0.06 ± 0.02, **p = 0.0097, WSR). (**G**) Histogram displaying the proportions of REM HFO chain events of specific lengths. (**H**) Histogram of all inter-event intervals (IEI) for HFO chains in (Left) REM or (Right) NREM sleep. Note the tight REM HFO IEI distribution centered around 125 ms, which corresponds to 8 Hz (theta frequency). Black and red vertical lines are displayed at 125 ms for both distributions. (**I**) PFC multiunit activity aligned to chained and isolated NREM ripple events. Note the absence of theta-modulated population activity during ripple chains in NREM sleep. (**J**) Comparison of HFO amplitude and frequency for chained and isolated PFC REM HFOs (Amplitude: Chain = 4.21 ± 0.26, Isolated = 4.17 ± 0.20, p = 0.995; Frequency: Chain = 172.54 ± 2.71 Hz, Isolated = 181.54 ± 1.51 Hz, *p = 0.018, WRS). (**K**) (Left) PFC multiunit activity aligned to the preferred theta phase of REM HFOs plotted according to theta power during surrogate events. (Right) Quantification of theta power in each quartile compared to the power during HFO events (Q1 = -1.10 ± 0.007, Q2 = -0.62 ± 0.004, Q3 = -0.20 ± 0.005, Q4 = 0.87 ± 0.024, HFO = 0.26 ± 0.018, ***p = 5.74×10^-114^, WRS). (**L**) PFC multiunit activity aligned to preferred theta phase bins within the specified time window around HFOs. For example, for the figure on the left, theta phase bins must have been within 0-0.5 s from the beginning or end of HFO windows for inclusion. Note that this analysis sampled phase bins across both putative tonic and phasic REM sleep. (**M**) PFC multiunit activity aligned to preferred theta phase bins within the specified time window around HFOs. (**N**) PFC neuron firing rate comparison during chain HFOs and theta periods >4 theta cycles away from HFOs during putative bouts of phasic REM sleep. A ±250 ms window around events was used to calculate the average firing rate across events in each condition (HFO = 1.56 ± 0.10 Hz, non-HFO = 1.75 ± 0.11 Hz, ***p = 6.50×10^-5^, WSR).

**Figure S6.**
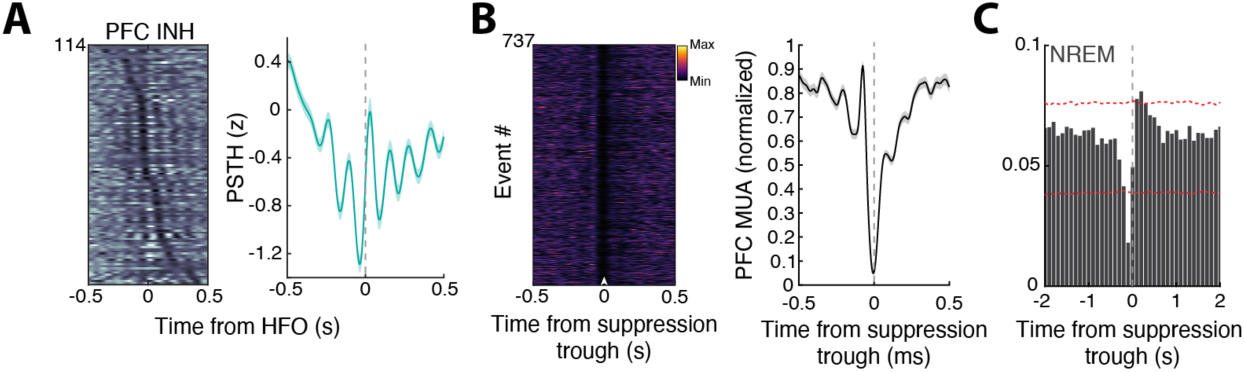
Related to Figure 4. REM PFC HFOs are associated with decreases in PFC activity. (**A**) PFC neurons that were negatively modulated (INH) during REM PFC HFO events (REM EXC modulation = 114). Although overall modulation in this subset of neurons surrounding PFC HFOs is suppressive, there are transient peaks in activity that occur approximately 125 ms apart (8 Hz theta frequency). (**B**) Extracted PFC population suppression events (see **Methods**) aligned by their troughs. (Left) All example suppression events in an example epoch. (Right) Averaged PFC multiunit activity aligned to these events. (**C**) The probability of NREM PFC ripples centered on population suppression events detected in NREM sleep. Unlike REM sleep, there is a decreased probability of NREM ripple occurrence surrounding suppression events, consistent with strong overall excitation during these events (**Figure 2C**). The slight peak to the right is likely reflective of the suppression in activity prior to the onset of NREM ripples (**Figure 2C**). The red dotted lines indicate the 99% confidence intervals based on suppression events extracted from shuffled multiunit activity.

**Figure S7.**
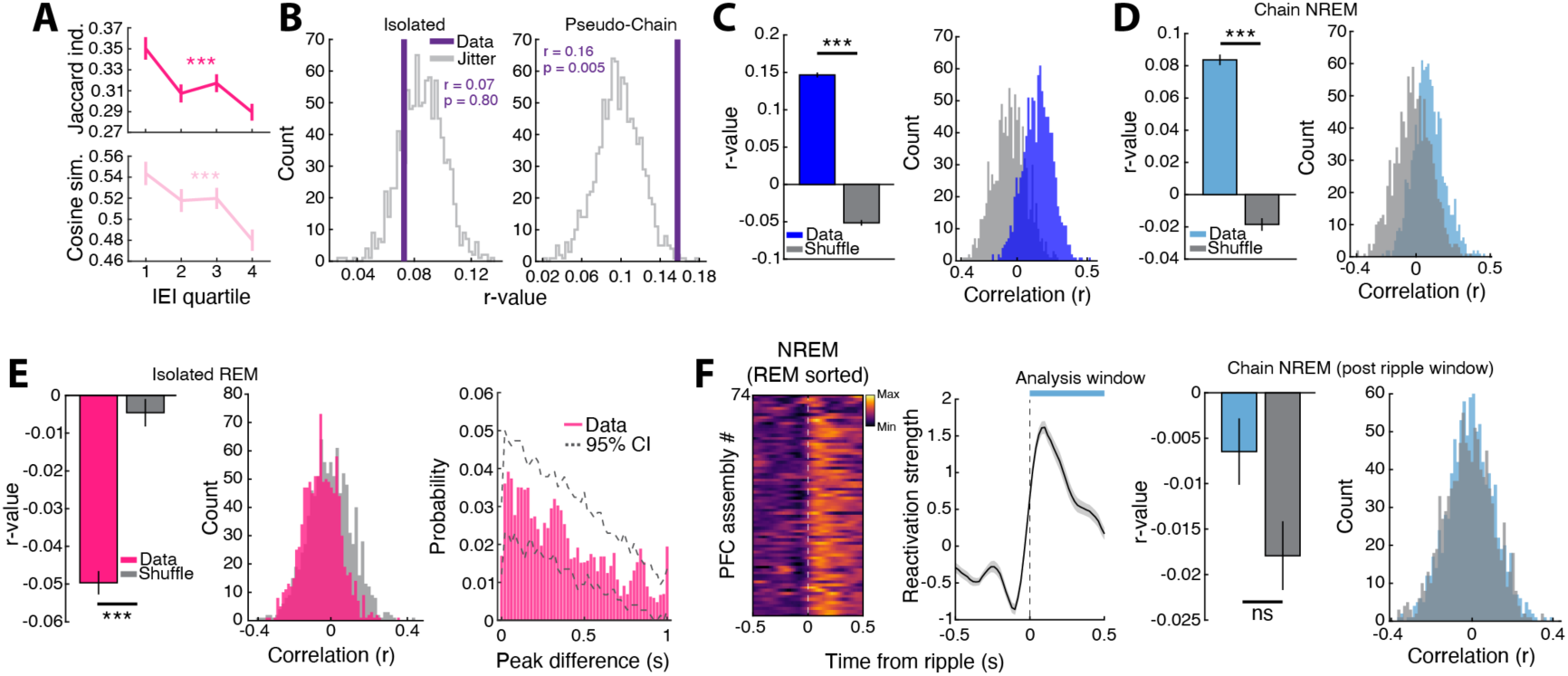
Related to Figures 5 and 6. REM PFC HFOs are associated with sequential reactivation as compared to NREM PFC ripples. (**A**) Relationship between IRI and (Top) Jaccard index and (Bottom) cosine similarity (Jaccard index, r = -0.094, ***p = 1.30×10^-4^; Cosine similarity, r = -0.097, ***p = 7.16×10^-5^, Pearson correlation). (**B**) Rank order correlation analysis on isolated REM PFC HFOs (Left) and pseudo-chain events (Right). (**C**) Cross-validation of the REM reactivation sequences surrounding gamma events (Data = 0.147 ± 0.003, Shuffle = -0.051 ± 0.004, ***p = 6.45×10^-208^, WRS). (**D**) (Left) Same as in (**C**) but for isolated REM HFOs (Data = 0.050 ± 0.003, Shuffle = - 0.005 ± 0.004, ***p = 4.82×10^-4^, WRS). (Right) Probability of assembly peak displacement between randomly chosen halves of the isolated REM HFO aligned data. Note the absence of peaks at small displacements as seen in **Figure 6E**. (**E**) Same as in (**C**) but for NREM ripple chains (Data = 0.084 ± 0.003, Shuffle = -0.019 ± 0.004, ***p = 1.37×10^-75^, WRS). (**F**) Cross-validation of the NREM reactivation sequences in the post ripple window (+500 ms from first chained ripple onset) (Data = -0.007 ± 0.004, Shuffle = -0.018 ± 0.004, p = 0.062, WRS).

**Figure S8.**
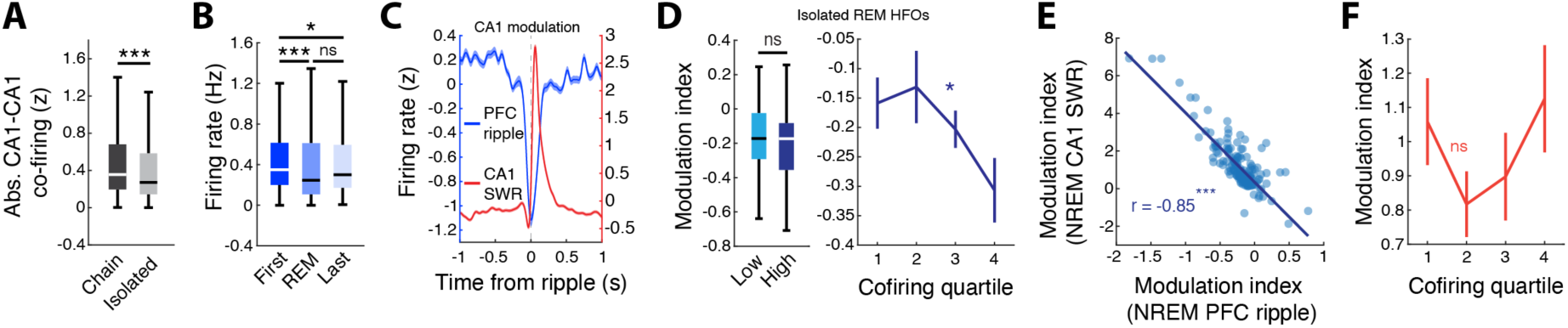
Related to Figure 7. CA1 suppression during NREM PFC ripples and CA1 REM HFO cofiring. (**A**) Absolute CA1-CA1 cell pair cofiring during chained and isolated HFOs (Chain = 0.53 ± 0.02, Isolated = 0.48 ± 0.02, ***p = 1.42×10^-4^, WRS). The magnitude of cofiring (positive and negative) was greater for HFO chains as compared to isolated HFOs, suggesting differential engagement of CA1 neurons. (**B**) Firing rates of CA1 pyramidal neurons during the first 30 s of NREM, REM, and the last 30 s of NREM sleep within an epoch (First 30 s NREM = 0.54 ± 0.04 Hz, REM = 0.51 ± 0.04 Hz, Last 30 s NREM = 0.51 ± 0.03, First vs REM, ***p = 1.10×10^-4^, REM vs Last, p = 0.61, First vs Last, *p = 0.013, Friedman test with Bonferroni correction). CA1 firing rates decreased during REM and later in NREM. (**C**) During NREM sleep, CA1 neurons were strongly suppressed during independent PFC ripples. (**D**) (Left) There was no difference in the degree of suppression during NREM PFC ripples for low and high cofiring (during isolated REM HFOs) CA1 neurons (Low cofiring = -0.16 ± 0.04, High cofiring = -0.21 ± 0.03, p = 0.40, WRS). (Right) The cofiring magnitude during isolated HFOs was correlated with the degree of suppression during NREM PFC ripples, similar to REM HFO chains (r = -0.19, *p = 0.011, Pearson correlation). (**E**) Correlation between CA1 modulation during independent NREM PFC ripples and coordinated SWRs. The more suppressed CA1 neurons are during PFC ripples, the stronger they are reactivated during coordinated SWRs (r = -0.85, ***p = 6.45×10^-208^, Pearson correlation). (**F**) There was no relationship between CA1 cofiring (with PFC) during REM HFO chains and the degree of modulation during coordinated SWRs in NREM sleep (r = 0.04, p = 0.44, Pearson correlation).

**Figure S9.**
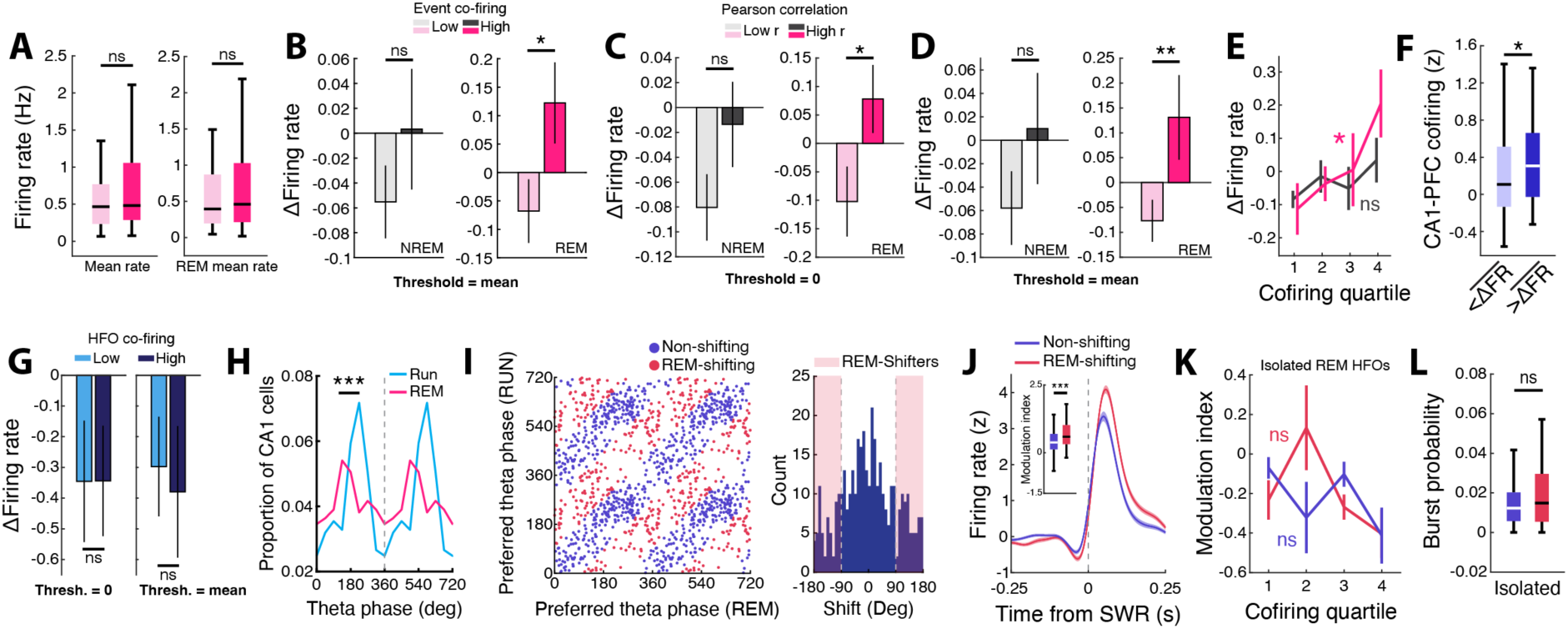
Related to Figures 7 and 8. Relationship between REM PFC HFO cofiring, changes in CA1 firing rates over the course of sleep, and CA1 theta phase shifting neurons. (**A**) (Left) Mean firing rates of high and low cofiring CA1 pyramidal neurons across the entire sleep epoch or (Right) restricted to REM sleep (Low cofiring mean rate = 0.69 ± 0.10, High cofiring mean rate = 0.79 ± 0.08, p = 0.41; Low cofiring REM mean rate = 0.73 ± 0.12, High cofiring REM mean rate = 0.78 ± 0.10, p = 0.49, WRS). (**B**) (Left) Changes in CA1 firing rates from the first NREM bout to the last NREM bout within a sleep epoch plotted according to degree of cofiring (cofiring as measured by the z-scored cofiring metric; high, z-cofiring > mean of distribution; low, z-cofiring < mean of distribution) with PFC either during CA1-independent NREM ripples or chained REM HFOs in PFC (Low NREM = -0.06 ± 0.03, High NREM = 0.003 ± 0.048, p = 0.40; Low REM = -0.068 ± 0.056, High REM = 0.122 ± 0.071, *p = 0.013, WRS). (**C**) Changes in CA1 firing rates from the first NREM bout to the last NREM bout within a sleep epoch plotted according to degree of cofiring (cofiring as measured by the Pearson correlation coefficient; high, r > 0; low, r < 0) with PFC either during CA1-independent NREM ripples or chained REM HFOs in PFC (Low NREM = -0.08 ± 0.03, High NREM = -0.01 ± 0.03, p = 0.080; Low REM = -0.10 ± 0.06, High REM = 0.08 ± 0.06, *p = 0.012, WRS). (**D**) Same as in (**E**) but using above or below the mean of the cofiring distribution to designate high or low cofiring CA1 units, respectively (Low NREM = -0.06 ± 0.03, High NREM = 0.01 ± 0.05, p = 0.30; Low REM = -0.08 ± 0.04, High REM = 0.13 ± 0.08, **p = 0.002, WRS). (**E**) Firing rate change separated by cofiring quartile. The change in CA1 firing rates over the course of sleep was correlated with PFC cofiring during REM HFOs but not NREM ripples (NREM, r = 0.13, p = 0.49; REM, r = 0.17, *p = 0.029, Robust linear regression). (**F**) CA1-PFC HFO cofiring compared for CA1 neurons with firing rate changes higher or lower than the mean (-0.04 Hz) of the distribution (Higher = 0.34 ± 0.04, Lower = 0.22 ± 0.06, *p = 0.038, WRS). Similar to (**Figure 7F**), CA1 neurons with positive shifts in firing rate over a sleep epoch had higher PFC cofiring than CA1 neurons with a decrease in rate. (**G**) Changes in PFC firing rates from the first NREM bout to the last NREM bout within a sleep epoch plotted according to degree of cofiring with CA1 during REM HFO chains in PFC. Results using 0 or the mean of the correlation coefficient distribution as the threshold shown on the left and right, respectively (0 threshold: Low = -0.35 ± 0.20, High = -0.34 ± 0.18, p = 0.90; Mean threshold: Low = -0.30 ± 0.21, High = -0.38 ± 0.21, p = 0.98, WRS). (**H**) Preferred CA1 theta phase of CA1 neurons during running behavior or REM sleep. A subset of CA1 cells shifted their preferred theta phase during REM sleep (U^2^ = 0.48, ***p = 1.51×10^-5^, Watson’s U^2^ test). Note that only cells that were significantly phase locked to the theta oscillation during both run and REM were used for analysis. (**I**) (Left) Distributions of preferred CA1 theta phases during run and REM sleep. Only CA1 neurons with significant phase locking during both run and REM sleep were included for analysis. (Right) Histogram of theta phase shifts between run and the subsequent REM sleep session. Red shaded areas indicate neurons that had a phase shift >90°. (**J**) (Left) SWR aligned firing rates of non-shifting and REM-shifting CA1 neurons. (Right) REM-shifting neurons had higher modulation indices, indicating stronger excitation during SWRs (Non-shifting = 0.60 ± 0.07, REM-shifting = 0.80 ± 0.07, ***p = 8.51×10^-5^, WRS) (**K**) The cofiring magnitude during isolated REM HFOs of non-shifting or REM-shifting CA1 neurons was not correlated with the degree of suppression during independent NREM PFC ripples (Non-shifting, r = -0.25, p = 0.085, REM-shifting, r = -0.32, p = 0.13, Pearson correlation). (**L**) Burst probability of non-shifting and REM-shifting CA1 neurons during isolated REM H (Non-shifting = 0.015 ± 0.002, REM-shifting = 0.021 ± 0.003, p = 0.22, WRS).

**Figure S10.**
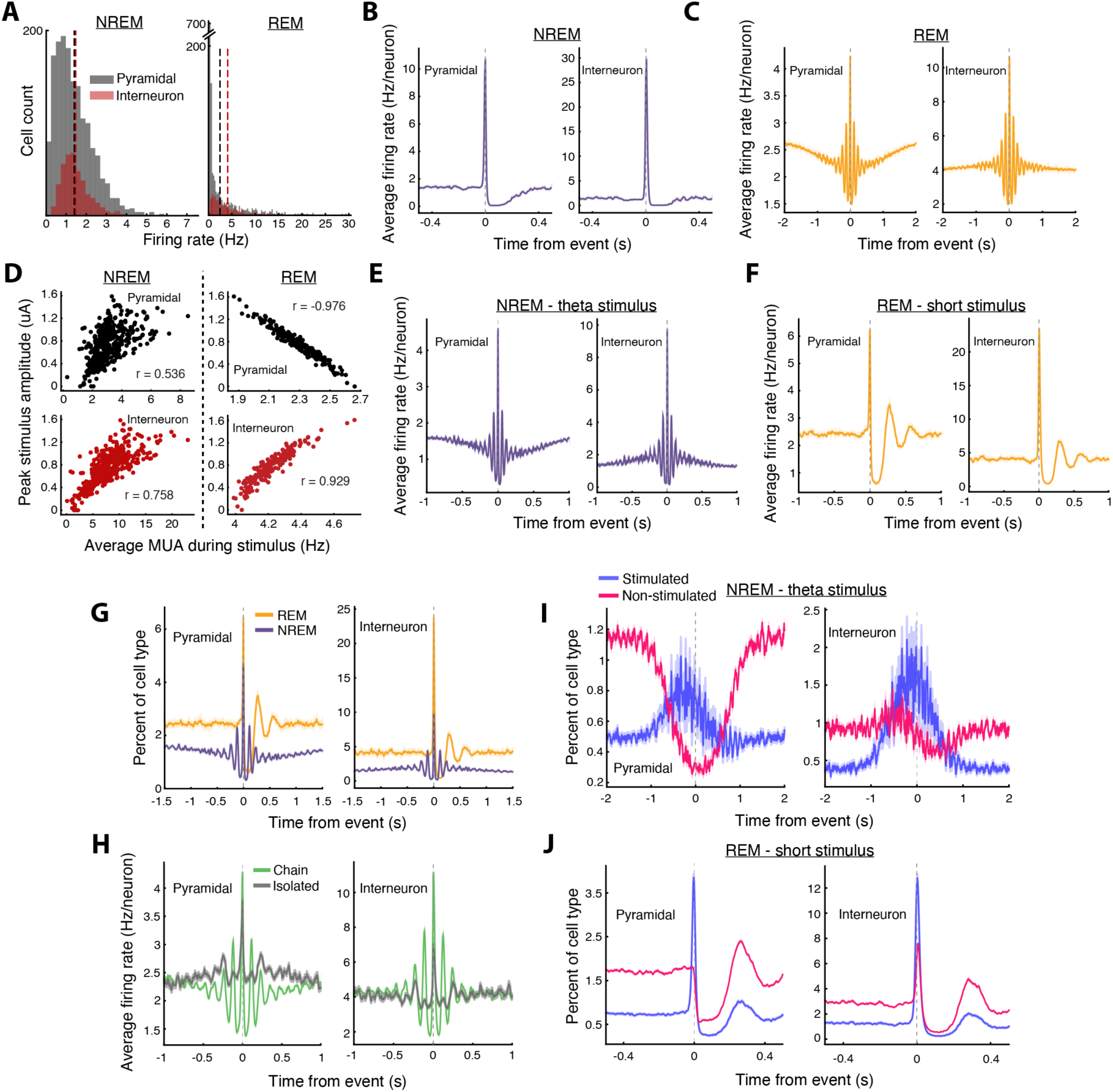
Related to Figures 8 and 9. Model analyses split by cell type and reversed condition controls demonstrating the specific effects of ACh tone on network response. (**A**) Firing rate histograms for NREM and REM simulations split by cell type. Bin size 0.25 Hz. NREM mean rate 1.41 Hz for pyramidal, 1.46 Hz for interneurons. REM mean rate 2.44 Hz for pyramidal, 4.14 Hz for interneurons. (**B**) Average network firing rate surrounding NREM MUA peaks in stimulus periods, split by cell type. (**C**) Same as (**B**), but for REM. (**D**) Peak stimulus amplitude vs average MUA within the stimulus period split by cell type for NREM (Left) and REM (Right). (**E**) Reversed stimulus condition for NREM. Same as (**B**), but swapping the stimulus profile by applying the REM theta-modulated stimulus profile during a low ACh (NREM) simulation. All other parameters are identical. (**F**) Reversed stimulus condition for REM. Same as (**C**), but for a simulation using the brief NREM stimulus combined with high ACh (REM). (**G**) Percent of each cell type with at least one spike in 10 ms bins for the swapped stimulus profile / ACh condition. Note the reduced breadth of pyramidal network activity when applying the brief stimulus in REM compared to NREM, compared to **Figure 10B**. (**H**) Same as **Figure 9G**, but split by cell type. (**I**) Swapped stimulus profile / ACh condition, applying the REM theta stimulus during a low ACh NREM simulation. Note the widespread recruitment of stimulated cells relative to non-stimulated cells, compared to REM response shown in **Figure 10D**. (**J**) Same as (**I**), but applying the NREM brief stimulus during a high ACh REM simulation. Note the lack of peak in non-stimulated cell recruitment by the stimulus compared to NREM response shown in **Figure 10C**.

**Figure S11.**
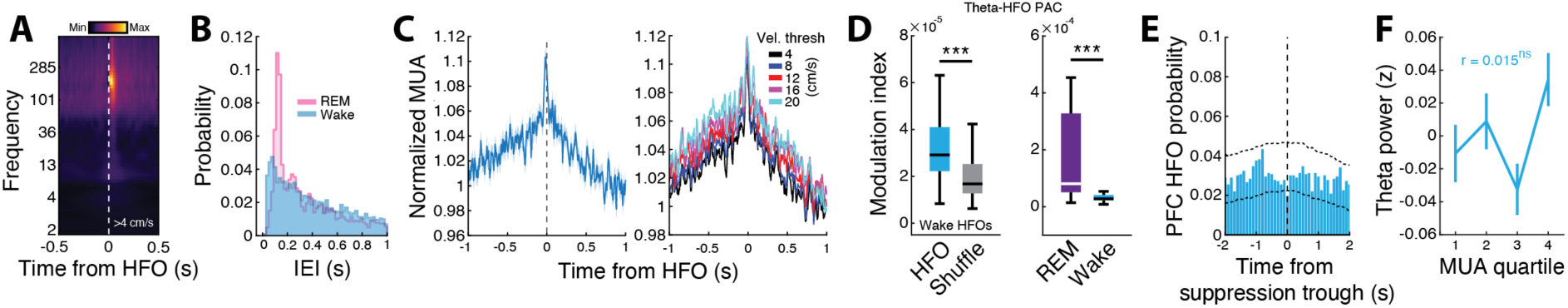
Related to Figures 1 through 4 and 9. HFOs during awake behavior on the W-Track. (**B**) Animal averaged wake HFO aligned PFC spectrogram. Note the absence of elevated gamma and theta power compared to REM events. Compare to **Figure 1F**. (**C**) Distributions of inter-event-intervals (IEIs) for Wake and REM HFOs (***p = 1.13×10^-17^, Kolmogorov-Smirnov test) (**D**) (Left) Wake HFO aligned PFC multiunit activity. (Right) Wake HFO aligned multiunit activity stratified by velocity at HFO occurrence. Compare to **Figure 2A**. (**E**) (Left) Wake theta-HFO phase amplitude coupling compared to shuffle. Note the y-axis scale compared to **Figure S4A** (HFO = 3.81×10^-5^ ± 3.63×10^-6^, Shuffle = 1.83×10^-5^ ± 1.24×10^-6^, ***p = 3.16×10^-12^, WRS). (Right) Comparison of theta-HFO phase amplitude coupling between wake and REM sleep (REM = 1.30×10^-3^ ± 4.54×10^-4^, Wake = 3.65×10^-5^ ± 4.03×10^-6^, ***p = 5.21×10^-5^, WRS). Here, REM modulation index values are lower than in **Figure S4A** because, in this case, all of REM sleep was evaluated, whereas only putative phasic bouts were analyzed in **Figure S4A**. (**F**) The probability of wake HFOs centered on population suppression events detected during wake behavior. Compare to **Figure 4H**. The black dotted lines indicate the 99% confidence intervals based on suppression events extracted from shuffled multiunit data. (**G**) There was no correlation between theta power during HFOs and multiunit activity. Compare to **Figure 9E** (r = 0.015, p = 0.063, Pearson correlation).

## Notes

### Competing Interest Statement

The authors have declared no competing interest.

### Summary of Updates

Addition of Figure 1C, Figure 3A, Figures 6F-I. Addition of Supplemental Figures S1B,H-I, Figures S3C,F-G, Figure S4H, Figures S5L-N, Figure S11. Figure-associated text added and other text revisions.

## REFERENCES

1. Blumberg, M.S., Lesku, J.A., Libourel, P.A., Schmidt, M.H., and Rattenborg, N.C. (2020). What Is REM Sleep? Curr Biol 30, R38–R49. 10.1016/j.cub.2019.11.045.

2. Peever, J., and Fuller, P.M. (2017). The Biology of REM Sleep. Curr Biol 27, R1237–R1248. 10.1016/j.cub.2017.10.026.

3. Tempesta, D., Socci, V., De Gennaro, L., and Ferrara, M. (2018). Sleep and emotional processing. Sleep Med Rev 40, 183–195. 10.1016/j.smrv.2017.12.005.

4. Mukai, Y., and Yamanaka, A. (2023). Functional roles of REM sleep. Neurosci Res 189, 44–53. 10.1016/j.neures.2022.12.009.

5. Lendner, J.D., Niethard, N., Mander, B.A., van Schalkwijk, F.J., Schuh-Hofer, S., Schmidt, H., Knight, R.T., Born, J., Walker, M.P., Lin, J.J., and Helfrich, R.F. (2023). Human REM sleep recalibrates neural activity in support of memory formation. Sci Adv 9, eadj1895. 10.1126/sciadv.adj1895.

6. Dauvilliers, Y., Schenck, C.H., Postuma, R.B., Iranzo, A., Luppi, P.H., Plazzi, G., Montplaisir, J., and Boeve, B. (2018). REM sleep behaviour disorder. Nat Rev Dis Primers 4, 19. 10.1038/s41572-018-0016-5.

7. Squire, L.R., and Alvarez, P. (1995). Retrograde amnesia and memory consolidation: a neurobiological perspective. Curr Opin Neurobiol 5, 169–177. 10.1016/0959-4388(95)80023-9.

8. Nadel, L., and Moscovitch, M. (1997). Memory consolidation, retrograde amnesia and the hippocampal complex. Curr Opin Neurobiol 7, 217–227. 10.1016/s0959-4388(97)80010-4.

9. Buzsaki, G. (1989). Two-stage model of memory trace formation: a role for "noisy" brain states. Neuroscience 31, 551–570. 10.1016/0306-4522(89)90423-5.

10. Sekeres, M.J., Winocur, G., and Moscovitch, M. (2018). The hippocampus and related neocortical structures in memory transformation. Neurosci Lett 680, 39–53. 10.1016/j.neulet.2018.05.006.

11. Staresina, B.P. (2024). Coupled sleep rhythms for memory consolidation. Trends Cogn Sci 28, 339–351. 10.1016/j.tics.2024.02.002.

12. Buzsaki, G. (2015). Hippocampal sharp wave-ripple: A cognitive biomarker for episodic memory and planning. Hippocampus 25, 1073–1188. 10.1002/hipo.22488.

13. Girardeau, G., and Zugaro, M. (2011). Hippocampal ripples and memory consolidation. Curr Opin Neurobiol 21, 452–459. 10.1016/j.conb.2011.02.005.

14. Klinzing, J.G., Niethard, N., and Born, J. (2019). Mechanisms of systems memory consolidation during sleep. Nat Neurosci 22, 1598–1610. 10.1038/s41593-019-0467-3.

15. Mizuseki, K., Diba, K., Pastalkova, E., and Buzsaki, G. (2011). Hippocampal CA1 pyramidal cells form functionally distinct sublayers. Nat Neurosci 14, 1174–1181. 10.1038/nn.2894.

16. Tang, W., Shin, J.D., Frank, L.M., and Jadhav, S.P. (2017). Hippocampal-Prefrontal Reactivation during Learning Is Stronger in Awake Compared with Sleep States. J Neurosci 37, 11789–11805. 10.1523/JNEUROSCI.2291-17.2017.

17. Boyce, R., Glasgow, S.D., Williams, S., and Adamantidis, A. (2016). Causal evidence for the role of REM sleep theta rhythm in contextual memory consolidation. Science 352, 812–816. 10.1126/science.aad5252.

18. Montgomery, S.M., Sirota, A., and Buzsaki, G. (2008). Theta and gamma coordination of hippocampal networks during waking and rapid eye movement sleep. J Neurosci 28, 6731–6741. 10.1523/JNEUROSCI.1227-08.2008.

19. Brankack, J., Scheffzuk, C., Kukushka, V.I., Vyssotski, A.L., Tort, A.B., and Draguhn, A. (2012). Distinct features of fast oscillations in phasic and tonic rapid eye movement sleep. J Sleep Res 21, 630–633. 10.1111/j.1365-2869.2012.01037.x.

20. Bandarabadi, M., Boyce, R., Gutierrez Herrera, C., Bassetti, C.L., Williams, S., Schindler, K., and Adamantidis, A. (2019). Dynamic modulation of theta-gamma coupling during rapid eye movement sleep. Sleep 42. 10.1093/sleep/zsz182.

21. Bueno-Junior, L.S., Ruckstuhl, M.S., Lim, M.M., and Watson, B.O. (2023). The temporal structure of REM sleep shows minute-scale fluctuations across brain and body in mice and humans. Proc Natl Acad Sci U S A 120, e2213438120. 10.1073/pnas.2213438120.

22. Tort, A.B., Scheffer-Teixeira, R., Souza, B.C., Draguhn, A., and Brankack, J. (2013). Theta-associated high-frequency oscillations (110-160Hz) in the hippocampus and neocortex. Prog Neurobiol 100, 1–14. 10.1016/j.pneurobio.2012.09.002.

23. Ghosh, M., Yang, F.C., Rice, S.P., Hetrick, V., Gonzalez, A.L., Siu, D., Brennan, E.K.W., John, T.T., Ahrens, A.M., and Ahmed, O.J. (2022). Running speed and REM sleep control two distinct modes of rapid interhemispheric communication. Cell Rep 40, 111028. 10.1016/j.celrep.2022.111028.

24. Arndt, K.C., Gilbert, E.T., Klaver, L.M.F., Kim, J., Buhler, C.M., Basso, J.C., McKenzie, S., and English, D.F. (2024). Granular retrosplenial cortex layer 2/3 generates high-frequency oscillations dynamically coupled with hippocampal rhythms across brain states. Cell Rep 43, 113910. 10.1016/j.celrep.2024.113910.

25. Aleman-Zapata, A., Morris, R.G.M., and Genzel, L. (2022). Sleep deprivation and hippocampal ripple disruption after one-session learning eliminate memory expression the next day. Proc Natl Acad Sci U S A 119, e2123424119. 10.1073/pnas.2123424119.

26. Yamada, R.G., and Ueda, H.R. (2019). Molecular Mechanisms of REM Sleep. Front Neurosci 13, 1402. 10.3389/fnins.2019.01402.

27. Khodagholy, D., Gelinas, J.N., and Buzsaki, G. (2017). Learning-enhanced coupling between ripple oscillations in association cortices and hippocampus. Science 358, 369–372. 10.1126/science.aan6203.

28. Shin, J.D., and Jadhav, S.P. (2024). Prefrontal cortical ripples mediate top-down suppression of hippocampal reactivation during sleep memory consolidation. Curr Biol 34, 2801–2811 e2809. 10.1016/j.cub.2024.05.018.

29. Vaz, A.P., Inati, S.K., Brunel, N., and Zaghloul, K.A. (2019). Coupled ripple oscillations between the medial temporal lobe and neocortex retrieve human memory. Science 363, 975–978. 10.1126/science.aau8956.

30. van Schalkwijk, F.J., Weber, J., Hahn, M.A., Lendner, J.D., Inostroza, M., Lin, J.J., and Helfrich, R.F. (2023). An evolutionary conserved division-of-labor between archicortical and neocortical ripples organizes information transfer during sleep. Prog Neurobiol 227, 102485. 10.1016/j.pneurobio.2023.102485.

31. Scheffzuk, C., Kukushka, V.I., Vyssotski, A.L., Draguhn, A., Tort, A.B., and Brankack, J. (2011). Selective coupling between theta phase and neocortical fast gamma oscillations during REM-sleep in mice. PLoS One 6, e28489. 10.1371/journal.pone.0028489.

32. van Schalkwijk, F.J., and Helfrich, R.F. (2026). Aperiodic 1/f noise drives ripple activity in humans. Nat Commun 17, 746. 10.1038/s41467-026-68404-5.

33. Grosmark, A.D., Mizuseki, K., Pastalkova, E., Diba, K., and Buzsaki, G. (2012). REM sleep reorganizes hippocampal excitability. Neuron 75, 1001–1007. 10.1016/j.neuron.2012.08.015.

34. Miyawaki, H., and Diba, K. (2016). Regulation of Hippocampal Firing by Network Oscillations during Sleep. Curr Biol 26, 893–902. 10.1016/j.cub.2016.02.024.

35. Watson, B.O., Levenstein, D., Greene, J.P., Gelinas, J.N., and Buzsaki, G. (2016). Network Homeostasis and State Dynamics of Neocortical Sleep. Neuron 90, 839–852. 10.1016/j.neuron.2016.03.036.

36. Poe, G.R., Nitz, D.A., McNaughton, B.L., and Barnes, C.A. (2000). Experience-dependent phase-reversal of hippocampal neuron firing during REM sleep. Brain Res 855, 176–180. 10.1016/s0006-8993(99)02310-0.

37. Louie, K., and Wilson, M.A. (2001). Temporally structured replay of awake hippocampal ensemble activity during rapid eye movement sleep. Neuron 29, 145–156. 10.1016/s0896-6273(01)00186-6.

38. Abdou, K., Nomoto, M., Aly, M.H., Ibrahim, A.Z., Choko, K., Okubo-Suzuki, R., Muramatsu, S.I., and Inokuchi, K. (2024). Prefrontal coding of learned and inferred knowledge during REM and NREM sleep. Nat Commun 15, 4566. 10.1038/s41467-024-48816-x.

39. Maharjan, D.M., Dai, Y.Y., Glantz, E.H., and Jadhav, S.P. (2018). Disruption of dorsal hippocampal - prefrontal interactions using chemogenetic inactivation impairs spatial learning. Neurobiol Learn Mem 155, 351–360. 10.1016/j.nlm.2018.08.023.

40. Shin, J.D., Tang, W., and Jadhav, S.P. (2019). Dynamics of Awake Hippocampal-Prefrontal Replay for Spatial Learning and Memory-Guided Decision Making. Neuron 104, 1110–1125 e1117. 10.1016/j.neuron.2019.09.012.

41. Staresina, B.P., Niediek, J., Borger, V., Surges, R., and Mormann, F. (2023). How coupled slow oscillations, spindles and ripples coordinate neuronal processing and communication during human sleep. Nat Neurosci 26, 1429–1437. 10.1038/s41593-023-01381-w.

42. Canolty, R.T., and Knight, R.T. (2010). The functional role of cross-frequency coupling. Trends Cogn Sci 14, 506–515. 10.1016/j.tics.2010.09.001.

43. de Almeida-Filho, D.G., Koike, B.D.V., Billwiller, F., Farias, K.S., de Sales, I.R.P., Luppi, P.H., Ribeiro, S., and Queiroz, C.M. (2021). Hippocampus-retrosplenial cortex interaction is increased during phasic REM and contributes to memory consolidation. Sci Rep 11, 13078. 10.1038/s41598-021-91659-5.

44. Davidson, T.J., Kloosterman, F., and Wilson, M.A. (2009). Hippocampal replay of extended experience. Neuron 63, 497–507. 10.1016/j.neuron.2009.07.027.

45. Darevsky, D., Kim, J., and Ganguly, K. (2024). Coupling of Slow Oscillations in the Prefrontal and Motor Cortex Predicts Onset of Spindle Trains and Persistent Memory Reactivations. J Neurosci 44. 10.1523/JNEUROSCI.0621-24.2024.

46. Antony, J.W., Piloto, L., Wang, M., Pacheco, P., Norman, K.A., and Paller, K.A. (2018). Sleep Spindle Refractoriness Segregates Periods of Memory Reactivation. Curr Biol 28, 1736–1743 e1734. 10.1016/j.cub.2018.04.020.

47. Mallory, C.S., Widloski, J., and Foster, D.J. (2025). The time course and organization of hippocampal replay. Science 387, 541–548. 10.1126/science.ads4760.

48. Cheng, S., and Frank, L.M. (2008). New experiences enhance coordinated neural activity in the hippocampus. Neuron 57, 303–313. 10.1016/j.neuron.2007.11.035.

49. Judak, L., Chiovini, B., Juhasz, G., Palfi, D., Mezriczky, Z., Szadai, Z., Katona, G., Szmola, B., Ocsai, K., Martinecz, B., et al. (2022). Sharp-wave ripple doublets induce complex dendritic spikes in parvalbumin interneurons in vivo. Nat Commun 13, 6715. 10.1038/s41467-022-34520-1.

50. Peyrache, A., Khamassi, M., Benchenane, K., Wiener, S.I., and Battaglia, F.P. (2009). Replay of rule-learning related neural patterns in the prefrontal cortex during sleep. Nat Neurosci 12, 919–926. 10.1038/nn.2337.

51. van de Ven, G.M., Trouche, S., McNamara, C.G., Allen, K., and Dupret, D. (2016). Hippocampal Offline Reactivation Consolidates Recently Formed Cell Assembly Patterns during Sharp Wave-Ripples. Neuron 92, 968–974. 10.1016/j.neuron.2016.10.020.

52. Fries, P. (2005). A mechanism for cognitive dynamics: neuronal communication through neuronal coherence. Trends Cogn Sci 9, 474–480. 10.1016/j.tics.2005.08.011.

53. Miyawaki, H., Watson, B.O., and Diba, K. (2019). Neuronal firing rates diverge during REM and homogenize during non-REM. Sci Rep 9, 689. 10.1038/s41598-018-36710-8.

54. Valero, M., Cid, E., Averkin, R.G., Aguilar, J., Sanchez-Aguilera, A., Viney, T.J., Gomez-Dominguez, D., Bellistri, E., and de la Prida, L.M. (2015). Determinants of different deep and superficial CA1 pyramidal cell dynamics during sharp-wave ripples. Nat Neurosci 18, 1281–1290. 10.1038/nn.4074.

55. Gu, L., Ren, M., Lin, L., and Xu, J. (2023). Calbindin-Expressing CA1 Pyramidal Neurons Encode Spatial Information More Efficiently. eNeuro 10. 10.1523/ENEURO.0411-22.2023.

56. de la Prida, L.M. (2020). Potential factors influencing replay across CA1 during sharp-wave ripples. Philos Trans R Soc Lond B Biol Sci 375, 20190236. 10.1098/rstb.2019.0236.

57. Harvey, R.E., Robinson, H.L., Liu, C., Oliva, A., and Fernandez-Ruiz, A. (2023). Hippocampo-cortical circuits for selective memory encoding, routing, and replay. Neuron 111, 2076–2090 e2079. 10.1016/j.neuron.2023.04.015.

58. Sharif, F., Tayebi, B., Buzsaki, G., Royer, S., and Fernandez-Ruiz, A. (2021). Subcircuits of Deep and Superficial CA1 Place Cells Support Efficient Spatial Coding across Heterogeneous Environments. Neuron 109, 363–376 e366. 10.1016/j.neuron.2020.10.034.

59. Picciotto, M.R., Higley, M.J., and Mineur, Y.S. (2012). Acetylcholine as a neuromodulator: cholinergic signaling shapes nervous system function and behavior. Neuron 76, 116–129. 10.1016/j.neuron.2012.08.036.

60. Eban-Rothschild, A., and de Lecea, L. (2017). Neuronal substrates for initiation, maintenance, and structural organization of sleep/wake states. F1000Res 6, 212. 10.12688/f1000research.9677.1.

61. Colangelo, C., Shichkova, P., Keller, D., Markram, H., and Ramaswamy, S. (2019). Cellular, Synaptic and Network Effects of Acetylcholine in the Neocortex. Front Neural Circuits 13, 24. 10.3389/fncir.2019.00024.

62. Stiefel, K.M., Gutkin, B.S., and Sejnowski, T.J. (2009). The effects of cholinergic neuromodulation on neuronal phase-response curves of modeled cortical neurons. J Comput Neurosci 26, 289–301. 10.1007/s10827-008-0111-9.

63. Roach, J.P., Ben-Jacob, E., Sander, L.M., and Zochowski, M.R. (2015). Formation and Dynamics of Waves in a Cortical Model of Cholinergic Modulation. PLoS Comput Biol 11, e1004449. 10.1371/journal.pcbi.1004449.

64. Satchell, M., Butel-Fry, E., Noureddine, Z., Simmons, A., Ognjanovski, N., Aton, S.J., and Zochowski, M.R. (2025). Cholinergic modulation of neural networks supports sequential and complementary roles for NREM and REM states in memory consolidation. PLoS Comput Biol 21, e1013097. 10.1371/journal.pcbi.1013097.

65. Czarnecki, P., Lin, J., Aton, S.J., and Zochowski, M. (2021). Dynamical Mechanism Underlying Scale-Free Network Reorganization in Low Acetylcholine States Corresponding to Slow Wave Sleep. Front Netw Physiol 1. 10.3389/fnetp.2021.759131.

66. Stiefel, K.M., Gutkin, B.S., and Sejnowski, T.J. (2008). Cholinergic neuromodulation changes phase response curve shape and type in cortical pyramidal neurons. PLoS One 3, e3947. 10.1371/journal.pone.0003947.

67. Gulledge, A.T., Bucci, D.J., Zhang, S.S., Matsui, M., and Yeh, H.H. (2009). M1 receptors mediate cholinergic modulation of excitability in neocortical pyramidal neurons. J Neurosci 29, 9888–9902. 10.1523/JNEUROSCI.1366-09.2009.

68. Dasari, S., and Gulledge, A.T. (2011). M1 and M4 receptors modulate hippocampal pyramidal neurons. J Neurophysiol 105, 779–792. 10.1152/jn.00686.2010.

69. Mishra, W., Kheradpezhouh, E., and Arabzadeh, E. (2024). Activation of M1 cholinergic receptors in mouse somatosensory cortex enhances information processing and detection behaviour. Commun Biol 7, 3. 10.1038/s42003-023-05699-w.

70. Kimura, F., Fukuda, M., and Tsumoto, T. (1999). Acetylcholine suppresses the spread of excitation in the visual cortex revealed by optical recording: possible differential effect depending on the source of input. Eur J Neurosci 11, 3597–3609. 10.1046/j.1460-9568.1999.00779.x.

71. Dasgupta, R., Seibt, F., and Beierlein, M. (2018). Synaptic Release of Acetylcholine Rapidly Suppresses Cortical Activity by Recruiting Muscarinic Receptors in Layer 4. J Neurosci 38, 5338–5350. 10.1523/JNEUROSCI.0566-18.2018.

72. Zerimech, S., Chever, O., Scalmani, P., Pizzamiglio, L., Duprat, F., and Mantegazza, M. (2020). Cholinergic modulation inhibits cortical spreading depression in mouse neocortex through activation of muscarinic receptors and decreased excitatory/inhibitory drive. Neuropharmacology 166, 107951. 10.1016/j.neuropharm.2020.107951.

73. Chen, N., Sugihara, H., and Sur, M. (2015). An acetylcholine-activated microcircuit drives temporal dynamics of cortical activity. Nat Neurosci 18, 892–902. 10.1038/nn.4002.

74. Boyce, R., Williams, S., and Adamantidis, A. (2017). REM sleep and memory. Curr Opin Neurobiol 44, 167–177. 10.1016/j.conb.2017.05.001.

75. Bollmann, L., Baracskay, P., Stella, F., and Csicsvari, J. (2025). Sleep stages antagonistically modulate reactivation drift. Neuron. 10.1016/j.neuron.2025.02.025.

76. Helfrich, R.F., Lendner, J.D., Mander, B.A., Guillen, H., Paff, M., Mnatsakanyan, L., Vadera, S., Walker, M.P., Lin, J.J., and Knight, R.T. (2019). Bidirectional prefrontal-hippocampal dynamics organize information transfer during sleep in humans. Nat Commun 10, 3572. 10.1038/s41467-019-11444-x.

77. Drieu, C., and Zugaro, M. (2019). Hippocampal Sequences During Exploration: Mechanisms and Functions. Front Cell Neurosci 13, 232. 10.3389/fncel.2019.00232.

78. Sanzeni, A., Akitake, B., Goldbach, H.C., Leedy, C.E., Brunel, N., and Histed, M.H. (2020). Inhibition stabilization is a widespread property of cortical networks. Elife 9. 10.7554/eLife.54875.

79. Niethard, N., Hasegawa, M., Itokazu, T., Oyanedel, C.N., Born, J., and Sato, T.R. (2016). Sleep-Stage-Specific Regulation of Cortical Excitation and Inhibition. Curr Biol 26, 2739–2749. 10.1016/j.cub.2016.08.035.

80. Ramirez-Villegas, J.F., Besserve, M., Murayama, Y., Evrard, H.C., Oeltermann, A., and Logothetis, N.K. (2021). Coupling of hippocampal theta and ripples with pontogeniculooccipital waves. Nature 589, 96–102. 10.1038/s41586-020-2914-4.

81. Tsunematsu, T., Matsumoto, S., Merkler, M., and Sakata, S. (2023). Pontine Waves Accompanied by Short Hippocampal Sharp Wave-Ripples During Non-rapid Eye Movement Sleep. Sleep 46. 10.1093/sleep/zsad193.

82. Teles-Grilo Ruivo, L.M., Baker, K.L., Conway, M.W., Kinsley, P.J., Gilmour, G., Phillips, K.G., Isaac, J.T.R., Lowry, J.P., and Mellor, J.R. (2017). Coordinated Acetylcholine Release in Prefrontal Cortex and Hippocampus Is Associated with Arousal and Reward on Distinct Timescales. Cell Rep 18, 905–917. 10.1016/j.celrep.2016.12.085.

83. Nunez, A., and Buno, W. (2021). The Theta Rhythm of the Hippocampus: From Neuronal and Circuit Mechanisms to Behavior. Front Cell Neurosci 15, 649262. 10.3389/fncel.2021.649262.

84. Goard, M., and Dan, Y. (2009). Basal forebrain activation enhances cortical coding of natural scenes. Nat Neurosci 12, 1444–1449. 10.1038/nn.2402.

85. Zhang, Y., Cao, L., Varga, V., Jing, M., Karadas, M., Li, Y., and Buzsaki, G. (2021). Cholinergic suppression of hippocampal sharp-wave ripples impairs working memory. Proc Natl Acad Sci U S A 118. 10.1073/pnas.2016432118.

86. Vandecasteele, M., Varga, V., Berenyi, A., Papp, E., Bartho, P., Venance, L., Freund, T.F., and Buzsaki, G. (2014). Optogenetic activation of septal cholinergic neurons suppresses sharp wave ripples and enhances theta oscillations in the hippocampus. Proc Natl Acad Sci U S A 111, 13535–13540. 10.1073/pnas.1411233111.

87. Girardeau, G., Benchenane, K., Wiener, S.I., Buzsaki, G., and Zugaro, M.B. (2009). Selective suppression of hippocampal ripples impairs spatial memory. Nat Neurosci 12, 1222–1223. 10.1038/nn.2384.

88. Maingret, N., Girardeau, G., Todorova, R., Goutierre, M., and Zugaro, M. (2016). Hippocampo-cortical coupling mediates memory consolidation during sleep. Nat Neurosci 19, 959–964. 10.1038/nn.4304.

89. Chang, H., Tang, W., Wulf, A.M., Nyasulu, T., Wolf, M.E., Fernandez-Ruiz, A., and Oliva, A. (2025). Sleep microstructure organizes memory replay. Nature 637, 1161–1169. 10.1038/s41586-024-08340-w.

90. Gridchyn, I., Schoenenberger, P., O’Neill, J., and Csicsvari, J. (2020). Assembly-Specific Disruption of Hippocampal Replay Leads to Selective Memory Deficit. Neuron 106, 291–300 e296. 10.1016/j.neuron.2020.01.021.

91. Porter, B.S., Shi, C., Kozlova, E., and Jadhav, S.P. (In Press). A novel rat spatial transitive inference paradigm reveals rapid deduction of transitive relationships using schemas and deliberation. Behavioral Neuroscience.

92. Zeithamova, D., Schlichting, M.L., and Preston, A.R. (2012). The hippocampus and inferential reasoning: building memories to navigate future decisions. Front Hum Neurosci 6, 70. 10.3389/fnhum.2012.00070.

93. Cairney, S.A., Durrant, S.J., Power, R., and Lewis, P.A. (2015). Complementary roles of slow-wave sleep and rapid eye movement sleep in emotional memory consolidation. Cereb Cortex 25, 1565–1575. 10.1093/cercor/bht349.

94. Simor, P., van der Wijk, G., Nobili, L., and Peigneux, P. (2020). The microstructure of REM sleep: Why phasic and tonic? Sleep Med Rev 52, 101305. 10.1016/j.smrv.2020.101305.

95. Shin, J.D., Tang, W., and Jadhav, S.P. (2023). Protocol for geometric transformation of cognitive maps for generalization across hippocampal-prefrontal circuits. STAR Protoc 4, 102513. 10.1016/j.xpro.2023.102513.

96. Jadhav, S.P., Kemere, C., German, P.W., and Frank, L.M. (2012). Awake hippocampal sharp-wave ripples support spatial memory. Science 336, 1454–1458. 10.1126/science.1217230.

97. Fernandez-Ruiz, A., Oliva, A., Fermino de Oliveira, E., Rocha-Almeida, F., Tingley, D., and Buzsaki, G. (2019). Long-duration hippocampal sharp wave ripples improve memory. Science 364, 1082–1086. 10.1126/science.aax0758.

98. Schmitzer-Torbert, N., Jackson, J., Henze, D., Harris, K., and Redish, A.D. (2005). Quantitative measures of cluster quality for use in extracellular recordings. Neuroscience 131, 1–11. 10.1016/j.neuroscience.2004.09.066.

99. Zhang, L.B., Zhang, J., Sun, M.J., Chen, H., Yan, J., Luo, F.L., Yao, Z.X., Wu, Y.M., and Hu, B. (2020). Neuronal Activity in the Cerebellum During the Sleep-Wakefulness Transition in Mice. Neurosci Bull 36, 919–931. 10.1007/s12264-020-00511-9.

100. Rothschild, G., Eban, E., and Frank, L.M. (2017). A cortical-hippocampal-cortical loop of information processing during memory consolidation. Nat Neurosci 20, 251–259. 10.1038/nn.4457.

101. Schomburg, E.W., Fernandez-Ruiz, A., Mizuseki, K., Berenyi, A., Anastassiou, C.A., Koch, C., and Buzsaki, G. (2014). Theta phase segregation of input-specific gamma patterns in entorhinal-hippocampal networks. Neuron 84, 470–485. 10.1016/j.neuron.2014.08.051.

102. Mitra, P., and Bokil, H. (2008). Observed brain dynamics (Oxford University Press).

103. Sullivan, D., Csicsvari, J., Mizuseki, K., Montgomery, S., Diba, K., and Buzsaki, G. (2011). Relationships between hippocampal sharp waves, ripples, and fast gamma oscillation: influence of dentate and entorhinal cortical activity. J Neurosci 31, 8605–8616. 10.1523/JNEUROSCI.0294-11.2011.

104. Tort, A.B., Komorowski, R., Eichenbaum, H., and Kopell, N. (2010). Measuring phase-amplitude coupling between neuronal oscillations of different frequencies. J Neurophysiol 104, 1195–1210. 10.1152/jn.00106.2010.

105. Vaz, A.P., Yaffe, R.B., Wittig, J.H., Jr., Inati, S.K., and Zaghloul, K.A. (2017). Dual origins of measured phase-amplitude coupling reveal distinct neural mechanisms underlying episodic memory in the human cortex. Neuroimage 148, 148–159. 10.1016/j.neuroimage.2017.01.001.

106. Daume, J., Kaminski, J., Schjetnan, A.G.P., Salimpour, Y., Khan, U., Kyzar, M., Reed, C.M., Anderson, W.S., Valiante, T.A., Mamelak, A.N., and Rutishauser, U. (2024). Control of working memory by phase-amplitude coupling of human hippocampal neurons. Nature 629, 393–401. 10.1038/s41586-024-07309-z.

107. Siapas, A.G., Lubenov, E.V., and Wilson, M.A. (2005). Prefrontal phase locking to hippocampal theta oscillations. Neuron 46, 141–151. 10.1016/j.neuron.2005.02.028.

108. Vinck, M., van Wingerden, M., Womelsdorf, T., Fries, P., and Pennartz, C.M. (2010). The pairwise phase consistency: a bias-free measure of rhythmic neuronal synchronization. Neuroimage 51, 112–122. 10.1016/j.neuroimage.2010.01.073.

109. Nolte, G., Ziehe, A., Nikulin, V.V., Schlogl, A., Kramer, N., Brismar, T., and Muller, K.R. (2008). Robustly estimating the flow direction of information in complex physical systems. Phys Rev Lett 100, 234101. 10.1103/PhysRevLett.100.234101.

110. Oostenveld, R., Fries, P., Maris, E., and Schoffelen, J.M. (2011). FieldTrip: Open source software for advanced analysis of MEG, EEG, and invasive electrophysiological data. Comput Intell Neurosci 2011, 156869. 10.1155/2011/156869.

111. Jadhav, S.P., Rothschild, G., Roumis, D.K., and Frank, L.M. (2016). Coordinated Excitation and Inhibition of Prefrontal Ensembles during Awake Hippocampal Sharp-Wave Ripple Events. Neuron 90, 113–127. 10.1016/j.neuron.2016.02.010.

112. Stark, E., Roux, L., Eichler, R., and Buzsaki, G. (2015). Local generation of multineuronal spike sequences in the hippocampal CA1 region. Proc Natl Acad Sci U S A 112, 10521–10526. 10.1073/pnas.1508785112.

113. Valero, M., Viney, T.J., Machold, R., Mederos, S., Zutshi, I., Schuman, B., Senzai, Y., Rudy, B., and Buzsaki, G. (2021). Sleep down state-active ID2/Nkx2.1 interneurons in the neocortex. Nat Neurosci 24, 401–411. 10.1038/s41593-021-00797-6.

114. Zarzoso, V., and Comon, P. (2010). Robust independent component analysis by iterative maximization of the kurtosis contrast with algebraic optimal step size. IEEE Trans Neural Netw 21, 248–261. 10.1109/TNN.2009.2035920.

115. Sosa, M., Plitt, M.H., and Giocomo, L.M. (2025). A flexible hippocampal population code for experience relative to reward. Nat Neurosci 28, 1497–1509. 10.1038/s41593-025-01985-4.

116. Plitt, M.H., and Giocomo, L.M. (2021). Experience-dependent contextual codes in the hippocampus. Nat Neurosci 24, 705–714. 10.1038/s41593-021-00816-6.

117. Harris, K.D., Hirase, H., Leinekugel, X., Henze, D.A., and Buzsaki, G. (2001). Temporal interaction between single spikes and complex spike bursts in hippocampal pyramidal cells. Neuron 32, 141–149. 10.1016/s0896-6273(01)00447-0.

118. Delmas, P., and Brown, D.A. (2005). Pathways modulating neural KCNQ/M (Kv7) potassium channels. Nat Rev Neurosci 6, 850–862. 10.1038/nrn1785.

119. McCormick, D.A., Wang, Z., and Huguenard, J. (1993). Neurotransmitter control of neocortical neuronal activity and excitability. Cereb Cortex 3, 387–398. 10.1093/cercor/3.5.387.

120. Meir, I., Katz, Y., and Lampl, I. (2018). Membrane Potential Correlates of Network Decorrelation and Improved SNR by Cholinergic Activation in the Somatosensory Cortex. J Neurosci 38, 10692–10708. 10.1523/JNEUROSCI.1159-18.2018.

